# Cystobactamid off-target profiling reveals favorable safety, superoxide reduction, and SCARB1 inhibition in eukaryotes

**DOI:** 10.1101/2024.10.10.617689

**Authors:** Timo Risch, Benedikt Hellwinkel, Dietrich Mostert, Andreas M. Kany, Danny Solga, Tim Seedorf, Dominik Heimann, Jessica Hoppstädter, Daniel Kohnhäuser, Jil-Sophie Hilgers, Franziska Fries, Felix Deschner, Mark Brönstrup, Andreas Kirschning, Stephan A. Sieber, Thomas Pietschmann, Alexandra K. Kiemer, Jennifer Herrmann, Rolf Müller

## Abstract

Antimicrobial resistance (AMR) poses a fundamental global threat, necessitating new strategies for effective therapies. Cystobactamids (CYS), a class of antibacterial agents targeting bacterial gyrase and topoisomerase IV, represent a non-traditional chemical scaffold with broad-spectrum activity. For toxicological de-risking, we performed a comprehensive profiling on eukaryotic cells, focusing on cytotoxicity, genotoxicity, and mitochondrial toxicity, demonstrating cellular safety and superoxide scavenging properties. Studies in zebrafish embryos assessed developmental, cardiovascular, and hepatic toxicity, indicating a favorable *in vivo* safety profile. Metabolism studies revealed glucuronidation and amide bond hydrolysis as key pathways, whereby CYS metabolic stability substantially improved by cobicistat co-treatment. Affinity-based protein profiling identified the cholesterol- and HCV-receptor scavenger receptor class B member 1 (SCARB1) as a primary eukaryotic off-target protein, with cystobactamids shown to inhibit SCARB1’s function, preventing hepatitis C virus pseudoparticle entry into cells. These findings suggest a high therapeutic potential for cystobactamids and highlight SCARB1 as a primary eukaryotic target.

## Introduction

Leading health institutions, including the European medicine agency (EMA) and the World Health Organization (WHO), call attention to antimicrobial resistance (AMR) as an increasing global risk to patients and healthcare systems^1,2,3,4^ The misuse and overuse of antibiotics in humans and also in animals drive the emergence and spread of multidrug-resistant (MDR) bacteria. At the same time, the number of future treatment options using novel antibiotics with innovative structures and mechanisms of action is insufficient.^4,5^ The development of previously identified cystobactamids (CYS) might help to fill this gap, as they represent promising broad-spectrum antibiotics comprising a novel scaffold.^6^

CYS are natural products derived from *Cystobacter and Myxococcus* spp.^7^ They exhibit antibacterial activity against a broad spectrum of clinically relevant Gram-negative and Gram-positive bacteria including MDR isolates of *Acinetobacter baumannii*, *Enterococcus faecalis*, *Streptococcus pneumoniae*, *Staphylococcus aureus* and *Escherichia coli*.^6,7^ CYS act through a new mechanism of action by inhibiting bacterial type IIa topoisomerases (gyrase and topoisomerase IV).^6^ A dual mode of binding for the structurally related albicidin and CYS was recently described with one part of the molecule blocking the gyrase dimer interface and the other end intercalating between cleaved DNA fragments, hence preventing DNA religation.^8^ Importantly, CYS and the structurally related compound classes of albicidin and coralmycin represent a novel chemical scaffold consisting of *para*-nitrobenzoic acid (PNBA) and multiple *para*-aminobenzoic acid (PABA) units connected through an amino acid linker.^9,10^ The novel structure, the lack of cross-resistance with commercially used drugs and their new mechanism of action contribute to the observed resistance-breaking properties within clinical isolates of MDR pathogens.^6,7,11–13^ Furthermore, CYS were shown to have a low frequency of resistance (FoR)^13^, which is of utmost importance for the wide and sustainable use as an antibiotic. Several total syntheses of CYS were established and modified to successfully yield more than hundred derivatives.^7,11,12,14,15^ This allowed for large-scale production and structural modifications to derive structure-activity and structure-property relationships for optimization of potency, antibacterial spectrum coverage, on-target activity, and physicochemical properties.^14,15^

To ensure efficacy and safety, pharmacological and toxicological properties need to be determined before entering clinical studies. In fact, leading causes for failure of drug candidates in the drug development process are lack of efficacy (52%) or an unfavorable safety profile (24%) (2013-2015).^16,17^ Drug approval requires three key properties, namely high quality standards during the production procedure, efficacy against the target disease or symptoms, and a favorable safety profile.^18^ Overall, the benefit of using a certain drug must exceed the risk of serious adverse effects.^19^ Adverse or side effects include innumerable symptoms, which can range from uncritical effects like dizziness or headaches to live-threatening events such as liver damage or arrhythmia.^20,21^

In the presented study, we evaluated the cyto-, geno- and mitotoxicity of CYS derivatives CN-861-2, CN-DM-861 and Cysto-180 using *in vitro* cell culture models, and we investigated more complex organotoxicity with regards to general developmental, cardio- and hepatotoxicity by *in vivo* zebrafish embryo models. Furthermore, metabolic pathways and biotransformation of CYS were investigated *in vitro,* and a strategy to improve their *in vivo* exposure was proposed. Moreover, eukaryotic off-target proteins of CYS were identified and analyzed on a molecular level by affinity-based proteome profiling and functional inhibition assays, surprisingly uncovering yet unexplored potential therapeutic areas for CYS outside their use as antibacterial agents.

## Results and Discussion

### CYS demonstrate general safety in cytotoxicity and genotoxicity assays, with a mild effect on uncoupling the mitochondrial electron transfer chain (ETC)

Three CYS derivatives (CN-861-2, CN-DM-861 and Cysto-180; **Figure 1A**) were chosen to further investigate properties of the class with respect to their biological activity on eukaryotic cells. The selected derivatives comprise early stage frontrunner molecules with improved antibacterial properties (CN-861-2 and CN-DM-861^15^; the former served as main reference compound in this study due to its availability) and a further improved derivative (Cysto-180^13^).

**Figure 1.**
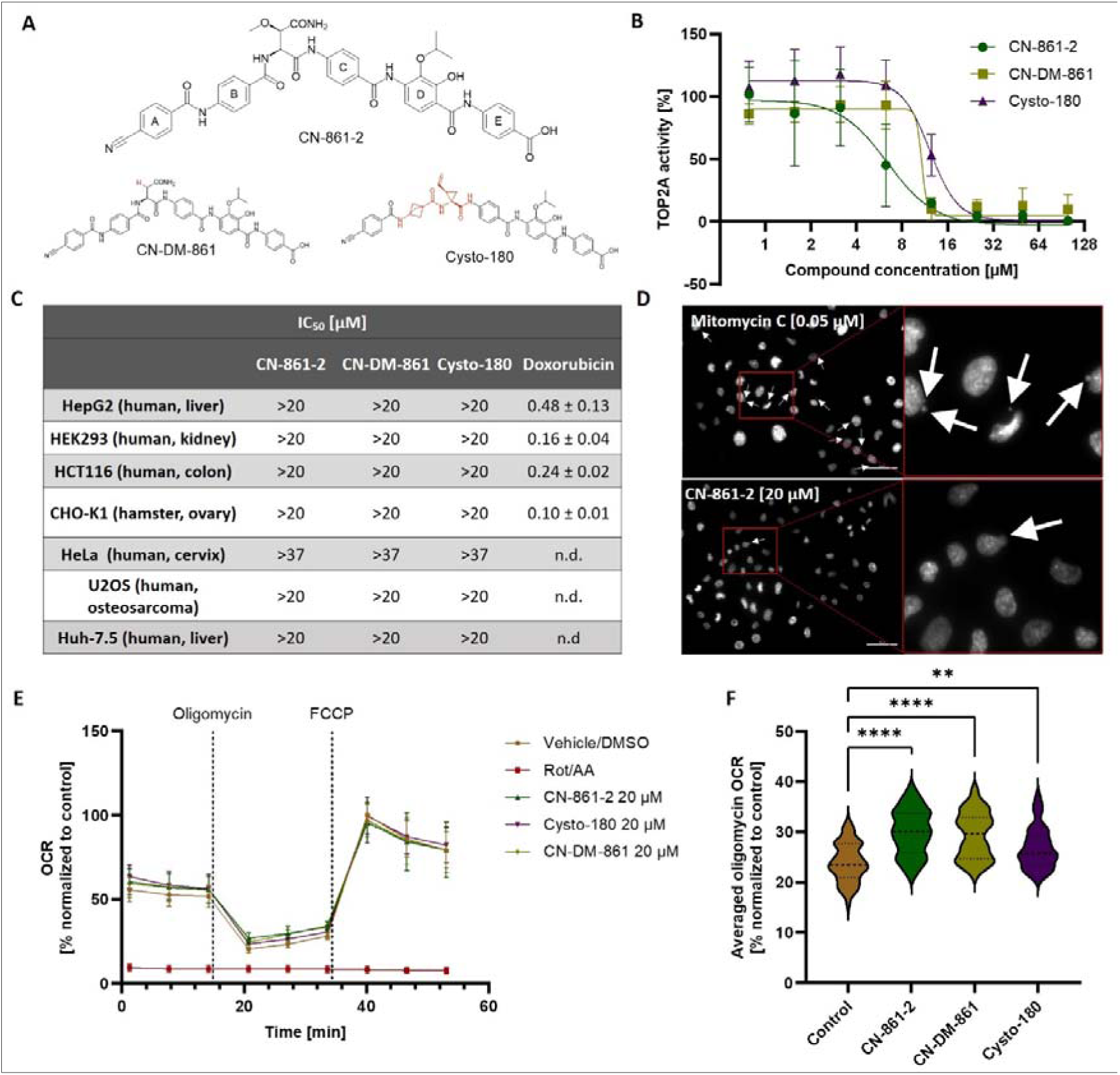
CYS demonstrate general safety in cytotoxicity and genotoxicity assays, with a mild effect on uncoupling the mitochondrial electron transfer chain (ETC). **(A)** Chemical structures of tested CYS derivatives CN-861-2 (top), CN-DM-861 (bottom left) and Cysto-180 (bottom right) in comparison. N-terminal rings A and B are connected via an amino-acid linker with C-terminal rings C to E. Structural differences to CN-861-2 (reference in this study) are marked in red. **(B)** Cystobactamids show concentration dependent inhibition of the human topoisomerase IIα. Data are represented as mean values with standard deviation. IC_50_ was evaluated using non-linear regression (*n* = 3). **(C)** CYS derivatives showed no harming effect on cell viability on human and non-human immortalized cell lines in their solubility range (*n* = 3) (n.d., not determined). **(D)** CYS treated cells showed micronucleus formation comparable to the DMSO control (see also Supplementary Figure 1), while mitomycin C treated cells showed extensive micronucleus formation. (nuclei colored in white, white arrows indicate micronuclei, scale bar = 50 µm) (*n* = 3). (E) Seahorse mitotoxicity assay revealed a slight uncoupling effect of CYS, determined as an increased oxygen consumption rate (OCR) following oligomycin addition. Data are represented as mean values with standard deviation (*n* = 3 x 6 wells). (F) Normalized mean values (% normalized to max OCR of control group) with standard deviations are shown (*n* = 3 x 6). Statistically significant differences of the oligomycin OCR were analyzed by ordinary one-way ANOVA with multiple comparison to the DMSO treated control group. (**, *p* < 0.005; ****, *p* < 0.0001). Violin plot is used for data representation.

Since CYS are bacterial topoisomerase II inhibitors and thus, affect DNA synthesis and repair^22^, an obvious potential molecular off-target of CYS is the human DNA topoisomerase IIα (TOP2A). Half inhibitory concentrations (IC_50_) of 6.26-12.22 µM (**Figure 1B**) against TOP2A were determined (CN-861-2 IC_50_ = 6.26 ± 4.69 µM, CN-DM-861 IC_50_ = 10.83 ± 2.38 µM, Cysto-180 IC_50_ = 12.22 ± 0.63µM), which range far beyond (∼100 fold) previously observed minimal inhibitory concentrations (MIC) of CYS for their target pathogens as well as identified IC_50_ on *E. coli* gyrase (CN-DM-861 IC_50_ = 0.08 µM).^13,15^

Cell culture models are widely used to determine the cytotoxicity of compounds and to initially assess whether molecules interact with fundamental cellular functions affecting cell division and viability.^23^ Interestingly, though their inhibitory effect on the isolated TOP2A activity, CYS did not show any reduction of cell viability in their tested solubility range (≤ 20 µM) for any tested human and non-human cell line (**Figure 1C**). Cysto-180 was more soluble than CN-861-2 and CN-DM-861 and could be tested at higher concentrations. Cysto-180 did not exert cytotoxic effects in Huh-7.5 or CHO-K1 cells with an IC_50_ > 100 µM, underlining the safety of this compound class in the cell viability assay. False negatives resulting from CYS binding to FBS present in the cell media can be excluded, since we observed only a negligible shift in MIC (2-fold) when supplementing the bacterial growth media (MHCII) with FBS (10 %).

In order to assess whether topoisomerase inhibition or previously observed minor groove binding would translate into genotoxicity in a cellular context, a micronucleus assay was performed.^11,8^ The DNA cross-linker mitomycin C (0.05 µM), the intercalator doxorubicin (0.05 µM) and the topoisomerase II poison etoposide (0.25 µM) showed a high amount of nuclear bud/micronuclei formation. Interestingly, etoposide as TOP2 poison showed substantial micronuclei formation far below its IC_50_ for TOP2A (46.3 µM, determined by Inspiralis). Cells treated with CYS at concentrations exceeding their TOP2A IC_50_ (20 µM, 100 µM for Cysto-180) also exhibit some micronuclei/nuclear bud formation, but the extent was comparable to that observed in the DMSO control (**Figure 1D**, **Supplementary** Figure 1). Despite the topoisomerase inhibition observed on the isolated protein, these results suggest a relatively safe genotoxic profile in a whole cell environment, potentially explainable by poor passive eukaryotic membrane permeability of tested CYS derivatives. Nevertheless, further de-risking measures in the development of CYS are highly recommended, as the possibility of a genotoxic effect cannot be excluded.^24,25^

Given the known risk of quinolone antibiotics interfering with the mitochondrial topoisomerases and ETC-proteins, thereby harming mitochondrial functions^26,27^, CYS were further tested in a Seahorse XF Mito Tox Assay (Agilent) to evaluate mitotoxic risks. This assay measures the oxygen consumption rate (OCR) of HepG2 cells treated with the test compounds. During the assay, oligomycin is added as an inhibitor of the oxidative phosphorylation, thus oversaturating the electrochemical gradient at the inner mitochondrial membrane and thereby causing a decrease in the OCR. Subsequently, carbonyl cyanide-p-trifluoromethoxyphenylhydrazone (FCCP) (ionophore) addition leads to an uncoupling of the ETC from the oxidative phosphorylation, thereby increasing the OCR. Inhibition of the oxidative phosphorylation can be seen as a decrease of the starting OCR, whereas uncoupling is defined as an increase in OCR after oligomycin addition. Furthermore, a reduction of the starting OCR and after FCCP addition indicates an inhibition of the ETC caused by the test compound. CYS showed neither inhibition of the oxidative phosphorylation, nor of the ETC due to reduced oxygen consumption at the starting conditions or after FCCP addition, respectively.^28^ However, CYS showed a slight but significant uncoupling of the ETC from the oxidative phosphorylation after oligomycin addition with an uncoupling mitochondrial toxicity index (MTI) of 13-17% at their highest soluble concentrations (**Figure 1E** and **F**). This indicates that CYS as lipophilic weak acids might act as mitochondrial protonophores, shuttling protons across the inner mitochondrial membrane.^29^ CYS are likely to be present in a neutral charge form in the acidic environment of the mitochondrial intermembrane space. This protonation could enhance their membrane migration into the mitochondrial matrix, where the carboxylic acid group is subsequently deprotonated. This process might partially restore the electrochemical gradient and thereby the functionality of the ETC. However, this slight uncoupling did not seem to have a direct harming effect for cell growth and viability as shown above.

### CYS treatment resulted in a significant reduction of superoxide radical formation

It has been shown that anti-infective agents frequently influence the mitochondrial function.^26^ To investigate potential mitochondrial toxicity caused by increased levels of reactive oxygen species (ROS) due to interruption of the ETC upon CYS treatment, superoxide radical (O_2_^•-^) production was examined using MitoSOX Red. ROS are byproducts of various essential biological functions and are involved in cell homeostasis and signaling. However, excess of ROS due to cellular stress, (UV-)irradiation or xenobiotics leads to oxidation of biological components such as lipids, proteins and DNA, which ultimately interrupt their physiological function.^30^ It has been shown that an increase in ROS *in vivo* can lead to *e.g.* heart or liver failure with increased mortality, as seen *e.g*. for the cancer chemotherapeutic class of anthracyclines.^31–33^ The superoxide radical is the precursor of most ROS and thus, represents an indicator used for quantification of ROS formation.

When comparing the superoxide formation of the DMSO control with the CYS treated cells, there was no increase observable. To our surprise, a decrease of superoxide radical levels was observed in CN-861-2 and Cysto-180 treated samples compared to the control group. We also noticed precipitation of CN-DM-861, probably contributing to increased variation in measured fluorescence intensity caused by light scattering at the particles (**Figure 2A** top row, **Figure 2B**). In order to investigate whether CYS are able to intercept superoxide radicals, ROS formation via treatment with the redox-cycler menadione was induced and cells were co-treated with CYS.^34^ Indeed, in the presence of CN-DM-861 and Cysto-180 no significant increase of superoxide was observed. CN-861-2 co-treated cells exhibited a slight yet statistically significant increase in superoxide production, though the induction remained considerably lower compared to the menadione control (**Figure 2A** bottom row, **Figure 2C**).

**Figure 2.**
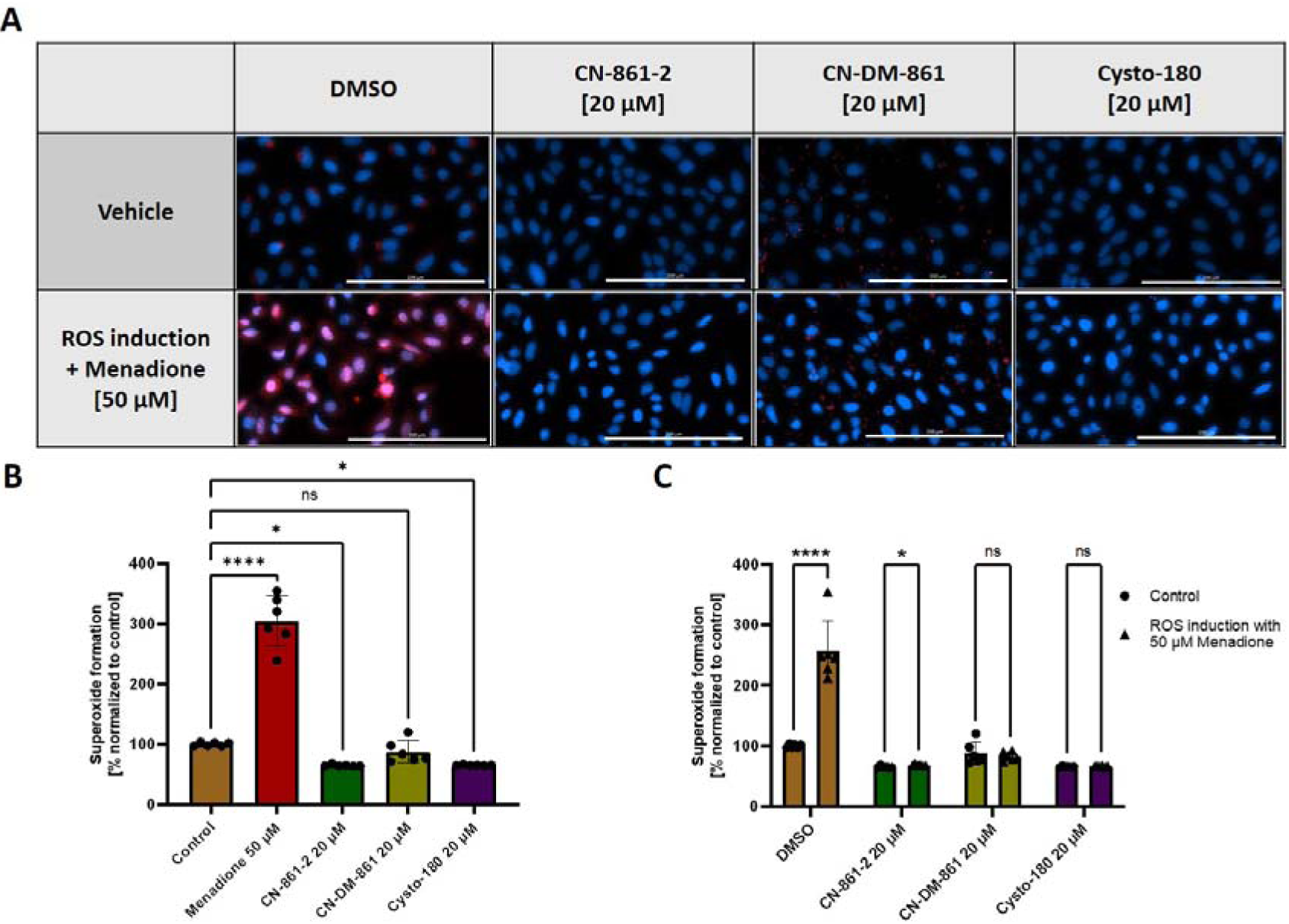
CYS treatment resulted in a significant reduction of superoxide radical formation. **(A)** Superoxide formation in U-2 OS cells (indicated by MitoSOX-red fluorescence (pseudocolor red), Hoechst-stained nuclei (pseudocolor blue)) (scale bars are set to 200 µm). **(B)** Cells showed significantly reduced basal levels of superoxide, when treated with CN-861-2 and Cysto-180. **(C)** Co-treatment with CYS suppresses menadione-induced ROS formation. Mean values with standard deviation are shown. Statistical significance was analyzed by ordinary one-way ANOVA with multiple comparison to the control group (A) and multiple unpaired two-stage step-up t-test (B) (*n* = 6) (ns, non-significant; *, *p* < 0.05; ****, *p* < 0.0001).

Nevertheless, some minor morphological abnormalities were also observed in CYS co-treated cells, indicating that a comprehensive ROS-protection was not achieved. However, these results show that CYS are able to counteract superoxide radical formation induced by menadione, and suggest that CYS generally prevent ROS formation to a certain extent. This effect might partially be explained by the observed slight uncoupling of the ETC. It was previously reported that mild uncoupling by mitochondrial uncoupling proteins (UCPs) reduces superoxide formation due to a slight proton leakage.^35^ This proton leakage potentially decreases the transfer of excess electrons in the ETC to oxygen (electron leakage), which would subsequently lead to superoxide formation.^36^ Furthermore, the structure of CYS could additionally serve as a radical scavenger due to radical resonance stabilization captured in its aromatic moieties.^37^ These findings indicate that CYS do not induce ROS formation but serve as protective agents against oxidative cell stress.

### CYS are metabolized in hepatocytes by amide bond hydrolysis and glucuronidation, which can be suppressed by cobicistat supplementation

Investigation of the *in vitro* drug metabolism and pharmacokinetic (DMPK) properties of potential new drugs is a crucial aspect of its pharmacological and toxicological assessment that is essential for evaluating an efficacy and safety profile. The characterization of new molecules comprises *e.g.* plasma or metabolic half-life and thereby the drug’s ability to reach sufficient exposure *in vivo* in relevant compartments. In addition, the metabolic pathway of a compound can guide compound optimization and has potential impact on toxicological properties, *e.g.* by formation of reactive or toxic intermediates, or by depleting detoxifying agents like glutathione.^38,39^

*In vitro* evaluations revealed metabolic stability of tested CYS in mouse plasma (t_1/2_ > 240 min). In mouse liver microsomes (MLM), some turnover was observed with slight differences between the derivatives. Cysto-180 and CN-861-2 were stable with t_1/2_ > 120 min and 81 % and 59 % of parent compound remaining after 120 min, respectively, while CN-DM-861 had a half-life of 89 min (**Supplementary Table 1**). When tested in murine hepatocytes containing the full complement of phase I and phase II drug metabolizing enzymes, degradation was observed for CN-DM-861 (t_1/2_ 94 ± 15 min), and Cysto-180 (t_1/2_ 41 ± 16 min), while CN-861-2 was stable for over > 3 h (**Supplementary Table 1**). In order to obtain information about the metabolic pathways of CYS, the metabolites of CN-DM-861 and Cysto-180 in mouse hepatocytes were analyzed qualitatively. These studies revealed amide bond hydrolysis and glucuronidation of the parent compound as the major metabolic pathways (**Supplementary** Figures 2-5). In particular, the amide bond between ring C and ring D appeared to be metabolically labile.

With the aim of reducing the turnover of CYS *in vivo* by reducing hepatic uptake, we investigated its metabolic stability in the presence of cobicistat. This drug is known as an OATP1B and CYP3A inhibitor and is used as a co-treatment with HIV therapeutics to enhance their metabolic stability.^40,41^

Supplementation substantially increased the metabolic stability of CN-DM-861 in mouse hepatocytes in a concentration-dependent manner. Whether the observed increase in CYS stability is solely related to reduced transport into hepatocytes via OATP inhibition or also influenced by reduced CYP-mediated amide cleavage^42^, cannot be fully answered by these assays. The fact that CN-DM-861 metabolism was also reduced by cobicistat in liver microsomes shows that in principle, reduction of CYS metabolism via CYP inhibition is feasible (**Supplementary** Figure 6).^42^ In any case, combination of CYS with cobicistat represents a promising strategy to enhance their metabolic stability and availability *in vivo*. However, verification in murine pharmacokinetic studies is required to determine whether a combination with cobicistat leads to improved *in vivo* stability.

All compounds showed high plasma protein binding (PPB ∼ 96-100%, **Supplementary Table 1**), which might raise an issue *in vivo* due to reduced availability of free drug. Only the free compound fraction is capable of interacting with its antimicrobial target and thus, capable of causing a pharmacological effect. However, previous experiments demonstrated the capability of CYS for being efficacious *in vivo* as assessed in various murine infection models.^15,43^ It is also important to consider that partial PPB might even be beneficial for increased metabolic half-life and prolonged exposure at the site of infection.

### *In vivo* toxicity evaluation of CYS in zebrafish embryos revealed no abnormalities in development, cardiac function, or liver morphology

*In vitro* assays are sufficient in providing a basic understanding of the pharmacological properties of a new compound on a cellular level but can often not cover the complexity of a whole organism. Zebrafish embryos can be used as a more complex *in vivo* model *e.g*., for the early evaluation of potential (organo-)toxicity of a compound and its potential phase I and phase II metabolites.^44–46^ In compliance with the 3R principle on the reduction, refinement and replacement of animal experiments, we applied this model since zebrafish at early developmental stages (≤ 120 hours post-fertilization, hpf) are not classified as animal models according to Directive 2010/63/EU.^47^

Developmental toxicity can be caused by various mechanisms including interference of the compound with gene expression, cell growth and differentiation, homeostasis or inhibition of angiogenesis.^48,49^ To investigate potential developmental and *in vivo* genotoxicity, zebrafish embryos were incubated in the presence of CYS from a very early developmental stage (1 day post fertilization (dpf)) until most organs are fully developed (5 dpf). For the positive control 3,4-dichloroaniline^50^ various malformations including spinal curvature and edema were observed, which ultimately led to death. CYS-treated zebrafish embryos did not develop malformations with 100% survival until 5 dpf at their highest tested soluble concentration (20 µM) (**Figure 3A**). Due to its higher solubility in the incubation medium, Cysto-180 was tested up to 200 µM, showing the same outcome. Thus, the maximum tolerated concentration (MTC) of CYS in zebrafish embryos is above their solubility limit. In view of their potent antibacterial activity with minimum inhibitory concentrations (MICs) in the sub-µM range^15^, these data suggest that CYS might have a large therapeutic window.

**Figure 3.**
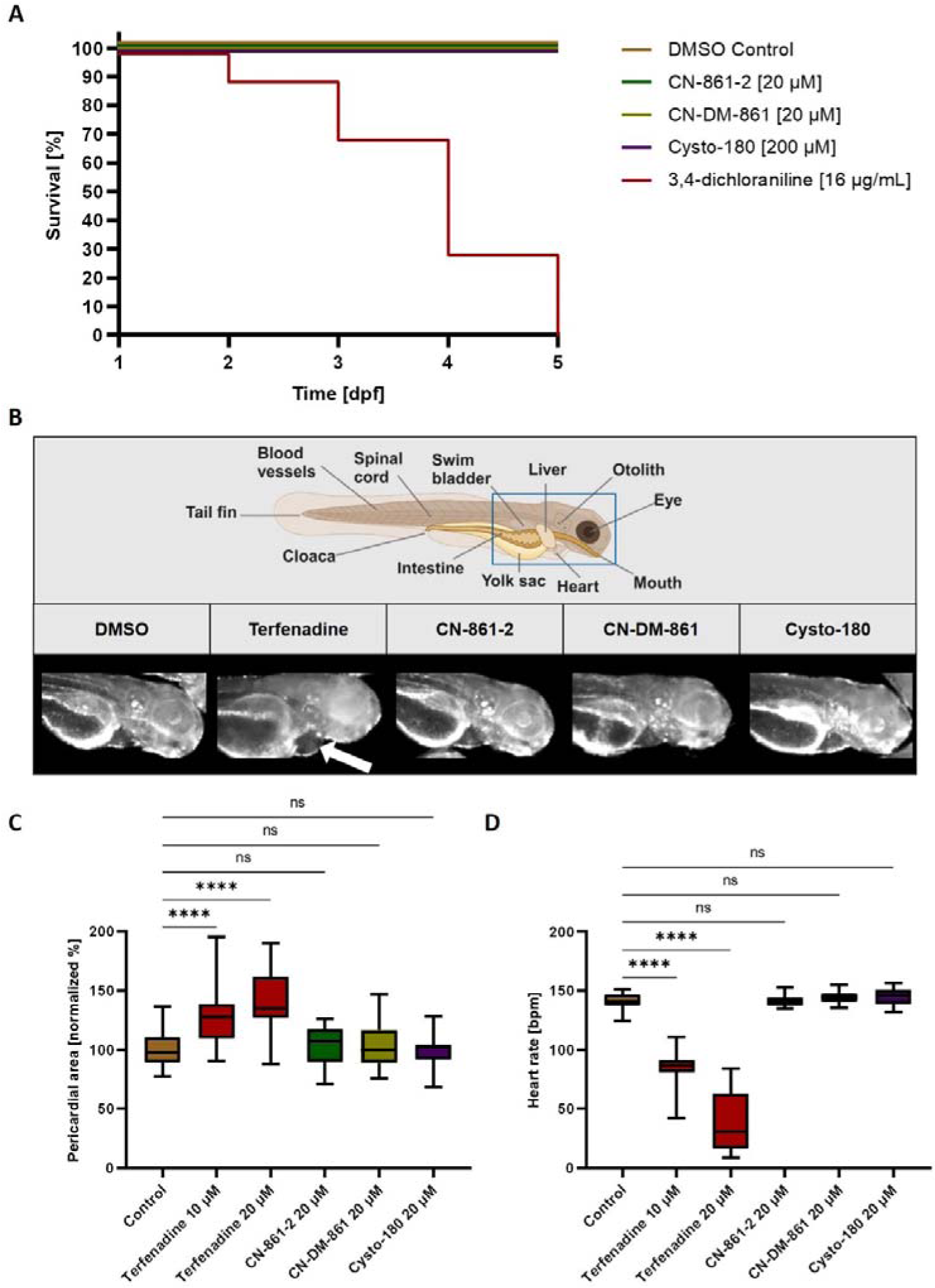
*In vivo* toxicity evaluation of CYS in zebrafish embryos revealed no abnormalities in development or cardiac function. **(A)** CYS showed no developmental *in vivo* toxicity. All CYS treated zebrafish embryos survived and showed no abnormal development when exposed from 1 to 5 dpf. Maximum tolerated concentration (MTC) of CYS was shown to be above their solubility limit (*n* = 20). **(B)** For assessment of cardiotoxicity, zebrafish embryos were treated from 2 to 3 dpf with no observable pathological cardiac morphology for CYS. For the positive control terfenadine, pericardial edemas were observable (white arrow) (created with BioRender.com). **(C)** Statistical evaluation of the pericardial area showed no abnormalities for CYS treated embryos, but an increase in pericardial area after terfenadine treatment. **(D)** Statistical evaluation of the embryós heart rates showed no abnormalities after CYS treatment, but a decreased heart rate after treatment with the positive control terfenadine. Box plot (quartiles Q1 to Q3, including median) with whiskers (min to max) is shown. Statistical significance was analyzed by ordinary one-way ANOVA with multiple comparison to the control group (*n* ≥ 20) (ns, non-significant; ****, *p* < 0.0001).

Furthermore, it was reported that substances, which are cardiotoxic in humans, show the same effect in zebrafish embryos with a very high accuracy.^51^ Especially human ether-à-go-go-related gene (hERG) channel inhibition is regarded as a major risk factor being predictive for cardiotoxicity. Inhibition of this ion channel induces cardiac QT interval prolongation, resulting in a specific type of arrhythmia called torsades de pointes, which is associated with high mortality rates.^21,52^ The phenotypical endpoints determined in the zebrafish embryo assay included heartbeat rate, cardiac rhythm and the development of pericardial edema.^45^ Upon treatment with the known hERG-inhibitor terfenadine, embryos showed a significant, concentration-dependent increase in pericardial area (**Figure 3B** and **C**). Additionally, examination of the heartbeat rate revealed a significant and concentration-dependent decrease of beats per minute with apparent arrhythmia in the vast majority of treated embryos (**Figure 3D**). CYS-treated embryos neither showed pericardial malformations, nor significant changes in pericardial area or heartbeat rate (**Figure 3B-D**). These results suggest a favorable cardiotoxic safety profile of CYS *in vivo*.

The correlation between human and zebrafish toxicology was also observed for drug-induced hepatotoxicity. The assessment of *in vivo* hepatotoxicity offers the opportunity to identify not only toxic effects by the parent molecule but also potential toxicity of *in vivo* metabolites. At the same time the actual *in vivo* distribution of a drug and its metabolites is considered, which can cause adverse effects if a compound shows accumulation in certain compartments such as the liver. Functional toxicity, such as bile acid pump inhibition or steatosis, is also more effectively assessed *in vivo* than *in vitro*. For intoxicated zebrafish embryos, liver degradation can be easily observed by quantifying liver size reduction and observation of histological changes after treatment.^46^

Several mechanisms of drug-induced hepatotoxicity were proposed for the positive control valproic acid, namely formation of reactive metabolites, disturbance of the mitochondrial function and fatty acid metabolism and finally induction of oxidative stress.^31^

The transgenic fish line [Tg(fabp10a:DsRed; elaA:EGFP)] with fluorescent liver cells was used, facilitating the quantification of liver size.^46^ Valproic acid-treated embryos showed significant liver size reduction, reflecting its hepatotoxicity. CYS did not show any significant reduction in liver size, thus suggesting an advantageous and non-hepatotoxic profile *in vivo* (**Figure 4A** and **B**).

**Figure 4.**
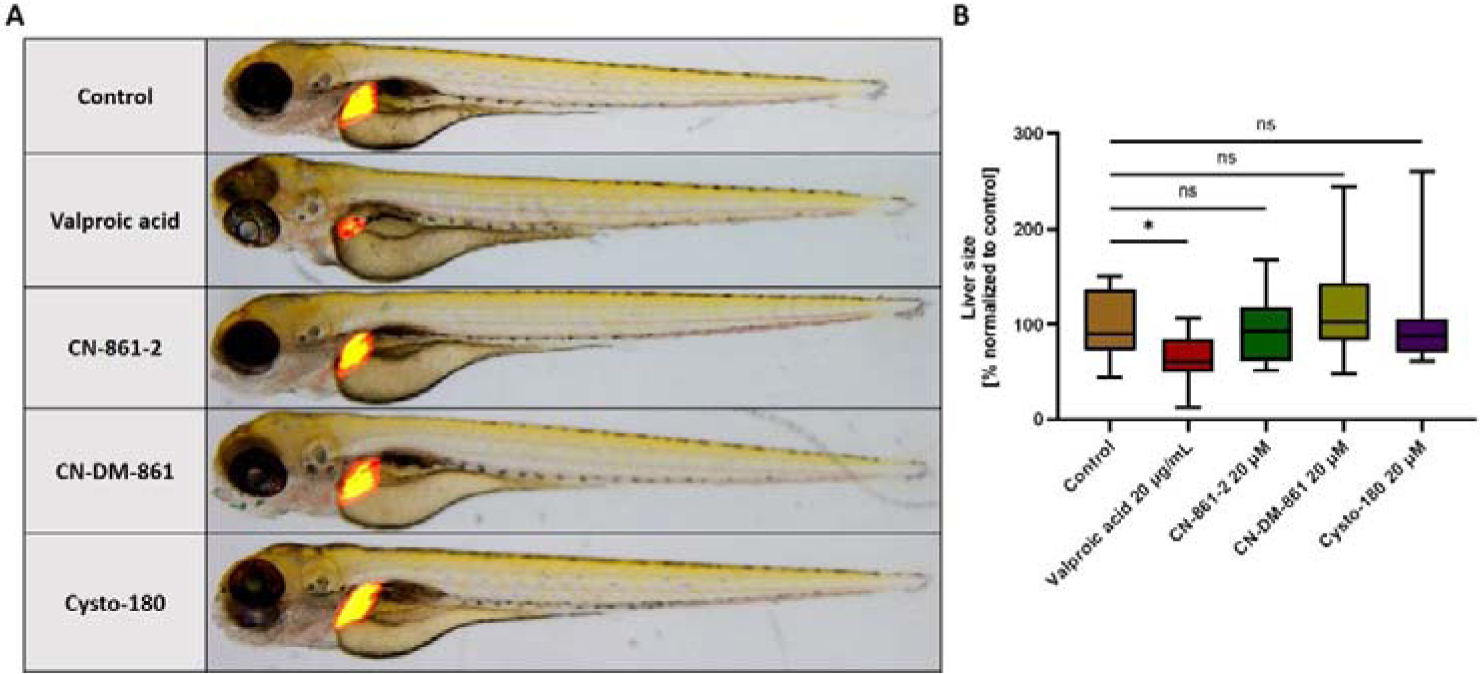
*In vivo* toxicity evaluation of CYS in zebrafish embryos revealed no abnormalities in liver morphology. **(A)** No *in vivo* hepatotoxic effect was observed for CYS treated zebrafish embryos. Transgenic zebrafish embryos [Tg(fabp10a:DsRed; elaA:EGFP)] were checked for liver size reduction after treatment as endpoint readout. Liver degeneration was examined by comparing the fluorescent liver area to the control group. **(B)** CYS treated embryos did not show a significant reduction in liver size, in contrast to the positive control valproic acid. Box plot (quartiles Q1 to Q3, including median) with whiskers (min to max) is shown for a quantitative representation of liver size. Statistical significance was analyzed by ordinary one-way ANOVA with multiple comparison to the control group (*n* ≥ 15) (ns, non-significant; *, *p* < 0.05).

### Molecular off-target identification revealed SCARB1 as primary binding partner of CYS in eukaryotes

Phenotypical assays offer a comprehensive overview of potential intoxications; nevertheless, they frequently fail to provide insights into molecular mechanisms including binding partners of the investigated compounds.

For this purpose we designed and synthesized the CYS photo-probe Cysto-354 to investigate eukaryotic target proteins of CYS via affinity-based protein profiling (AfBPP) (**Supplementary** Figures 7–27).^53^ Cysto-354 was tested for potential cytotoxicity, revealing a slight effect on cell viability at the highest assay concentration (37 µM) and no effect at any of the lower concentrations tested (≤ 12.3 µM) (**Supplementary** Figure 28).

AfBPP aims to identify molecular binding partners via photo-reactive cross-linking followed by enrichment of the bound proteins. In order to exclude enriched but unspecifically bound proteins, competition of the binding site was performed by co-treatment with the unfunctionalized parent compound (here: CN-861-2). In such cases, specific binding is identified by reduction of enrichment in the presence of the parent compound.

By performing the AfBPP assay with Cysto-354-treated HepG2 cells, an immortalized liver cell line which is widely used in early toxicity profiling of drug candidates, we were able to identify significantly enriched proteins, which belong to a functional cluster of cholesterol transfer activity due to lipoprotein and lipid binding (**Figure 5A** and **B**). Proteins within this functional cluster had in common, that they interact with the highly enriched cholesterol-, lipid- and lipoprotein-receptor scavenger receptor class B member 1 (SCARB1).

**Figure 5.**
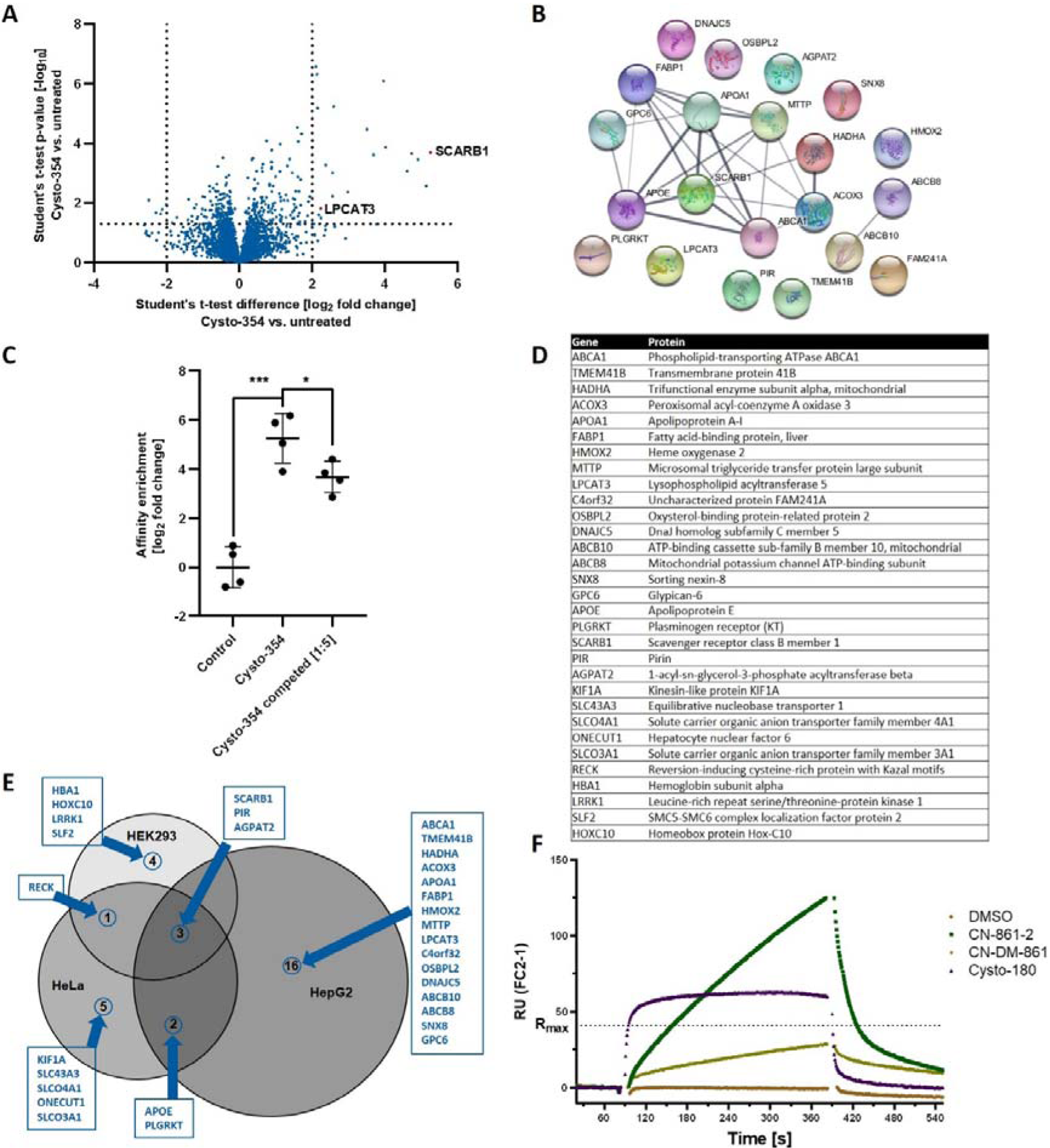
Molecular off-target identification revealed SCARB1 as primary binding partner of CYS in eukaryotes. **(A)** Affinity-based protein profiling (AfBPP) revealed significant enrichment of the SCARB1 protein. HepG2 cells were treated with the CYS photo-probe Cysto-354. After affinity enrichment, the protein abundances were compared to an untreated control (volcano plot). Significant enrichment was analyzed by two-tailed unpaired student’s t-test (cut-offs for enrichment: *p* [–log_10_] > 1.3 and abundance difference [log_2_] > 2) (*n* = 4). **(B)** STRING database analysis of significantly enriched proteins showed functional clustering with respect to cholesterol transfer activity due to lipoprotein and lipid binding (GO-term false discovery rate: 5.05e-7) (created using STRING db (version 12.0)). **(C)** SCARB1 showed the highest enrichment and significant competition upon CN-861-2 addition, indicating specific binding by CYS. Data are shown as mean ± standard deviation. Significant enrichment and competition was analyzed by two-tailed unpaired student’s t-test (*, *p* < 0.05; ***, *p* < 0.001) (*n* = 4). **(D)** List of significantly enriched proteins after Cysto-354 treatment in HepG2, HeLa-CCL-2 and HEK293. **(E)** Venn diagram of enriched proteins for CYS AfBPP in HeLa, HEK293 and HepG2 cells (created using BioVenn). **(F)** Binding analysis using surface plasmon resonance (SPR) confirmed binding of CYS to the SCARB1 protein, exceeding theoretical maximal response (R_max_).

By comparing significantly enriched and competed proteins, we were able to identify SCARB1 (**Figure 5C**) and LPCAT3 as potential specific binding partners of CYS. To gain insights across diverse cell types, this assay was replicated utilizing both HEK293 (kidney) and HeLa (cervix) cells. SCARB1 was observed to be enriched in all tested cell types, whereas LPCAT3 was not enriched in HEK293 or HeLa cells (**Figure 5D** and **E**). The transcriptional coregulator pirin (PIR) and the acyltransferase AGPAT2 were observed to be enriched in all cell types (**Figure 5E**), however, the proteins were not competed by CN-861-2 addition, indicating only unspecific binding of Cysto-354.^54^ Noteworthy, human topoisomerase (TOP1, TOP2A, TOP2B) enrichment was not observed in the whole cell environment of HepG2, HeLa and HEK293, potentially explaining the lack of observable cellular (geno-)toxicity (**Supplementary** Figure 29).

In conclusion, SCARB1 appears to be the primary off-target protein of CYS in eukaryotic cells. Biophysical binding analysis via surface plasmon resonance (SPR) (Biacore X100, Cytiva) was used to characterize and confirm the direct interaction of CYS and SCARB1. For this, biotinylated SCARB1 (Acrobiosystems) was immobilized on a SPR sensorchip, coated with streptavidin (sensor chip SA, Cytiva). All tested CYS derivatives (20 µM) showed interaction with the immobilized SCARB1 protein, exhibiting different binding kinetics (**Figure 5F**). However, CYS appeared to not only interact specifically, as indicated in the competition assay, but also unspecifically with SCARB1. This can be concluded from the observed exceeding of the theoretical maximal response R_max_ (∼41 RU) with increasing CYS contact time and concentration, which did not result in saturation. This effect is most likely mediated by the previously observed π-π-stacking of CYŚs aromatic systems, which occurs subsequently to initial specific binding.^55^ Consequently, we were not able to determine the equilibrium dissociation constant (K_D_) of CYS to SCARB1 reliably. Nevertheless, encouraged by this finding, we further characterized SCARB1 binding and functional inhibition with regards to a potential novel therapeutic indication of CYS.

### Binding of CYS to SCARB1 leads to functional inhibition of HCVpp entry into hepatocytes

The functional inhibition of SCARB1 was evaluated to examine the effects of CYS interaction with the protein.

SCARB1 binds different ligands such as phospholipids, cholesterol esters, lipoproteins, phosphatidylserine, but especially high density lipoproteins (HDL) with high affinity. Thus, SCARB1 plays a dual role in cholesterol metabolism. Firstly, it promotes cholesterol efflux in peripheral tissues such as the arterial wall, aiding in the removal of excess cholesterol. Secondly, it serves as a substantial receptor for HDL cholesterol particles, facilitating the transfer of cholesterol, lipids and lipoproteins back into the liver.^56–58^ Thus, it plays a crucial role in many physiological and pathophysiological processes including cholesterol and lipid homeostasis, cardiovascular disease, liver disease and cancer.^59–61^ Importantly, SCARB1 also serves as a crucial entry receptor for the hepatitis C virus (HCV) into hepatocytes.^62,63^ Chronic hepatitis C is a viral infection that causes liver cirrhosis and hepatocellular carcinoma. In 2019, 1.5 million people were newly infected with the hepatitis C virus, with 290,000 infection-related deaths in the same year.^64^ After infection, HCV circulates in the bloodstream as lipoviral particle and thereby gets in contact with the basolateral surface of hepatocytes to which it attaches by low affinity interaction of lipoviral-associated ApoE to low-density lipoprotein (LDL) receptors and glycosaminoglycans (GAGs). Subsequently, SCARB1 binds the lipoviral-associated lipoproteins, initiating the lipid- and cholesterol-transfer activity of the protein, which probably mediates dissociation of the virus particle from its associated lipoproteins. Importantly, SCARB1 binds the HCV surface glycoprotein E2, causing a conformational change, which enables binding of E2 by CD81. Consequently, CD81 mediates lateral movement and interaction of HCV with claudin-1 (CLDN1), facilitating its endosomal cell entry and release of the viral genome (**Figure 6A**).^65^

**Figure 6.**
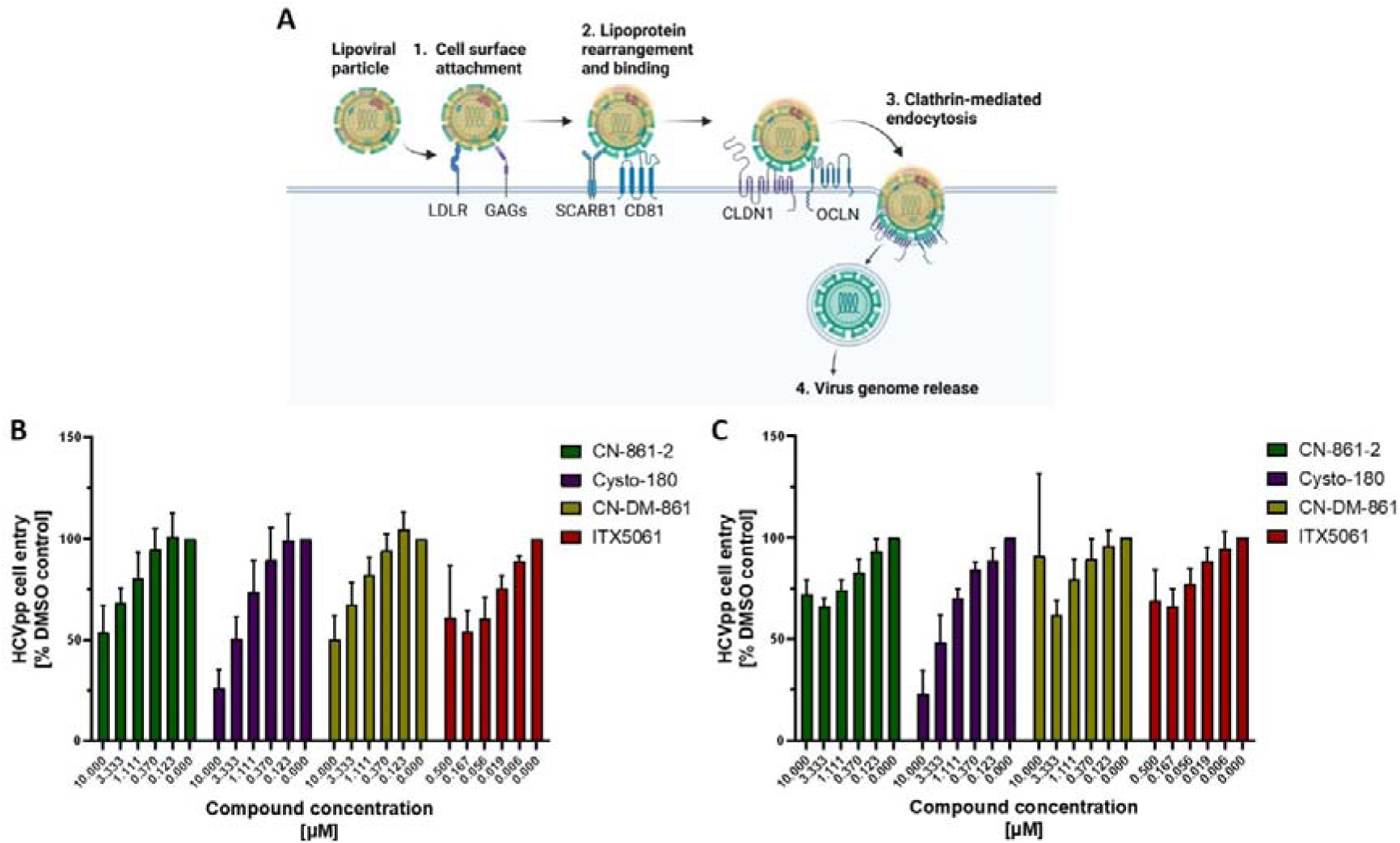
Binding of CYS to SCARB1 leads to functional inhibition of HCVpp entry into hepatocytes. **(A)** SCARB1 is known for its crucial role in binding and transfer of HCV lipoviral particles into hepatocytes (created with BioRender.com). **(B)** HCVpp entry assays with glycoprotein JFH-1 (genotype 2a). Binding of CYS to SCARB1 leads to concentration dependent inhibition of HCVpp entry into hepatocytes. ITX5061 was used as positive control. **(C)** HCVpp entry assays with glycoprotein Con1 (genotype 1b) also showed concentration dependent entry inhibitory activity of CYS. Data are shown as mean values with standard deviation (*n* = 5).

We investigated the functional interaction of CYS with SCARB1 by examining the reduction of HCV pseudoparticle (HCVpp) cell entry into hepatocytes (Huh-7.5) upon CYS treatment. To this end, we used the lentivirus-based pseudotype assay, where the HCVpp was equipped with glycoproteins derived from two different genotypes (JFH1 from genotype 2a and Con1 from genotype 1b). The SCARB1-inhibitor ITX5061 was used as positive control.^61^ The luciferase reporter gene was used to quantify HCVpp entry. CYS treatment successfully inhibited HCVpp cell-entry in a concentration-dependent manner for both tested genotypes. CN-861-2 and CN-DM-861 exhibited similar inhibitory capacities, while Cysto-180 was the most effective derivative with 50% entry inhibition at a concentration of ∼3.3 µM (**Figure 6B** and **C**).

To elucidate the specificity of CYS in inhibiting SCARB1 as the entry receptor of HCVpp, we examined the effect of CYS to the SCARB1-independent entry of vesiculovirus pseudoparticles (VSVpp) equipped with respective viral glycoprotein. Indeed, we demonstrated that CYS exhibited no inhibitory effect on VSVpp cell entry (**Supplementary** Figure 30). Hence, these results provide evidence for the specificity of functional inhibition of SCARB1 as cause of CYS-mediated HCVpp cell entry inhibition.

Applying a bioinformatic molecular docking approach (MOE), only one potential CYS binding site within the modeled structure of SCARB1 was identified. The determined binding pocket is located at the extracellular domain of the protein, potentially important for interaction with native binding partners. CYS blocking of this site might hinder interaction of these binding partners with SCARB1, potentially explaining the inhibitory properties of CYS against HCVpp entry.^66^ However, based on the docking results, multiple poses of CYS in the binding pocket appear feasible (**Supplementary** Figure 31). Future studies will validate the mode of binding of CYS to SCARB1 in more detail and enable the opportunity for structure-guided optimization towards efficient inhibition of this host target.

These findings could not only broaden the potential therapeutic applications of CYS from broad-spectrum antibiotic to antiviral agent, but also extend to various non anti-infective indications. The inhibition of SCARB1 by CYS might affect its lipid- and cholesterol transporter activity *in vivo*. In previous *in vivo* mouse studies, suppression of SCARB1 by ITX5061 successfully led to increased beneficial HDL cholesterol levels and partially reduced atherosclerotic lesions.^61^ In addition, inhibition of SCARB1 is currently under investigation for potential therapeutic benefits across various medical conditions, including arteriosclerosis and other cardiovascular diseases, non-alcoholic fatty liver disease, or specific cancer types associated with the upregulation of SCARB1.^59–61,67,68^

### Summary and Conclusion

Herein, we demonstrated the safety of cystobactamids in cell culture assays and in *in vivo* zebrafish embryo models. The antibiotics showed slight uncoupling of the mitochondrial ETC, which might contribute to their ROS protective properties. This influence on the ETC does not seem to have harming effects *in vitro* or *in vivo* as assessed by cell viability as well as developmental, cardio- and hepatotoxicity in zebrafish embryos. *In vitro* DMPK studies revealed glucuronidation and amide bond hydrolysis as the main biotransformation pathways of cystobactamids. Their metabolic stability was substantially enhanced by supplementation with cobicistat, an OATP- and CYP-inhibitor, offering the opportunity to explore combination therapy as viable approach to improve PK/PD properties of cystobactamids *in vivo*. In addition, on a molecular level, cystobactamids bound to the cholesterol and lipoprotein receptor SCARB1 as the main eukaryotic off-target proteins in the whole cell environment. By binding to SCARB1, they actively prevented the entry of hepatitis C virus into hepatocytes. Analysis of the binding mode to SCARB1 might guide further development and optimization of cystobactamids, along with related compound classes such as albicidin and coralmycin, for various possible therapeutic areas.

## Materials and Methods

### Cell culture

All cell types were cultivated at 37 °C with 5 % CO_2_. HepG2, HEK293, HCT116, CHO-K1, HeLa-CCL-2 and Huh-7.5 were purchased at ATCC. The respective cell media (Gibco) was supplemented with 10 % FBS (Gibco) before usage. Cells were used between passage #5 and #30. To obtain biological repeats, cells were split separately for at least two passages.

### Topoisomerase IIα inhibition

Inhibition of topoisomerase activity was examined using human topoisomerase II alpha decatenation assay kit (Inspriralis) as indicated by the manufacturer. Briefly, a compound dilution series was prepared in DMSO. Water, dilution buffer, assay buffer, plasmid and enzyme were mixed with compound solution and incubated at 37°C for 30 min. A positive control (100% activity) was prepared without compound and a negative control (0% activity) was prepared without compound and enzyme. The reaction was stopped by adding gSTEB-buffer and chloroform/isoamylalcohol (24:1 v/v). The samples were vortexed and centrifuged before running gel-electrophoresis. The gel was stained with ethidium bromide and imaged using a Fusion Fx gel imager (Vilber Lourmat). Subsequent analysis was done using ImageJ and GraphPad Prism (Version 10.0.2).

### Cytotoxicity

Cells were washed with PBS and 0.5 mL trypsin was added. Afterwards, cells were incubated for 5 min before adding 10 mL medium containing 10 % FBS. Per well, 120 µL cell suspension (5×10^4^ cells/mL) was seeded in transparent 96 well cell bind plates and incubated for 2 h (37 °C, 5 % CO_2_). Dilution series (1:3) of test compounds was prepared in respective media containing 10 % FBS and added (60 µL) to the prepared cell plate. Cells were incubated for further 72 h (37 °C, 5 % CO_2_). Next, 20 µL MTT (5 mg/mL in PBS) was added to each well and incubated for 2 h (37 °C, 5 % CO_2_). The wells were emptied and 100 µL isopropanol/10 N HCl (1000:4) per well was added.

The plates were analyzed by measuring the absorbance at 570 nm (plate reader Infinite® 200 Pro, Tecan). After normalization of data to the respective solvent controls, the calculated percentage of growth inhibition was plotted using GraphPad Prism software (version 10.0.2).

### Micronucleus test

CHO-K1 cells were washed and seeded like described above with a concentration of 5×10^4^ cells/mL. Cells were treated with CYS (20 µM, 100 µM for Cysto-180) and incubated for 24 h. Mitomycin C (0.05 µM), doxorubicin (0.05 µM) and etoposide (0.25 µM) were used as positive controls. Cells were washed and stained with Hoechst (5 µg/mL) (Thermo) in F12 media for 15 min. Subsequently, cells were washed three times with PBS and imaged using Celldiscoverer 7 (Zeiss).

### Mitochondrial toxicity

Seahorse mitotox assay was carried out according to Agilent seahorse XF mito tox assay kit user guide (kit 103595-100, Agilent). Briefly, sensor cartridge was hydrated at 37 °C, non-CO_2_, one day before the assay started in 200 µL water and 2 h in calibrant solution before usage. HepG2 cells were seeded with a density of 10×10^3^ per well in 80 µL and incubated overnight at 37 °C, 5 % CO_2_. At the day of the assay, 100 mL Seahorse XF DMEM medium was prepared by addition of glucose (10 mM), pyruvate (1.0 mM) and glutamine (2.0 mM). CYS compound dilutions were prepared in assay medium as well as positive controls menadione (50 µM) and Rot/AA (1.0 µM). Cells were washed with assay medium and compounds were added, followed by 2 h incubation at 37 °C, non-CO_2_. Oligomycin (1.5 µM) and FCCP (1.0 µM) were prepared and added to the respective ports in the sensor cartridge. The utility plate with sensor cartridge was inserted into the Seahorse system (Agilent) and calibrated. Afterwards the cell plate was added and measurement of the oxygen consumption rate (OCR) was started. The assay was repeated three times independently with at least 6 technical replicates per condition. Uncoupling MTI was calculated as follows:

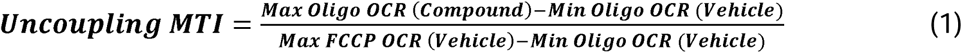

### Mitochondrial superoxide formation

CYS were tested for their potential of inducing oxidative stress in U-2 OS cells. Therefore, 120 µL of a cell suspension in McCoy’s (+10 % FBS) containing 5 x 10^4^ cells/mL were seeded into black 96-well imaging plate (BD Falcon). The plates were incubated for two days (37 °C, 5 % CO_2_) until the cells reached approximately 70 % confluence. Subsequently, the cells were washed with HBSS buffer, prior to adding CYS (20 µM) and the ROS inducer menadione (50 µM) as positive control, along with the staining solution (HBSS, calcium, magnesium, 5 µM MitoSOX red and 10 µg/mL Hoechst (Thermo)). To check for ROS protective properties, menadione (50 µM) was added directly to the staining solution. The cells were incubated for 1-2 h at 37 °C and 5 % CO_2_. Afterwards, the cells were washed twice with HBSS. The cells were imaged using an automated fluorescence microscope (Celldiscoverer 7, Zeiss) (excitation: 485 nm, emission: 535 nm) to analyze the fluorescence intensity of the superoxide tracker MitoSOX red. The experiment was carried out with six replicates per condition. Quantification was done by high-content image analysis using the ZEN software (version 3.4, blue edition, Zeiss). Data presentation was done using GraphPad Prism (version 10.0.3) with ordinary one-way ANOVA including multiple comparison to the control group for statistical analysis of sample without ROS induction via menadione in the staining solution. Multiple unpaired t-test was used to compare samples for ROS protective properties, which were incubated with menadione in the staining solution.

### Metabolic stability in mouse liver microsomes

For the evaluation of phase I metabolic stability, the compound (1 μM) was incubated with 0.5 mg/mL pooled mouse liver microsomes (Xenotech, Kansas City, USA), 2 mM NADPH, 10 mM MgCl_2_ at 37 °C for 120 min on a microplate shaker (Eppendorf, Hamburg, Germany). The metabolic stability of testosterone, verapamil and ketoconazole was determined in parallel to confirm the enzymatic activity of mouse liver microsomes. For combination experiments, cobicistat was added to the incubation mixture together with test compounds. The incubation was stopped after defined time points by precipitation of aliquots of enzymes with 2 volumes of cold internal standard solution (15 nM diphenhydramine in 10% methanol/acetonitrile). Samples were stored on ice until the end of the incubation and precipitated protein was removed by centrifugation (15 min, 4 °C, 4,000 g). Concentration of the remaining test compound at the different time points was analyzed by HPLC-MS/MS (Vanquish Flex coupled to a TSQ Altis Plus, Thermo Fisher, Dreieich, Germany) and used to determine half-life (t_1/2_).

### Metabolic stability in mouse hepatocytes

For the evaluation of combined phase I and phase II metabolic stability, the compound (1 μM) was incubated with 0.25 x 10^6^ cells/mL of pooled mouse hepatocytes (Xenotech, Kansas City, USA). Cells were thawed in Leibovitz’s L-15 medium without phenol red (ThermoFisher Scientific, Waltham, USA). Briefly, cells were transferred into 50 mL of medium, followed by centrifugation at 55 g for 6 min. Supernatant was discarded and the cell count determined after gently resuspending the cell pellet in 1 mL of medium. Hepatocytes were diluted to 0.5 x 10^6^ cells/mL and incubated at 37 °C, 700 rpm for 10 min, to achieve the desired final cell count after addition of an equal volume of test compounds in medium, leading to final test concentration of 1 µM at 1% DMSO. Samples were incubated for 240 min at 37 °C, 700 rpm and the incubation was stopped after defined time points by precipitation of aliquots in 4 volumes cold internal standard solution (12.5 nM diphenhydramine in 10% methanol/acetonitrile). For the determination of metabolism in presence of cobicistat, this drug was added together with test compounds in the desired concentration range. The metabolic stability of testosterone, verapamil, ketoconazole and 7-hydroxycoumarine were determined in parallel to confirm the enzymatic activity of mouse hepatocytes. Samples were stored on ice until the end of the incubation and precipitated protein was removed by centrifugation (15 min, 4 °C, 4,000 g). Concentration of the remaining test compound at the different time points was analyzed by HPLC-MS/MS (Vanquish Flex coupled to a TSQ Altis Plus, Thermo Fisher, Dreieich, Germany) and used to determine half-life (t_1/2_). Intrinsic clearance was calculated as follows:

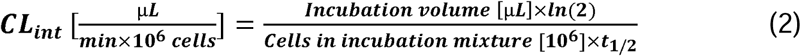

For metabolite idenfication studies, test compounds were incubated at 10 µM final concentration and samples were analyzed using HPLC-HRMS (Ultimate 3000 coupled to a Q Exactive Focus, Thermo Fisher, Dreieich, Germany). LC conditions were as follows: column: Accucore Phenyl-Hexyl (2.6 µm, 100 x 2.1 mm; Thermo Fisher, Dreieich, Germany); temperature 40 °C; flow rate 0.500 mL/min; solvent A: water + 0.1% formic acid; solvent B: acetoni-trile + 0.1% formic acid; gradient^69^: 0–4.0 min 2–35% B, 4.0–7.0 min 35–98% B, 7.0–8.0 min 98% B, 8.0–10.0 min 2% B.

MS analysis was performed using full scan mode (switching polarity, full MS resolution 35,000, scan range 200–2,000; data-dependent MS/MS (ddMS^2^) resolution 17,500, stepped collision energy with 17.5, 35, 52.5). Blank samples using DMSO were run in parallel for background subtraction. Sample processing for metabolite identification was performed usind Compound Discoverer 3.2 (Thermo Fisher, Dreieich, Germany). Metabolites were identified based on mass shifts and feasibility of the metabolic reaction also in view of MS peak intensities over time.

### Zebrafish handling

Handling of adult zebrafish and experiments with zebrafish embryos were performed in accordance to the EU directive 2010/63/EU and the German Animal Welfare Act (§11 Abs. 1 TierSchG). An automatic aquatic ecosystem (PENTAIR, Apoka, UK) was used for zebrafish housing. Adult zebrafish were mated pairwise in our zebrafish facility after one night in the mating cages (light/dark cycle: 14 h/10 h). The eggs were collected after 2 h mating and placed in a petri dish with fish water (pH 7.36 ± 0.08, conductivity: 800 ± 50 µs). The eggs were washed with 0.3x Danieaús medium (17 mM NaCl, 0.2 mM KCl, 0.12 mM MgSO_4_, 0.18 mM Ca(NO_3_)_2_, 1.5 mM HEPES, pH 7.1–7.3, and 1.2 µM methylene blue). 300 eggs were placed per petri dish and stored in the incubator at 28 °C overnight. 24 h post mating, the unfertilized eggs were discarded. The 0.3x Danieaús media in the petri dishes was replaced every 24 h and embryos with developmental issues were euthanized using ice water. Embryo-/genotoxicity, cardiotoxicity and hepatotoxicity were tested using zebrafish embryos younger than 120 hours post-fertilization (hpf). The wild-type AB line was used for embryo-/genotoxicity and cardiotoxicity, and the transgenic line Tg(fabp10a:DsRed; elaA:EGFP) was used for hepatotoxic evaluation. Embryos were treated with phenylthiourea (PTU) starting from 1 day post fertilization (dpf) for hepato- and cardiotoxicity analysis. Previous to compound addition, embryos were dechorionated using pronase (1 mg/mL) and washed several times. The embryos were euthanized on ice after maximum 120 hpf and frozen at −20 °C.

### Maximum tolerated concentration (MTC)

Zebrafish embryos (AB line) were dechorionated with pronase at 1 dpf. For embryo-/genotoxicity testing, 20 embryos per condition were treated with 20 µM CN-861-2, CN-DM-861 (solubility limit) and 200 µM (solubility limit), 100 µM, 50 µM and 20 µM Cysto-180 from 1 dpf till 5 dpf. DMSO (0.2 %) was used as negative control and 3,4-dichloroaniline (16 µg/mL) as positive control. For acute *in vivo* toxicity testing, embryos were treated from 4 to 5 dpf accordingly with maximal soluble concentrations of CYS and DMSO (0.2 %) and 3,4-dichloroaniline (32 and 64 µg/mL) as negative and positive controls, respectively. Kaplan-Meier curves were generated using GraphPad Prism (version 10.0.3, GraphPad, Boston, MA, USA). Dead embryos were defined when no heart beat was observable.

### In vivo cardiotoxicity in zebrafish embryos

Embryos were prepared as described above. At 2 dpf, the embryos were placed in 6 well plates with 10 embryos per well. Embryos were treated with CYS (20 µM) and the positive control terfenadin (10, 20 µM) in Danieaús media. Danieaús with DMSO was used as negative control. At least 20 embryos were tested per condition. Embryos were incubated at 28 °C for 24 h. Afterwards, embryos were anesthetized with tricaine added to the media (40 µg/mL, 0.004 % (*w/v*)) and videos were recorded using a stereomicroscope (1.25x magnification) (Stemi 508, Zeiss) with Media Recorder (version 4.0) und analyzed via DanioScope software (version 1.2.208, Noldus). The embryos were checked with attention to specific cardiotoxic phenotypes defined as decreased heartrate, arrhythmia and pericardial edema. Death was defined as absence of heartbeat. Data presentation was done using GraphPad Prism (version 10.0.3) with ordinary one-way ANOVA including multiple comparison to the control group and multiple unpaired t-test for statistical analysis.

### In vivo hepatotoxicity in zebrafish embryos

Zebrafish embryos [Tg(fabp10a:DsRed; elaA:EGFP)] were prepared as described above. Embryos were placed in 6 well plates with 10 embryos per well. Treatment with CYS as well as the positive control valproic acid (20, 40 µg/mL) was done by soaking from 3 to 5 dpf in 0.3x Danieaús media. Per condition, at least 15 embryos were treated. Danieaús media with DMSO was used as negative control. The embryos were incubated at 28 °C. After 48 h treatment, the embryos were checked via a fluorescence stereomicroscope (M205 FA, Leica) with attention to liver degeneration. Fluorescence images were taken after anesthetizing the embryos with tricaine (40 µg/mL) and placing them laterally (excitation: 558 nm, emission 583 nm, 35x magnification). The specific phenotypic hepatotoxic endpoint was liver degeneration defined as liver size reduction. The liver size was calculated using ImageJ software (format: 8 bit, threshold: 40-255, binary, particle size: 10.0-infinite). The liver size of the control embryos was set to 100%. The relative liver sizes of treated embryos were calculated accordingly and normalized to the control group. Data presentation was done using GraphPad Prism (version 10.0.3) with ordinary one-way ANOVA including multiple comparison to the control group for statistical analysis.

### Affinity-based protein profiling (AfBPP) sample preparation

Cells were seeded in 100 mm cell-bind petri dishes with a concentration of 2.8 x 10^6^ cells per sample and incubated (37 °C, 5 % CO_2_) for 3 days to reach ∼80 % confluence. Cells were washed with warm PBS prior to compound addition. Affinity based proteome profiling was performed by treating the respective cell line with Cysto-354 (2.5 µM) for 3 h. For competition samples, cells were treated with CN-861-2 (12.5 µM) for 1h, prior to addition of photo-probe Cysto-354 (2.5 µM) for 3 h incubation time. Control samples were treated with DMSO. Afterwards, cells were UV-irradiated for 10 minutes on ice, scraped off and transferred to Eppendorf tubes for washing with 1 mL cold PBS (centrifuge 5 min, 500xg, 4 °C). The cell pellet was stored at −80 °C until lysis. Further sample preparation was done as previously described.^13^ Briefly, the cell pellet was lysed in 0.4 % SDS in PBS by sonication (Bandelin Sonoplus). After proteome adjustment (1000 µg/sample), azide-alkyne cycloaddition of labeled proteins with biotin was performed. Subsequently, the proteome was precipitated and washed with acetone and methanol, followed by avidin-bead enrichment. Afterwards, samples were digested (trypsin platinum, Promega), desalted and dried in a speedVac, before solving the samples in 1 % FA. Samples were filter (Merck Millipore, UFC30GV0S) and transferred to HPLC autosampler vials (QuanRecovery, Waters).

### LC-MS measurement of HepG2 proteome

Peptides were measured and online-separated using an UltiMate 3000 nano HPLC system (*Dionex*) coupled to a *Bruker* timsTOF Pro mass spectrometer via a CaptiveSpray nano-electrospray ion source and *Sonation* column oven. Peptides were first loaded on the trap column (Acclaim PepMap 100 C18, 75 µm ID x 2 cm, 3 µm particle size, *Thermo Scientific*), washed with 0.1 % formic acid in water for 7 min at 5 µL/min and subsequently transferred to the separation column (*IonOpticks* Aurora C18 column, 25 cm × 75 µm, 1.7 µm) and separated over a 60 min gradient from 5 % to 28 % B, then to 40 % B over 13 min, followed by 10 min at 95 % before re-equilibration and at a flow rate of 400 nL/min. The mobile phases A and B were 0.1 % (v/v) formic acid in water and 0.1 % (v/v) formic acid in acetonitrile, respectively. The timsTOF Pro was operated in data-dependent PASEF mode with the dual TIMS analyzer operating at equal accumulation and ramp times of 100 ms each with a set 1/K_0_ ion mobility range from 0.85 to 1.40 V × s × cm^-2^. The capillary voltage of the CaptiveSpray source was set to 1500 V. 10 PASEF scans per topN acquisition cycle were performed, resulting in a total cycle time of 1.17 s. The mass range was set from 100 to 1700 m/z. Only Precursors reaching an intensity threshold of 1750 arbitrary units were considered for fragmentation, precursors reaching a target intensity of 14500 arbitrary units were dynamically excluded for 0.4 min. The quadrupole isolation width was set to 2D*m/z* for *m/z*D<D700 and to 3D*m/z* for *m/z*D>D800. The collision energy was ramped linearly as a function of the mobility from 59DeV at 1/K_0_D=D1.6DV × s × cm^-2^ to 20DeV at 1/K_0_D=D0.6DV × s × cm^-2^. TIMS elution voltages were calibrated linearly to obtain the reduced ion mobility coefficients (1/K_0_) using three Agilent ESI-L Tuning Mix ions (*m/z* 622, 922 and 1,222) spiked on the CaptiveSpray Source inlet filter.

MS raw data was analyzed using MaxQuant software (version 2.0.3.0) and peptides were searched against Uniprot database for Homo sapiens (taxon identifier: 9606, downloaded on 11.03.2022, canonical, reviewed). Carbamidomethylation of cysteine was set as fixed modification and oxidation of methionine and acetylation of *N*-termini were set as variable modifications. Trypsin was set as proteolytic enzyme with a maximum of 2 missed cleavages. For main search, precursor mass tolerance was set to 4.5 ppm and fragment mass tolerance to 0.5 Da. Label free quantification (LFQ) mode was activated with a LFQ minimum ratio count of 2. Second peptide identification was enabled, and false discovery rate (FDR) determination carried out by applying a decoy database and thresholds were set to 1 % FDR at peptide-spectrum match and at protein levels and “match between runs” (0.7 min match and 20 min alignment time windows) option was enabled. Normalized LFQ intensities extracted from the MaxQuant result table proteinGroups.txt were further analyzed with Perseus software (version 2.03.1).

### LC-MS measurement of HeLa-CCL-2 and HEK293 proteome

Sample analysis was done as previously described.^13^ Samples have been analyzed using nanoElute nano flow liquid chromatography system (Bruker, Germany) coupled to a timsTOF Pro (Bruker, Germany). Loading of the samples to the trap column (Thermo Trap Cartridge 5 mm) was performed, followed by washing with 6 µL 0.1 % FA with a flow rate of 10 µL/min. Transferring of the peptide samples to the analytical column (Aurora Ultimate CSI 25 cm x 75 µm ID, 1.6 µm FSC C18, IonOpticks) was done, with subsequent separation by a gradient elution (eluent A: H2O + 0.1 % FA, B: ACN + 0.1 % FA; 0 % to 3 % in 1 min, 3 % to 17 % in 57 min, 17 % to 25 % in 21 min, 25 % to 34 % in 13 min, 34 % to 85 % in 1 min, 85 % kept for 8 min) using a flow rate of 400 nL/min. Captive Spray nanoESI source (Bruker, Germany) was applied for ionizing the peptides at 1.5 kV with 180 °C dry temperature at 3 L/min gas flow. timsTOF Pro (Bruker, Germany) was operated using default dia-PASEF long gradient mode with TIMS set to 1/K0 start at 0.6 Vs/cm^2^, end at 1.6 Vs/cm^2^, with a ramp and accumulation time of 100 ms each and a ramp rate of 9.43 Hz. Mass range was set from 100.0 Da to 1700 Da with positive ion polarity. Dia-PASEF mass range was arranged to 400.0 Da to 1201.0 Da with a mobility range of 0.60 1/K0 to 1.43 1/K0 and a cycle time of 1.80 s. Collision energy for 0.60 1/K0 was fixed to 20.00 eV and ramped for 1.6 1/K0 to 59.00 eV. Tuning MIX ES-TOF (Agilent) was applied for calibration of *m/z* and mobility. Raw data were processed using DIA-NN (version 1.8.1), and proteins were identified against Uniprot Homo sapiens reference proteome (Proteome ID: UP000005640, downloaded 27/12/2023). Default settings were used, except precursor charge range was from 2 to 4. C-carbamidomethylation was set as fixed modification. To allow further data processing with Perseus Software, “--relaxed-prot-inf” was added in additional options. Further data analysis was performed in Perseus software (version 2.0.5.0), were the values were transformed to their log_2_-value and the biological replicates were grouped. To allow the comparison of the whole datasets, missing values were imputed by default settings and the differential protein abundance between different treatment regimens were evaluated using two-tailed student’s t-test. The cut-offs for –log_10_ *P* was set to 1.3 (*P* = 0.05) and for t-test difference > 2. Proteins fitting these thresholds were seen as significantly enriched compared to the control only treated with DMSO. Significant competition was evaluated comparing CN-861-2 co-treated samples with Cysto-354-only treated samples.^13^

### Surface plasmon resonance (SPR)

Binding analysis of CYS to SCARB1 was done by using SPR (Biacore X100, Cytiva). Therefore, biotinylated SCARB1 (Acrobiosystems) was immobilized on a SPR sensor chip SA (Cytiva) on flow cell (FC) 2. Afterwards, CYS (20 µM) were solved and centrifuged in running buffer. CYS were tested by interaction analysis over the reference FC1 and FC2, where SCARB1 has been immobilized. To conclude binding, the subtraction of FC2-1 was analyzed. HBS-EP+ (Cytiva) with 1 % DMSO was used as running buffer.

### HCVpp cell entry assay

Lentiviral pseudotypes were prepared by 293T transfection as described previously.^70^ 293T cells were seeded in a density of 3×10^6^ cells in 10 cm plates and incubated at 37 °C and 5 % CO_2_. After 24 hours, the cells were transfected with 2 µg of the lentiviral Gag-Pol expression construct pCMV-ΔR8.74^71^, 2 µg of the reporter plasmid coding for a firefly luciferase (pWPI-F-Luc-BLR)^72,73^ and 2 µg of either pcDNA3_CMV_dcE1E2_Con1, pcDNA3_CMV_dcE1E2_JFH^74^ or pczVSV-G^75^ coding for the glycoproteins of interest or 2 µg of the empty vector control (pcDNA3). For transfection, the plasmids were mixed in Opti-MEM containing a final concentration of 0.035 mg/ml polyethylenimine (PEI) and afterwards added to the cells. After overnight incubation, sodium butyrate was added at a final concentration of 10 mM. After 6 hours of incubation, a medium change was performed and after a subsequent overnight incubation, the pseudoparticle containing supernatant was harvested and cleared of cell debris by passing through 0.45 µm pore size filter and used for entry assay.

The evaluation of effect on entry by the compounds was performed on Huh-7.5 cells. Huh-7.5 cells were seeded in 96-well plates at a density of 7.2 x 10^3^/well and incubated for 24 hours at 37 °C and 5 % CO_2_. Compounds were diluted in six subsequent 3-fold dilutions with a final DMSO concentration of 1 % and freshly harvested pseudoparticles were added and transferred to the cells. After 72 hours of incubation, the cells were lysed with lysis buffer (1 % triton-X-100, 25 mM glycylglycine, 15 mM MgSO_4_, 4 mM EGTA, 1 mM dithiothreitol) and frozen at −20°C. The luciferase activity measurements were performed by transferring 72 µL of assay buffer (25 mM glycylglycine, 5 mM KPO_4_, 50 mM MgSO_4_, 10 mM EGTA, 2 % ATP, 1 mM dithiotreitol) to a white 96-well plate and adding 20 µL of the cell-lysis suspension with subsequent adding of 40 µL D-Luciferin and immediate measuring using the Berthold LB960 Centro XS3 plate luminometer.

### Molecular Docking

For molecular docking, the AlphaFold structure (AF-Q8WTV0-F1) of scavenger receptor class B member 1 was downloaded as a PDB file and loaded into MOE (version 2022.02). Structural issues were corrected using the “Structure Preparation” tool and the protein was protonated using “Protonate3D” with a set pH to 7.4. The structure was energy minimized before screening for a potential binding site with “Site Finder”. Atom dummies were loaded in the potential binding site. CYS derivatives Cysto-180, CN-861-2 and CN-DM-861 were prepared (protonated, energy minimized) in MOE and loaded into a compound database as a .mdb file. This .mdb file was used for docking the compounds to the previous prepared SCARB1 structure. Dummy atoms were selected as target site with triangle matcher (score: London dG) as placement method and induced fit (score: GBVI/WSA dG) for refinement. Potential poses were browsed and optimized via protonation and energy minimization.

### Large Language Models (LLMs)

During the preparation of this work, the author(s) used ChatGPT (GPT-4) in order to polish the phrasing of some passages. After using this tool, the author(s) reviewed and edited the content as needed and take(s) full responsibility for the content of the publication.

## Data Availability

Datasets and further information of current and ongoing related studies are available upon request from the corresponding author. The mass spectrometry proteomics data have been deposited to the ProteomeXchange Consortium via the PRIDE partner repository with the data set identifiers PXD053711 (HepG2) and PXD052895 (HEK293, HeLa).

Reviewers access details (HepG2):

Project accession: PXD053711

Token: KfP437WEj9a6

Username

Password: zFzWzExpYFTk

Reviewers access details (HEK293 and HeLa):

Project accession: PXD052895

Token: nyCCMJvkf4bK

Username

Password: IrKDq3L4vxby

## Acknowledgements

This study was funded by the German Federal Ministry of Education and Research (OpCyBac-project grant number: 16GW0219K). The funder played no role in study design, data collection, analysis and interpretation of data, or writing of this manuscript.

## Author Contributions

TR designed, performed and analyzed the majority of mentioned experiments and wrote the manuscript. DH, TS, DS, DK synthesized and provided the cystobactamid derivatives. AMK performed and analyzed the *in vitro* pharmacokinetic experiments. JHo supported the seahorse assay. JSH and DM supported proteomic sample preparation and data analysis. BH performed and analyzed the HCVpp assay and cytotoxicity testing on Huh-7.5 cells. FF supported the topoisomerase assays and helped writing the manuscript. FD helped with scientific advice and manuscript revision. JH, RM, MB, AK, AKK, TP and SS provided supervision and funding. TR, JH and RM designed the project. All authors were involved in reviewing and editing the manuscript and approved the final version.

## Competing Interests

All authors declare no financial or non-financial competing interests.

## Notes

Supporting information for this article is provided separately

### Competing Interest Statement

The authors have declared no competing interest.

### Summary of Updates

Supplementary Information has been added and the manuscript file was updated

## Bibliography

1. WHO Bacterial Priority Pathogens List, 2024.; 2024.

2. Nathan C, Cars O. Antibiotic Resistance - Problems, Progress, and Prospects. New Engll J Med. 2014;371(19):1761–1763.

3. Rossolini GM, Mantengoli E. Antimicrobial resistance in Europe and its potential impact on empirical therapy. Clin Microbiol Infect. 2008;14(SUPPL. 6):2–8. doi:10.1111/j.1469-0691.2008.02126.x

4. European Medicines Agency. The European Medicines Agency Road map to 2015D: The Agency’s contribution to Science, Medicines, Health. 2015;44(October 2011). http://www.ema.europa.eu/docs/en_GB/document_library/Report/2011/01/WC500101373.pdf

5. Butler MS, Henderson IR, Capon RJ, Blaskovich MAT. Antibiotics in the clinical pipeline as of December 2022. J Antibiot (Tokyo). 2023;76(8):431–473. doi:10.1038/s41429-023-00629-8

6. Baumann S, Herrmann J, Raju R, et al. Cystobactamids: Myxobacterial topoisomerase inhibitors exhibiting potent antibacterial activity. Angew Chemie - Int Ed. 2014;53(52):14605–14609. doi:10.1002/anie.201409964

7. Hüttel S, Testolin G, Herrmann J, et al. Discovery and Total Synthesis of Natural Cystobactamid Derivatives with Superior Activity against Gram-Negative Pathogens. Angew Chemie - Int Ed. 2017;56(41):12760–12764. doi:10.1002/anie.201705913

8. Michalczyk E, Hommernick K, Behroz I, et al. Molecular mechanism of topoisomerase poisoning by the peptide antibiotic albicidin. Nat Catal. 2023;6(1):52–67. doi:10.1038/s41929-022-00904-1

9. Kim YJ, Kim HJ, Kim GW, et al. Isolation of Coralmycins A and B, Potent Anti-Gram Negative Compounds from the Myxobacteria Corallococcus coralloides M23. J Nat Prod. 2016;79(9):2223-2228. doi:10.1021/acs.jnatprod.6b00294

10. Hashimi SM. Albicidin, a potent DNA gyrase inhibitor with clinical potential. J Antibiot (Tokyo). 2019;72(11):785–792. doi:10.1038/s41429-019-0228-2

11. Elgaher WAM, Hamed MM, Baumann S, et al. Cystobactamid 507: Concise Synthesis, Mode of Action, and Optimization toward More Potent Antibiotics. Chem - A Eur J. 2020;26(32):7219–7225. doi:10.1002/chem.202000117

12. Cheng B, Müller R, Trauner D. Total Syntheses of Cystobactamids and Structural Confirmation of Cystobactamid 919-2. Angew Chemie - Int Ed. 2017;56(41):12755–12759. doi:10.1002/anie.201705387

13. Risch T, Kolling D, Mostert D, Herrmann J, Müller R. Point mutations in the ygiV promoter region lead to cystobactamid resistance and reduced virulence in. ResearchSquare. Published online 2024. 10.21203/rs.3.rs-3751821/v1 License:

14. Moeller M, Norris MD, Planke T, et al. Scalable Syntheses of Methoxyaspartate and Preparation of the Antibiotic Cystobactamid 861-2 and Highly Potent Derivatives. Org Lett. 2019;21(20):8369–8372. doi:10.1021/acs.orglett.9b03143

15. Testolin G, Cirnski K, Rox K, et al. Synthetic studies of cystobactamids as antibiotics and bacterial imaging carriers lead to compounds with high: In vivo efficacy. Chem Sci. 2020;11(5):1316–1334. doi:10.1039/c9sc04769g

16. Harrison RK. Phase II and phase III failures: 2013-2015. Nat Rev Drug Discov. 2016;15(12):817–818. doi:10.1038/nrd.2016.184

17. Arrowsmith J, Miller P. Trial Watch: Phase II and Phase III attrition rates 2011-2012. Nat Rev Drug Discov. 2013;12(8):569. doi:10.1038/nrd4090

18. Lipsky MS, Sharp LK. From idea to market: The drug approval process. J Am Board Fam Pract. 2001;14(5):362–367.

19. Muller PY, Milton MN. The determination and interpretation of the therapeutic index in drug development. Nat Rev Drug Discov. 2012;11(10):751–761. doi:10.1038/nrd3801

20. Kalyaanamoorthy S, Barakat KH. Development of Safe Drugs: The hERG Challenge. Med Res Rev. 2018;38(2):525–555. doi:10.1002/med.21445

21. Redfern WS, Carlsson L, Davis AS, et al. Relationships between preclinical cardiac electrophysiology, clinical QT interval prolongation and torsade de pointes for a broad range of drugs: Evidence for a provisional safety margin in drug development. Cardiovasc Res. 2003;58(1):32–45. doi:10.1016/S0008-6363(02)00846-5

22. Information S. Cystobactamid 507: Concise Synthesis, Mode of Action, and Optimization toward More Potent Antibiotics.

23. Fotakis G, Timbrell JA. In vitro cytotoxicity assays: Comparison of LDH, neutral red, MTT and protein assay in hepatoma cell lines following exposure to cadmium chloride. Toxicol Lett. 2006;160(2):171–177. doi:10.1016/j.toxlet.2005.07.001

24. Minotti G, Menna P, Salvatorelli E, Cairo G, Gianni L. Anthracyclines: Molecular advances and pharmacologie developments in antitumor activity and cardiotoxicity. Pharmacol Rev. 2004;56(2):185–229. doi:10.1124/pr.56.2.6

25. Tomasz M. Mitomycin C: small, fast and deadly (but very selective). Chem Biol. 1995;2(9):575–579. doi:10.1016/1074-5521(95)90120-5

26. Kalghatgi S, Spina CS, Costello JC, et al. Bactericidal antibiotics induce mitochondrial dysfunction and oxidative damage in mammalian cells. Sci Transl Med. 2013;5(192). doi:10.1126/scitranslmed.3006055

27. Hangas A, Aasumets K, Kekäläinen NJ, et al. Ciprofloxacin impairs mitochondrial DNA replication initiation through inhibition of Topoisomerase 2. Nucleic Acids Res. 2018;46(18):9625–9636. doi:10.1093/nar/gky793

28. Tilmant K, Gerets H, De Ron P, Hanon E, Bento-Pereira C, Atienzar FA. In vitro screening of cell bioenergetics to assess mitochondrial dysfunction in drug development. Toxicol Vitr. 2018;52(May):374–383. doi:10.1016/j.tiv.2018.07.012

29. Shrestha R, Johnson E, Byrne FL. Exploring the therapeutic potential of mitochondrial uncouplers in cancer. Mol Metab. 2021;51(March):101222. doi:10.1016/j.molmet.2021.101222

30. Herb M, Gluschko A, Schramm M. Reactive Oxygen Species: Not Omnipresent but Important in Many Locations. Front Cell Dev Biol. 2021;9(September):1–12. doi:10.3389/fcell.2021.716406

31. Chang TKH, Abbott FS. Oxidative stress as a mechanism of valproic acid-associated hepatotoxicity. Drug Metab Rev. 2006;38(4):627–639. doi:10.1080/03602530600959433

32. Maillet A, Tan K, Chai X, et al. Modeling Doxorubicin-Induced Cardiotoxicity in Human Pluripotent Stem Cell Derived-Cardiomyocytes. Sci Rep. 2016;6(January):1–13. doi:10.1038/srep25333

33. Carrasco R, Castillo RL, Gormaz JG, Carrillo M, Thavendiranathan P. Role of Oxidative Stress in the Mechanisms of Anthracycline-Induced Cardiotoxicity: Effects of Preventive Strategies. Oxid Med Cell Longev. 2021;2021. doi:10.1155/2021/8863789

34. Monteiro JP, Martins AF, Nunes C, et al. A biophysical approach to menadione membrane interactions: Relevance for menadione-induced mitochondria dysfunction and related deleterious/therapeutic effects. Biochim Biophys Acta -Biomembr. 2013;1828(8):1899–1908. doi:10.1016/j.bbamem.2013.04.006

35. Cadenas S. Mitochondrial uncoupling, ROS generation and cardioprotection. Biochim Biophys Acta - Bioenerg. 2018;1859(9):940–950. doi:10.1016/j.bbabio.2018.05.019

36. Zhao RZ, Jiang S, Zhang L, Yu Z Bin. Mitochondrial electron transport chain, ROS generation and uncoupling (Review). Int J Mol Med. 2019;44(1):3–15. doi:10.3892/ijmm.2019.4188

37. Haider K, Haider MR, Neha K, Yar MS. Free radical scavengers: An overview on heterocyclic advances and medicinal prospects. Eur J Med Chem. 2020;204. doi:10.1016/j.ejmech.2020.112607

38. Guengerich FP, Liebler DC, Reed DL. Enzymatic Activation of Chemicals to Toxic Metabolites. Vol 14.; 1985. doi:10.3109/10408448509037460

39. Baillie TA. Metabolism and toxicity of drugs. Two decades of progress in industrial drug metabolism. Chem Res Toxicol. 2008;21(1):129–137. doi:10.1021/tx7002273

40. Xu L, Liu H, Murray BP, et al. Cobicistat (GS-9350): A potent and selective inhibitor of human CYP3A as a novel pharmacoenhancer. ACS Med Chem Lett. 2010;1(5):209-213. doi:10.1021/ml1000257

41. Custodio JM, Wang H, Hao J, et al. Pharmacokinetics of cobicistat boosted-elvitegravir administered in combination with rosuvastatin. J Clin Pharmacol. 2014;54(6):649–656. doi:10.1002/jcph.256

42. Peng HM, Raner GM, Vaz ADN, Coon MJ. Oxidative cleavage of esters and amides to carbonyl products by cytochrome P450. Arch Biochem Biophys. 1995;318(2):333–339. doi:10.1006/abbi.1995.1237

43. Kohnhäuser D, Seedorf T, Cirnski K, et al. Optimization of the Central α-Amino Acid in Cystobactamids to the Broad-Spectrum, Resistance-Breaking Antibiotic CN-CC-861. J Med Chem. Published online 2024. doi:10.1021/acs.jmedchem.4c00927

44. Parng C, Seng WL, Semino C, McGrath P. Zebrafish: a preclinical model for drug screening. Assay Drug Dev Technol. 2002;1(1 Pt 1):41-48. doi:10.1089/154065802761001293

45. Zhu JJ, Xu YQ, He JH, et al. Human cardiotoxic drugs delivered by soaking and microinjection induce cardiovascular toxicity in zebrafish. J Appl Toxicol. 2014;34(2):139–148. doi:10.1002/jat.2843

46. He JH, Guo SY, Zhu F, et al. A zebrafish phenotypic assay for assessing drug-induced hepatotoxicity. J Pharmacol Toxicol Methods. 2013;67(1):25–32. doi:10.1016/j.vascn.2012.10.003

47. Strähle U, Scholz S, Geisler R, et al. Zebrafish embryos as an alternative to animal experiments-A commentary on the definition of the onset of protected life stages in animal welfare regulations. Reprod Toxicol. 2012;33(2):128–132. doi:10.1016/j.reprotox.2011.06.121

48. Dathe K, Schaefer C. Drug safety in pregnancy: the German Embryotox institute. Eur J Clin Pharmacol. 2018;74(2):171–179. doi:10.1007/s00228-017-2351-y

49. Vargesson N. Thalidomide-Induced Teratogenesis: History and Mechanisms. Birth Defects Res (Part C). 2015;(105):140–156. doi:10.1002/bdrc.21096

50. Nagel R, Bresch H, Caspers N, et al. Effect of 3,4-dichloroaniline on the early life stages of the zebrafish (Brachydanio retio): Results of a comparative laboratory study. Ecotoxicol Environ Saf. 1991;21(2):157–164. doi:10.1016/0147-6513(91)90017-J

51. Zakaria ZZ, Benslimane FM, Nasrallah GK, et al. Using Zebrafish for Investigating the Molecular Mechanisms of Drug-Induced Cardiotoxicity. Biomed Res Int. 2018;2018. doi:10.1155/2018/1642684

52. Walker DK. The use of pharmacokinetic and pharmacodynamic data in the assessment of drug safety in early drug development. Br J Clin Pharmacol. 2004;58(6):601–608. doi:10.1111/j.1365-2125.2004.02194.x

53. Wright MH, Fetzer C, Sieber SA. Chemical Probes Unravel an Antimicrobial Defense Response Triggered by Binding of the Human Opioid Dynorphin to a Bacterial Sensor Kinase. J Am Chem Soc. 2017;139(17):6152–6159. doi:10.1021/jacs.7b01072

54. Perez-Dominguez F, Carrillo-Beltrán D, Blanco R, et al. Role of pirin, an oxidative stress sensor protein, in epithelial carcinogenesis. Biology (Basel). 2021;10(2):1–13. doi:10.3390/biology10020116

55. Seedorf T, Kirschning A, Solga D. Natural and Synthetic Oligoarylamides: Privileged Structures for Medical Applications. Chem - A Eur J. 2021;27(26):7321–7339. doi:10.1002/chem.202005086

56. Xu S, Laccotripe M, Huang X, Rigotti A, Zannis VI, Krieger M. Apolipoproteins of HDL can directly mediate binding to the scavenger receptor SR-BI, an HDL receptor that mediates selective lipid uptake. J Lipid Res. 1997;38(7):1289–1298. doi:10.1016/s0022-2275(20)37413-7

57. Wen-Jun Shen, Salman Azhar and FBK. SR-B1: A Unique Multifunctional Receptor for Cholesterol Influx and Efflux. Annu Rev Physiol. 2018;176(5):139–148. doi:10.1146/annurev-physiol-021317-121550

58. Ji Y, Jian B, Wang N, et al. Scavenger receptor BI promotes high density lipoprotein-mediated cellular cholesterol efflux. J Biol Chem. 1997;272(34):20982–20985. doi:10.1074/jbc.272.34.20982

59. Mooberry LK, Sabnis NA, Panchoo M, Nagarajan B, Lacko AG. Targeting the SR-B1 receptor as a gateway for cancer therapy and imaging. Front Pharmacol. 2016;7(DEC):1–11. doi:10.3389/fphar.2016.00466

60. Qiu Y, Liu S, Chen HT, et al. Upregulation of caveolin-1 and SR-B1 in mice with non-alcoholic fatty liver disease. Hepatobiliary Pancreat Dis Int. 2013;12(6):630–636. doi:10.1016/S1499-3872(13)60099-5

61. Masson D, Koseki M, Ishibashi M, et al. Increased HDL cholesterol and ApoA-I in humans and mice treated with a novel SR-BI inhibitor. Arterioscler Thromb Vasc Biol. 2009;29(12):2054–2060. doi:10.1161/ATVBAHA.109.191320

62. Zeisel MB, Koutsoudakis G, Schnober EK, et al. Scavenger receptor class B type I is a key host factor for hepatitis C virus infection required for an entry step closely linked to CD81. Hepatology. 2007;46(6):1722–1731. doi:10.1002/hep.21994

63. Scarselli E, Ansuini H, Cerino R, et al. The human scavenger receptor class B type I is a novel candidate receptor for the hepatitis C virus. EMBO J. 2002;21(19):5017–5025. doi:10.1093/emboj/cdf529

64. Global HIV Hepatitis and Sexually Transmitted Infections Programmes. Global Progress Report on HIV, Viral Hepatitis and Sexually Transmitted Infections, 2021.; 2021. https://www.who.int/publications/i/item/9789240027077

65. Lindenbach BD, Rice CM. The ins and outs of hepatitis C virus entry and assembly. Nat Rev Microbiol. 2013;11(10):688–700. doi:10.1038/nrmicro3098

66. Powers HR, Sahoo D. SR-B1’s Next Top Model: Structural Perspectives on the Functions of the HDL Receptor. Curr Atheroscler Rep. 2022;24(4):277–288. doi:10.1007/s11883-022-01001-1

67. Gordon JA, Noble JW, Midha A, et al. Upregulation of scavenger receptor B1 is required for steroidogenic and nonsteroidogenic cholesterol metabolism in prostate cancer. Cancer Res. 2019;79(13):3320–3331. doi:10.1158/0008-5472.CAN-18-2529

68. Wei C, Wan L, Yan Q, et al. HDL-scavenger receptor B type 1 facilitates SARS-CoV-2 entry. Nat Metab. 2020;2(12):1391–1400. doi:10.1038/s42255-020-00324-0

69. A. Kenneth MacLeoda, Kevin-Sebastien Coquelinb, Leticia Huertasc, Frederick R. C. Simeonsa, Jennifer Rileya, Patricia Casadoc, Laura Guijarroc, Ruth Casanuevac, Laura Framea, Erika G. Pintoa, Liam Fergusona, Christina Duncana, Nicole Muttera, Yoko Shis 1. Acceleration of infectious disease drug discovery and development using a humanized model of drug metabolism. Proc Natl Acad Sci. 2024;121. doi:10.1073/pnas

70. Haid S, Grethe C, Bankwitz D, Grunwald T, Pietschmann T. Identification of a Human Respiratory Syncytial Virus Cell Entry Inhibitor by Using a Novel Lentiviral Pseudotype System. J Virol. 2016;90(6):3065–3073. doi:10.1128/jvi.03074-15

71. Dull T, Zufferey R, Kelly M, et al. A Third-Generation Lentivirus Vector with a Conditional Packaging System. J Virol. 1998;72(11):8463–8471. doi:10.1128/jvi.72.11.8463-8471.1998

72. Vieyres G, Welsch K, Gerold G, et al. ABHD5/CGI-58, the Chanarin-Dorfman Syndrome Protein, Mobilises Lipid Stores for Hepatitis C Virus Production. PLoS Pathog. 2016;12(4). doi:10.1371/journal.ppat.1005568

73. Pham HM, Argañaraz ER, Groschel B, Trono D, Lama J. Lentiviral Vectors Interfering with Virus-Induced CD4 Down-Modulation Potently Block Human Immunodeficiency Virus Type 1 Replication in Primary Lymphocytes. J Virol. 2004;78(23):13072–13081. doi:10.1128/jvi.78.23.13072-13081.2004

74. Haid S, Grethe C, Dill MT, Heim M, Kaderali L, Pietschmann T. Isolate-dependent use of claudins for cell entry by hepatitis C virus. Hepatology. 2014;59(1):24–34. doi:10.1002/hep.26567

75. Kalajzic I, Stover ML, Liu P, Kalajzic Z, Rowe DW, Lichtler AC. Use of VSV-G pseudotyped retroviral vectors to target murine osteoprogenitor cells. Virology. 2001;284(1):37–45. doi:10.1006/viro.2001.0903

