## Supplementary Information for "Cystobactamid off-target profiling reveals favorable safety, superoxide reduction, and SCARB1 inhibition in eukaryotes"

^1^T. Risch, Dr. A. M. Kany, J. Hilgers, F. Fries, F. Deschner, Dr. J. Herrmann, Prof. Dr. S. A. Sieber, Prof.

Dr. R. Müller

Helmholtz Institute for Pharmaceutical Research Saarland (HIPS), Helmholtz Centre for Infection

Research (HZI) and Saarland University Department of Pharmacy, Campus Building E8.1, 66123

Saarbrücken, Germany,

^2^T. Risch, Dr. A. M. Kany, Dr. D. Kohnhäuser, D. Heimann, F. Fries, F. Deschner, Prof. Dr. M. Brönstrup,

Prof. Dr. Thomas Pietschmann, Dr. J. Herrmann, Prof. Dr. R. Müller

German Centre for Infection Research (DZIF), Inhoffenstraße 7, 38124 Braunschweig, Germany

^3^B. Hellwinkel, Prof. Dr. Thomas Pietschmann

TWINCORE, Centre for Experimental and Clinical Infection Research, a joint venture between the

Helmholtz Centre for Infection Research and the Hannover Medical School, Hannover, Germany

^4^Dr. D. Mostert, Prof. Dr. S. A. Sieber

Center for Functional Protein Assemblies (CPA), Department of Chemistry, Chair of Organic

Chemistry II, Technical University of Munich, Ernst-Otto-Fischer-Straße 8, 85748 Garching, Germany

^5^Dr. D. Solga, Dr. T. Seedorf, Prof. Dr. A. Kirschning

Leibniz University Hannover, Institute of Organic Chemistry, Schneiderberg 1B, 30167 Hannover,

Germany

^6^Dr. D. Kohnhäuser, D. Heimann, Prof. Dr. M. Brönstrup, Prof. Dr. Thomas Pietschmann

Helmholtz Centre for Infection Research (HZI), Inhoffenstraße 7, 38124 Braunschweig, Germany

^7^Dr. J. Hoppstädter, Prof. Dr. A. K. Kiemer

Institut for Pharmaceutical Biology, Department of Pharmacy, Saarland University

Campus C2 3, 66123 Saarbruecken, Germany

^8^Prof. Dr. Andreas Kirschning

Uppsala Biomedical Center (BMC), Uppsala University, Husargatan 3, 752 37 Uppsala, Sweden

^9^Prof. Dr. Thomas Pietschmann

Cluster of Excellence RESIST (EXC 2155), Hanover Medical School, Hanover, Germany

*Co-corresponding Author

**This file contains:**

Methods for chemical syntheses

Supplementary Figures 1-31

Supplementary Table 1

Supporting References

**Synthesis of CN-861-2, CN-DM-861 and Cysto-180**

CN-861-2 was synthesized according to published literature.^1,2^

HRMS (ESI) calculated 842.2780 [M+H^+^], 842.2759 found.

Purity (LC-MS): 99%.

CN-DM-861 was synthesized according to published literature.^1,2^

HRMS (ESI) calculated 812.2675 [M+H^+^], 812.2675 found.

Purity (LC-MS): 99%.

Cysto-180 was synthesized according to published literature.^3^

HRMS (ESI) calculated 797.2930 [M+H^+^], 797.2935 found.

Purity (LC-MS): 99%.

Micronucleus test

The micronucleus test was performed like described in the methods section. Pictures displayed are exemplary for all wells imaged and examined.

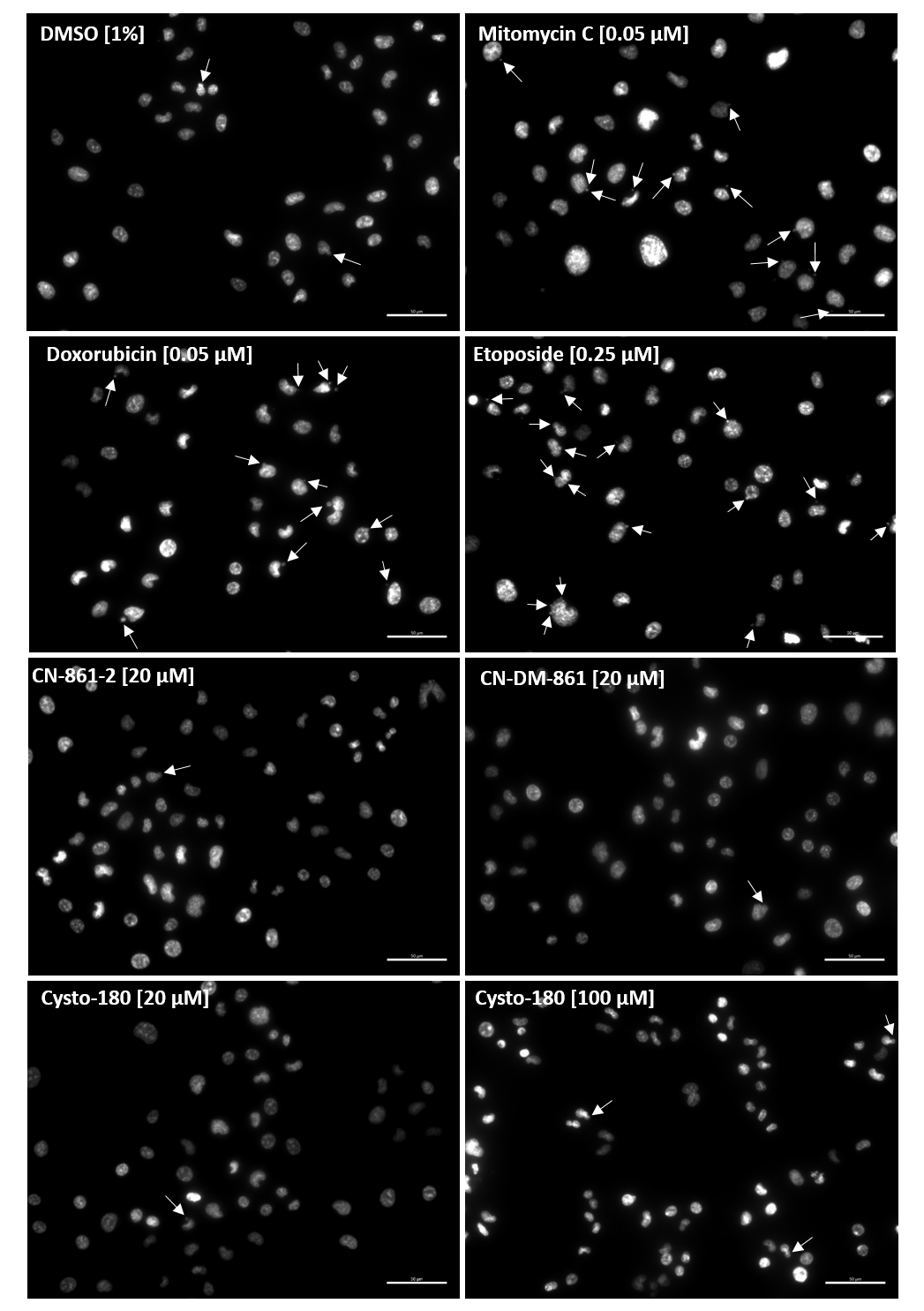

**Supplementary Figure 1 (Related to Figure 1).** Genotoxic evaluation showed extensive micronucleus formation for the positive controls mitomycin C, doxorubicin and etoposide as clastogenic and aneugenic agents. CYS showed micronucleus formation comparable to the DMSO control. (Nuclei colored in white, white arrows indicating micronucleus formation, scale bar is set to 50 µm) (*n* = 3).

***In vitro* drug metabolism and pharmacokinetics (DMPK)**

**Supplementary Table 1.** *In vitro* metabolism studies of CYS derivatives including mouse liver microsomal (MLM), murine S9 fraction, mouse hepatocyte and plasma stability testing as well as plasma protein binding (PPB). (*n* ≥ 2)

| **Code** | **MLM**  **t_1/2_ [min] / Cl_int_ [µl/mg/min] / % remaining at 2 h** | **Mouse Hepatocytes**  **t_1/2_ [min] / Cl_int_ [µl/mg/10^6^ cells]** | **Mouse Plasma**  **t_1/2_ [min]** | **Mouse PPB [%]** |
| --- | --- | --- | --- | --- |
| **CN-861-2** | **>120 / <11.6 / 59 ± 11** | **>180 / <5.1** | **>240** | **99.63 ± 0.16** |
| **CN-DM-861** | **89.0 ± 1.3 / 15.6 ± 0.2** | **94 ± 15 / 10 ± 2** | **>240** | **99.85 ± 0.08** |
| **Cysto-180** | **>120 / <11.6 / 81 ± 10** | **41 ± 16 / 26 ± 12** | **>240** | **95.9 ± 2.4** |

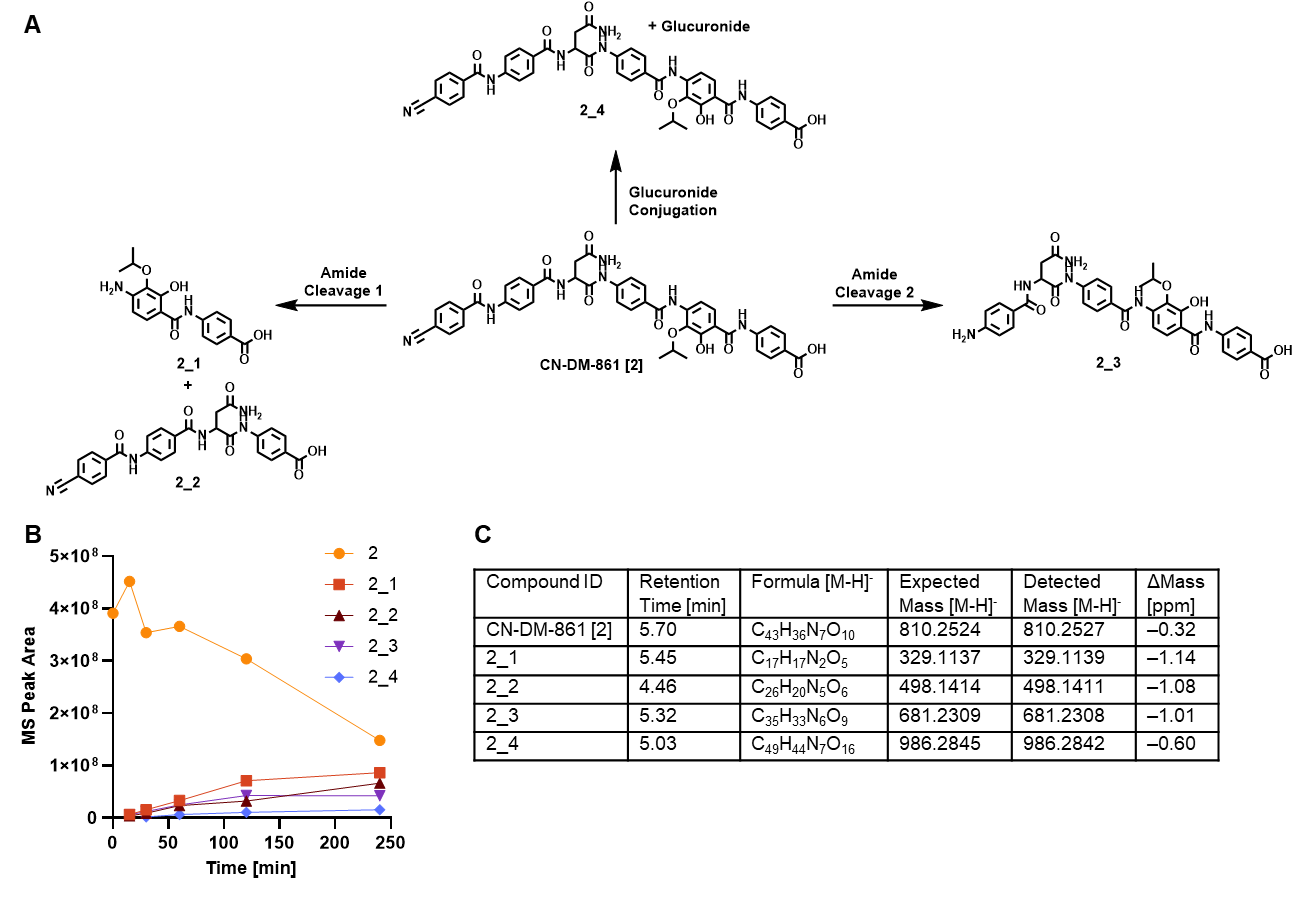

**Supplementary Figure 2.** Metabolite identification (MetID) of CN-DM-861. **(A)** MetID of CN-DM-861 showed amide bond hydrolysis and glucuronidation as main metabolic degradation pathways in mouse hepatocytes. **(B)** Time-dependent analysis of mass signals for the respective compounds showed a correlation between parent compound degradation and increase of identified metabolites, where amide cleavage between rings C and D seemed to be the most prominent path. The exact position of the glucuronidation could not be clearly identified between the hydroxy and the carboxy group. **(C)** Corresponding chromatographic and MS data of identified metabolites. (See also Supplementary Figure 3)

**2**

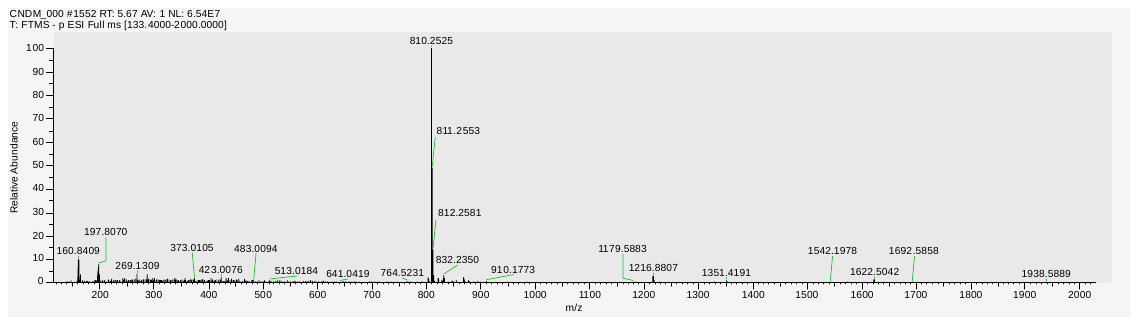

**2_1**

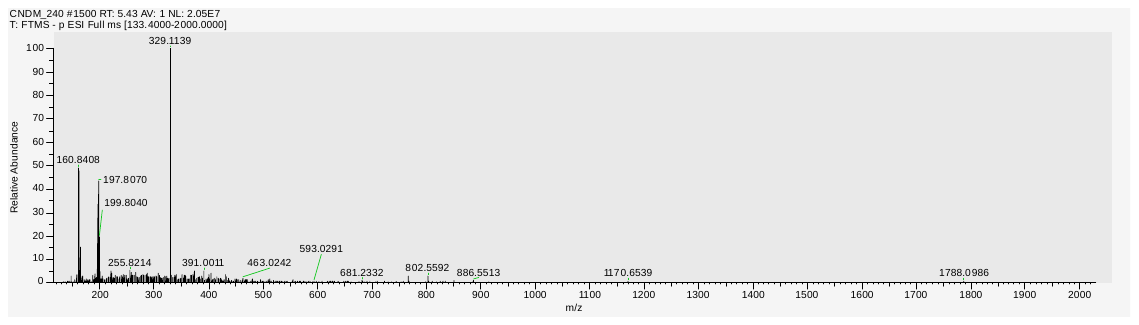

**2_2**

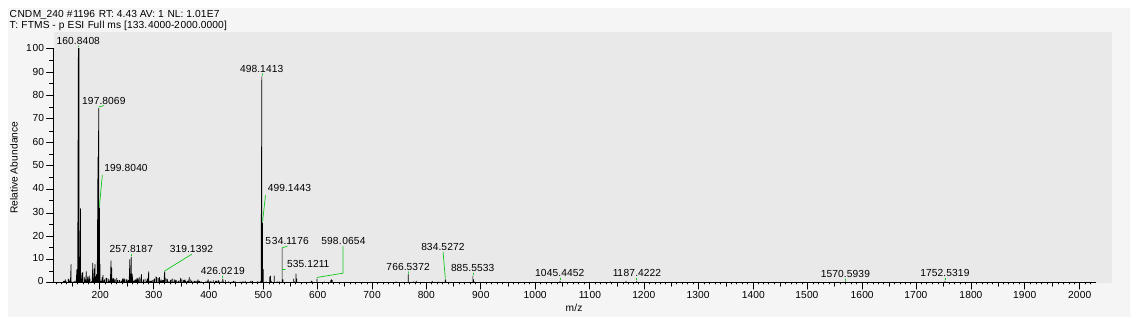

**2_3**

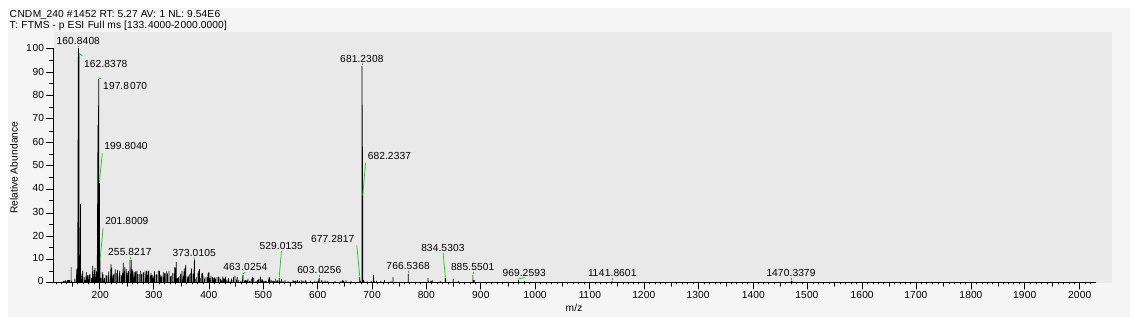

**2_4**

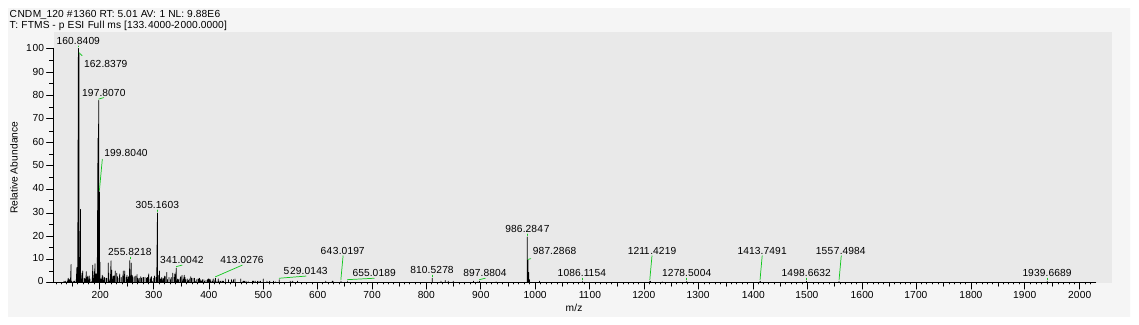

**Supplementary Figure 3.** MS1 spectra of CN-DM-861 and metabolites given in Supplementary Figure 2.

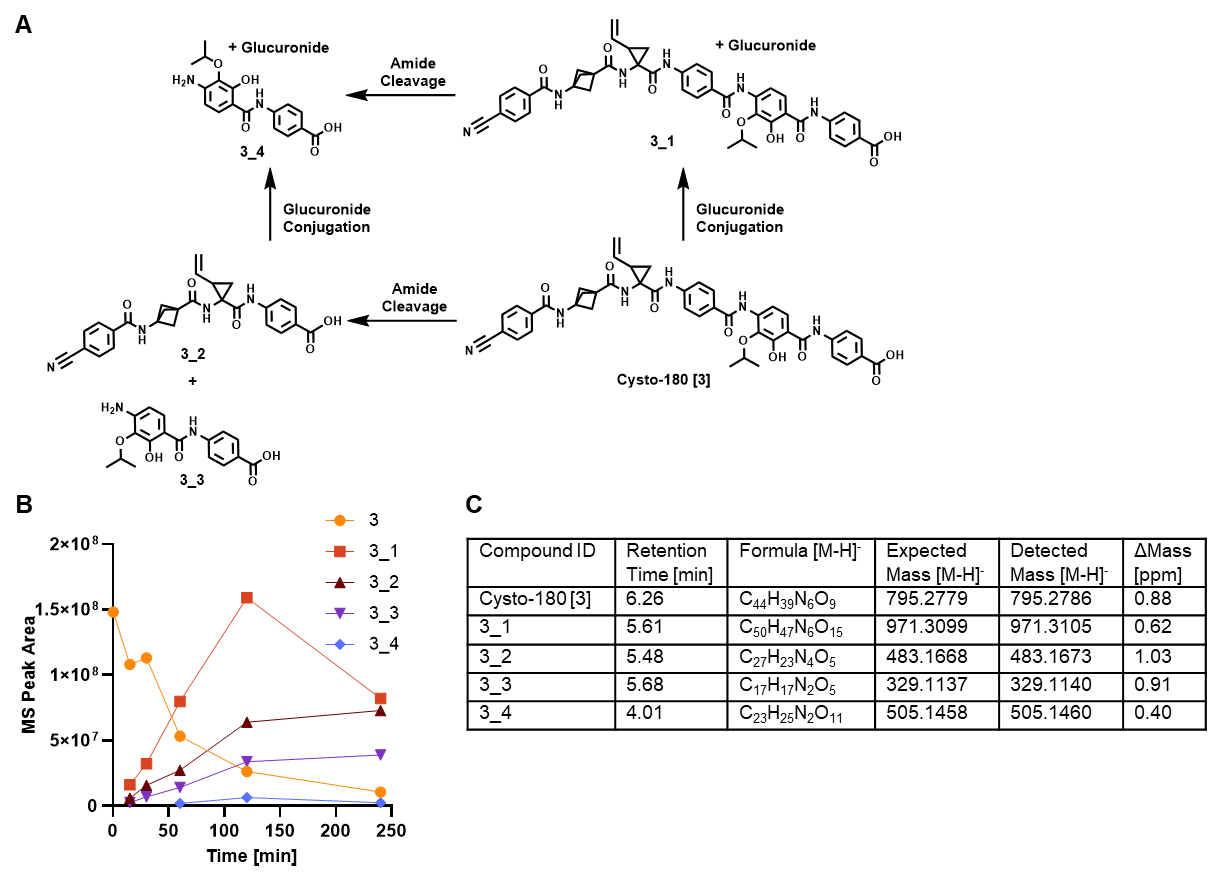

**Supplementary Figure 4.** MetID of Cysto-180. **(A)** MetID of Cysto-180 showed glucuronidation and amide bond hydrolysis as main metabolic degradation pathways in mouse hepatocytes. **(B)** Time-dependent analysis of mass signals for the respective compounds showed a correlation between parent compound degradation and increase of identified metabolites, where glucuronidation as well as amide cleavage between rings C and D of the parent compound seemed to be the most prominent paths. The exact position of the glucuronidation could not be clearly identified between the hydroxy and the carboxy group. **(C)** Corresponding chromatographic and MS data of identified metabolites. (See also Supplementary Figure 5)

**3**

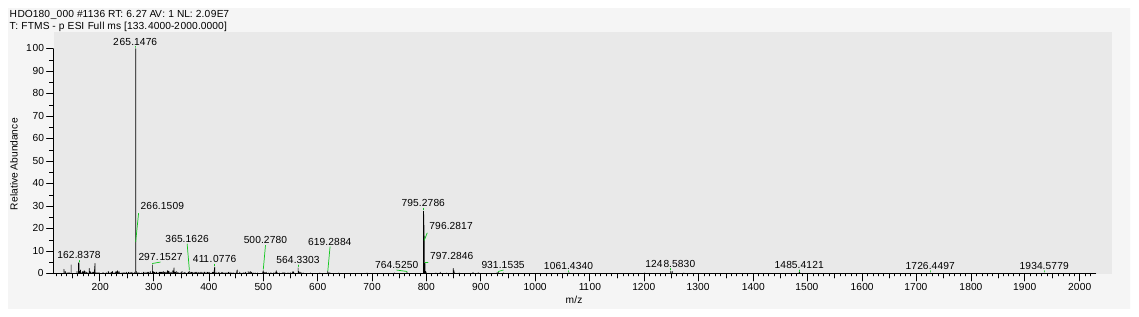

**3_1**

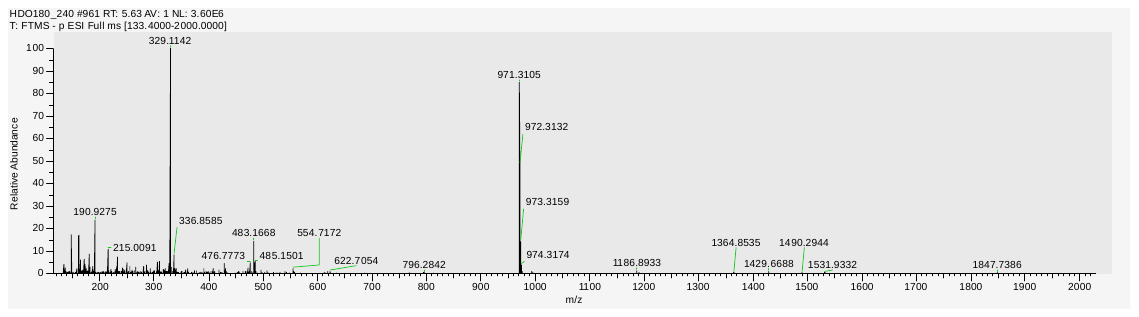

**3_2**

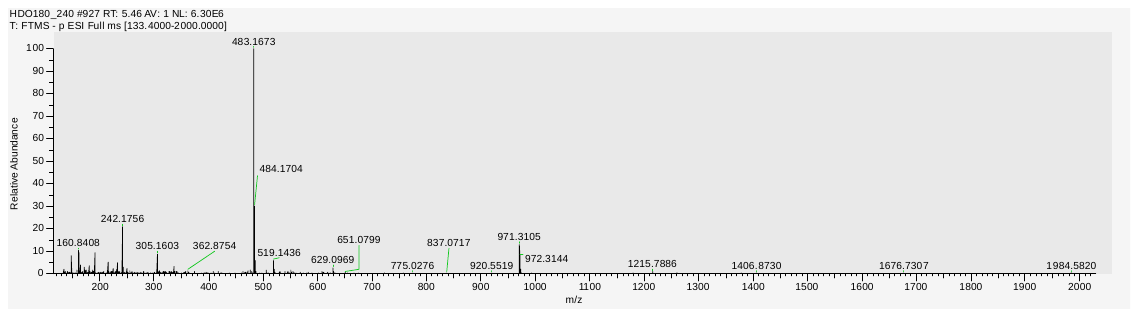

**3_3**

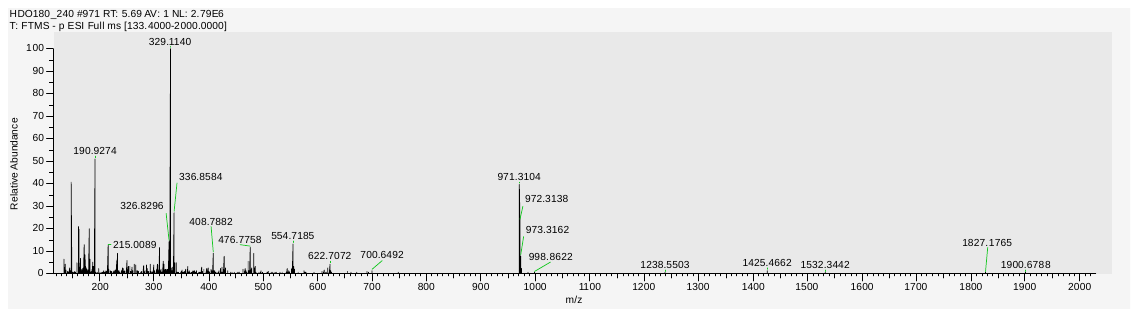

**3_4**

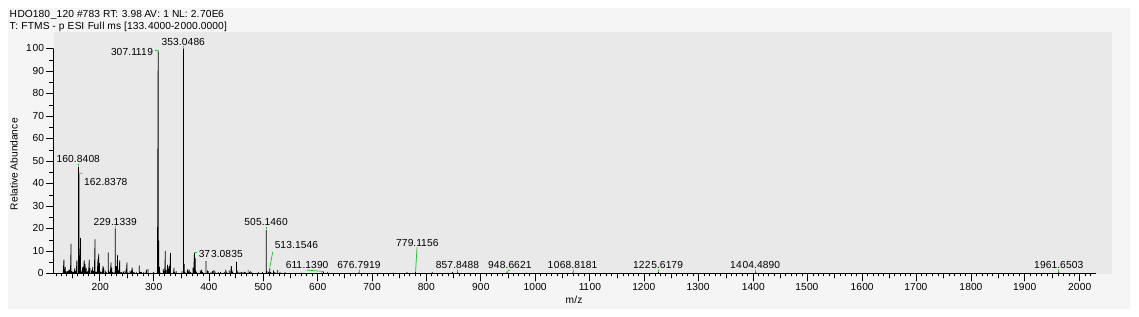

**Supplementary Figure 5.** MS1 spectra of Cysto-180 and metabolites given in Supplementary Figure 4.

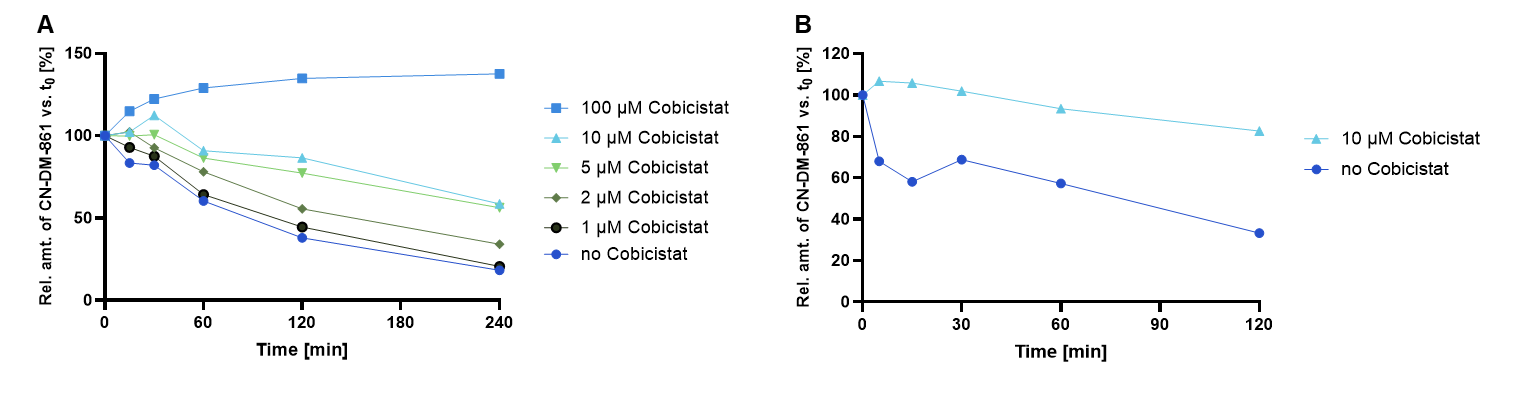

**Supplementary Figure 6.** Concentration-dependent increase of metabolic stability of CN-DM-861 via cobicistat co-treatment. **(A)** CN-DM-861 incubated with mouse hepatocytes in presence of cobicistat indicates stabilization due to OATP-inhibition. **(B)** In mouse liver microsomes, cobicistat shows a similar effect, showing a possible involvement of CYP3A inhibition.

Structures and synthesis of Cysto-354

General Methods

All non-aqueous reactions were carried out in dried glassware in dry solvents under inert conditions unless otherwise noted. Light sensitive reactions were carried out under light exclusion. Dry solvents (MeCN, DMF, Et_2_O) were taken from a MBraun solvent purification system. THF was freshly distilled over sodium (benzophenone as indicator). Petroleum ether was distilled (60°C). Et_3_N was freshly distilled over KOH. Other commercially available (dry) solvents were purchased from Merck or Acros Organics.

For reactions under microwave irradiation a CEM Discover S-Class was used with a power maximum of 300 W.

Chromatographic separations by flash chromatography were carried out on a Grace Reveleris X2 (Büchi) with FlashPure EcoFlex cartridges (Büchi) or conducted with the flash purification system Sepacore (Büchi) or Biotage SP using prepacked cartridges (puriFlash by Interchim or chromobond by Macherey‑Nagel). A Pure C-850 FlashPrep (Büchi) with FlashPure EcoFlex cartriges (Büchi) was utilized for reversed phase flash chromatography. For manual columns silica gel 60 0.04-0.063 mm; 230-400 mesh (Machery-Nagel) was used.

Purifications by high performance liquid chromatography (HPLC) were performed by a Thermo Scientific Dionex UltiMate 3000 system with a Phenomenex Luna C18 column (250 mm x 21.2 mm, 5 µm) column under basic (10 mM NH_4_HCO_3_) or acidic (0.1% formic acid or acetic acid) conditions. Alternatively, semi-preparative HPLC was performed by using a Waters Alliance 2695 HPLC-system with a 996 diode array detector (λ = 200-350 nm) and a Macherey-Nagel Nucleodur C18 ISIS column (5 µm, 250 mm, diameter = 8 mm). Mass detection was conducted with a Waters Quattro micro API mass spectrometer in negative ionization mode.

Thin-layer chromatography analytics were carried out on pre-coated silica gel 60 F_254_ plates (Merck) or on Macherey‑Nagel aluminium plates coated with silica gel 60 F245. The sample was detected by UV light at 254 nm or 366 nm. Non-UV-absorbent samples were stained by a cerium-ammonium-molybdate, potassium permanganate, ninhydrin, vanillin and anisaldehyde solution.

Bruker Advance-III HD 500 MHz and Bruker Advance-III HD 700 MHz spectrometer were used to measure NMR spectra. Alternatively, Bruker Ascend 600 MHz with Avance Neo console, Ultrashield 500 MHz with Avance-III HD console, Ascend 400 MHz with Avance- III console, Ascend 400 MHz with Avance-III HD console or Ultrashield 400 MHz with Avance-I console were used. The chemical shifts for ^1^H, ^13^C and ^19^F spectra are reported in ppm at a temperature of 300 K. ^19^F spectra lack an internal reference. One dimensional ^13^C were measured with ^1^H decoupling. Multiplicities are specified with following abbreviations: s = singlet, d = doublet, t = triplet, q = quartet, hept./sept. = septet, oct = octet, m = multiplet, br = broad signal and combinations thereof.

LCMS reaction controls were carried out by Agilent 1260 Infinity II LC connected to an Agilent 6130 (quadrupole MS) in ESI mode by a Phenomenex Gemini NX-C18 (50 mm x 2 mm, 3 µm) column. The gradient went from 0% – 100% acetonitrile to water with 0.1% formic acid in both solvents over three minutes at a flow rate of 1.5 ml/min.

High-resolution mass spectra were measured at a Bruker maXis HD or 4G spectrometer in positive or negative ESI mode or at a Micromass LCT with lock-spray unit and injection via loop modus in a Waters (Alliance 2695) HPLC device. Alternatively, a Micromass Q-TOF was used in combination with a Waters Aquity UPLC device. The ionization occurred through electron spray ionization. Calculated and found masses are reported.

The specific optical rotation [*α*] was measured with a polarimeter type 341 from Perkin‑Elmer at λ = 589.3 nm (sodium D line) in a 10 cm quartz cuvette. It is given in 10^-1^ cm^2^ g^-1^. The concentration c is given in 10 mg mL^-1^.

A Christ Alpha 1-4 LCSbasic was used for the lyophilization of the products after purification by HPLC.

Synthesis of Cysto-354

***tert*-Butyl 4-(2,2,2-trifluoroacetyl)benzoate (226)**

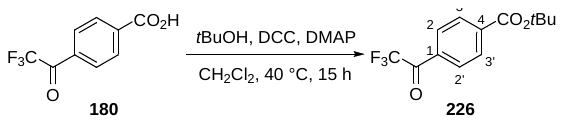

**Supplementary Figure 7.** Synthesis of *tert*-Butyl 4-(2,2,2-trifluoroacetyl)benzoate (**226**).

4‑(Trifluoroacetyl)benzoic acid (**180**) (1.00 g, 4.58 mmol, 1.00 eq.) was dissolved in CH_2_Cl_2_ (4.6 mL). Then, *tert*-butyl alcohol (1.29 mL, 13.8 mmol, 3.00 eq.), *N*,*N*‑dimethyl‑4‑amino-pyridine (448 mg, 3.67 mmol, 0.80 eq.), and *N*,*N*’‑dicyclohexylcarbodiimide (1.04 g, 5.04 mmol, 1.10 eq.) were added. After heating under refluxing conditions for 15 h, the reaction mixture was cooled to room temperature and filtered. The filtrate was washed with aq. 0.5 M HCl (5 mL), aq. sat. NaHCO_3_ (5 mL), and brine (5 mL), dried over MgSO_4_, filtered, and concentrated under reduced pressure. Flash column chromatography (CH_2_Cl_2_/PE = 3/7) afforded ester **226** (445 mg, 1.62 mmol, 35%) as a colorless resin.

The analytical data are in accordance with those reported in the literature.^[184]^

**^1^H-NMR (400 MHz, CDCl_3_):** δ = 8.15 ‑ 8.10 (m, 4 H, H‑2, H‑2‘, H‑3, H‑3‘), 1.62 (s, 9 H, CO_2_CMe_3_) ppm.

**R*_ƒ_*** (PE/EtOAc = 5/1): 0.46.

***tert*-Butyl 4-(2,2,2-trifluoro-1-(((methylsulfonyl)oxy)imino)ethyl)benzoate (227)**

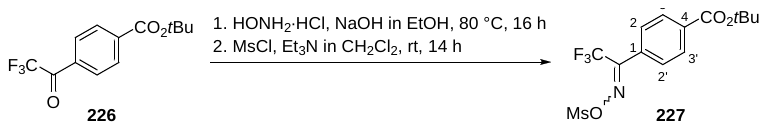

**Supplementary Figure 8.** Synthesis of *tert*-Butyl 4-(2,2,2-trifluoro-1-(((methylsulfonyl)oxy)imino)ethyl)benzoate (**227**).

Ketone **226** (460 mg, 1.68 mmol, 1.00 eq.) in EtOH (2.5 mL) was added dropwise to a refluxing mixture of hydroxylamine hydrochloride (525 mg, 7.55 mmol, 4.50 eq.) and sodium hydroxide (302 mg, 7.55 mmol, 4.50 eq.) in EtOH (3.5 mL). After heating under refluxing conditions for 16 h, the mixture was cooled to room temperature and EtOAc (200 mL) was added. The solution was washed with aq. 0.1 M HCl (3x 50 mL) and brine (50 mL), dried over MgSO_4_, filtered, and concentrated under reduced pressure. The residue was dissolved in CH_2_Cl_2_ (5.8 mL). At 0 °C, methanesulfonyl chloride (0.13 mL, 1.68 mmol, 1.00 eq.) was added followed by dropwise addition of triethylamine (0.37 mL, 2.68 mmol, 1.60 eq.) in CH_2_Cl_2_ (1.2 mL) over 15 min and stirring of additional 15 min. After stirring at room temperature for 14 h, the mixture was concentrated *in vacuo*. Then, the residue was dissolved in water (100 mL) and extracted with Et_2_O (3x 100 mL). The combined organic phases were dried over MgSO_4_, filtered, and concentrated under reduced pressure. Flash column chromatography (100% CHCl_3_) afforded product **227** (446 mg, 1.21 mmol, 72% o2s) as a yellow oil.

The analytical data are in accordance with those reported in the literature.^[184]^

**^1^H-NMR (400 MHz, CDCl_3_):** δ = 8.13 ‑ 8.08 (m, 2 H, H‑3, H‑3‘), 7.66 ‑ 7.52 (m, 2 H, H‑2, H‑2‘), 3.26 (s, 3 H, Ms), 1.61 (s, 9 H, CO_2_CMe_3_) ppm.

**R*_ƒ_*** (100% CHCl_3_): 0.28.

***tert*-Butyl 4-(3-(trifluoromethyl)diaziridin-3-yl)benzoate (228)**

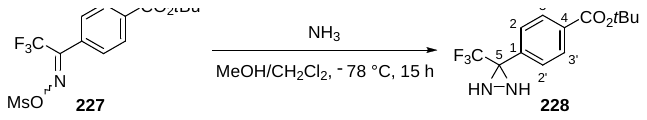

**Supplementary Figure 9.** Synthesis of *tert*-Butyl 4-(3-(trifluoromethyl)diaziridin-3-yl)benzoate (**228**).

At −78 °C, 7 M NH_3_/MeOH (0.9 mL) was added to a solution of compound **227** (250 mg, 0.68 mmol, 1.00 eq.) in CH_2_Cl_2_ (0.11 mL). After stirring at −78 °C for 15 h, the mixture was warmed up to room temperature and concentrated *in vacuo*. The residue was diluted in CH_2_Cl_2_ (30 mL), washed with water (10 mL) and brine (10 mL), dried over MgSO_4_, filtered, and concentrated under reduced pressure. Purification by flash column chromatography (100% CH_2_Cl_2_) afforded diaziridine **228** (108 mg, 0.38 mmol, 55%) as a colorless resin.

The analytical data are in accordance with those reported in the literature.^[184]^

**^1^H-NMR (400 MHz, CDCl_3_):** δ = 8.05 ‑ 8.03 (m, 2 H, H‑3, H‑3‘), 7.69 ‑ 7.67 (m, 2 H, H‑2, H‑2‘), 2.84 (bs, 1 H, NH), 2.24 (bs, 1 H, NH), 1.60 (s, 9 H, CO_2_CMe_3_) ppm.

**R*_ƒ_*** (100% CH_2_Cl_2_): 0.15.

**4-(3-(Trifluoromethyl)-3*H*-diazirin-3-yl)benzoic acid (181)**

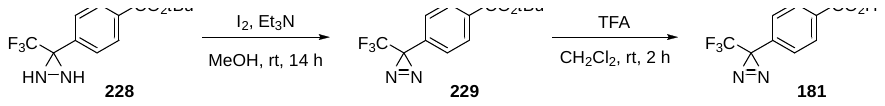

**Supplementary Figure 10.** Synthesis of 4-(3-(Trifluoromethyl)-3*H*-diazirin-3-yl)benzoic acid (**181**).

Iodine (275 mg, 1.08 mmol, 3.00 eq.) was added portion wise to a solution of diaziridine **228** (104 mg, 0.36 mmol, 1.00 eq.) and triethylamine (0.15 mL, 1.08 mmol, 3.00 eq.) in MeOH (5.2 mL). After stirring for 14 h, the mixture was diluted in CH_2_Cl_2_ (50 mL) and washed with 1 M aq. NaOH (50 mL), water (50 mL), and brine (50 mL), dried over MgSO_4_, filtered, and concentrated under reduced pressure. Flash column chromatography (PE/EtOAc = 99/1) afforded diazirine **229** (103 mg, 0.36 mmol, >99%) which was immediately used in the next step.

Compound **229** was synthesized following the protocol of Sakurai *et al*.^[184]^

**R*_ƒ_*** (PE/EtOAc = 19/1): 0.55.

Trifluoroacetic acid (0.1 mL) was added to a solution of diazirine **229** (103 mg, 0.36 mmol, 1.00 eq.) in CH_2_Cl_2_ (1.1 mL). After stirring for 2 h, the solvent was removed under reduced pressure. The residue was diluted in CH_2_Cl_2_ (25 mL) and was treated with sat. aq. NaHCO_3_ (25 mL). The phases were separated and the aqueous phase was acidified to pH = 3 with 1 M aq. HCl. The aqueous phase was extracted with CH_2_Cl_2_ (50 mL) and the organic phase was dried over MgSO_4_, filtered, and concentrated under reduced pressure to furnish carboxylic acid **181** (82 mg, 0.36 mmol, >99%) as a colorless semisolid which was immediately used in the next step without further purification.

Compound **181** was synthesized following the protocol of Sakurai *et al*.^4^

**R*_ƒ_*** (PE/EtOAc = 19/1): 0.04.

***tert*-Butyl 4-(4-(3-(trifluoromethyl)-3*H*-diazirin-3-yl)benzamido)benzoate (230)**

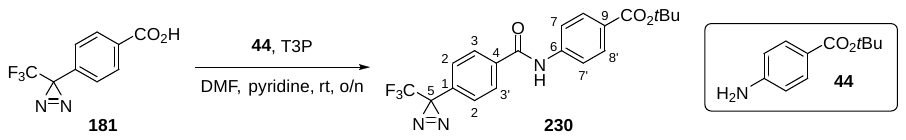

**Supplementary Figure 11.** Synthesis of *tert*-Butyl 4-(4-(3-(trifluoromethyl)-3*H*-diazirin-3-yl)benzamido)benzoate (**230**).

Pyridine (0.1 mL) was added to a stirred suspension of amine **44** (66 mg, 344 µmol, 1.00 eq.) and carboxylic acid **181** (87 mg, 378 µmol, 1.10 eq.) in DMF (0.2 mL) at 0 °C. Then, T3P (0.48 mL, 50% in DMF, 756 µmol, 2.00 eq.) was added dropwise and the reaction mixture was stirred overnight while slowly warming up to room temperature. After addition of aq. HCl (1 mL, 1 M) at 0 °C, the suspension was stirred at room temperature for 2 h. The resulting precipitate was filtered, washed with an excess of water (3x), transferred to a clean flask, and stirring was continued for 1 h in water (5 mL). Once again, the precipitate was filtered and washed with an excess of water (3x). Drying *in vacuo* overnight afforded amide **230** (138 mg, 0.34 mmol, 99%) as a colorless foam.

**^1^H-NMR (400 MHz, DMSO-*d*_6_):** δ = 10.66 (s, 1 H, CONH), 8.09 ‑ 8.06 (m, 2 H, H‑3, H‑3’), 7.92 ‑ 7.88 (m, 4 H, H‑7, H‑7’, H‑8, H‑8’), 7.47 ‑ 7.45 (m, 2 H, H‑2, H‑2’), 1.55 (s, 9 H, CO_2_CMe_3_) ppm.

**^13^C-NMR (100 MHz,** **DMSO-*d*_6_):** δ = 164.9 (q, CONH), 164.6 (q, *C*O_2_CMe_3_), 143.0 (q, C‑6), 136.2 (q, C‑4), 130.8 (q, C‑1), 130.0 (t, C‑8, C‑8’), 128.7 (t, C‑3, C‑3’), 126.5 (t, quartet, *J* = 1.2 Hz, C‑2, C‑2‘), 126.3 (q, C‑9), 121.8 (q, quartet, *J* = 274.7 Hz, CF_3_), 119.5 (t, C‑7, C‑7’), 80.4 (q, CO_2_*C*Me_3_), 28.1 (q, quartet, *J* = 40.1 Hz, C‑5), 27.8 (p, CO_2_C*Me_3_*) ppm.

**^19^F-NMR (376 MHz,** **DMSO-*d*_6_):** δ = −64.66 ppm.

**HRMS (ESI):** m/z calculated for C_20_H_18_N_3_O_3_F_3_Na [M+Na]^+^: 428.1198; found: 428.1191.

**R*_ƒ_*** (PE/EtOAc = 5/1): 0.37.

**4-(4-(3-(Trifluoromethyl)-3*H*-diazirin-3-yl)benzamido)benzoic acid (182)**

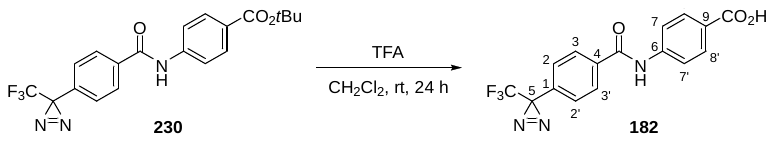

**Supplementary Figure 12.** Synthesis of 4-(4-(3-(Trifluoromethyl)-3*H*-diazirin-3-yl)benzamido)benzoic acid (**182**).

Trifluoroacetic acid (0.07 mL) was added to a solution of ester **230** (110 mg, 271 µmol, 1.00 eq.) in CH_2_Cl_2_ (0.82 mL). After stirring for 24 h, all volatiles were removed under reduced pressure. After drying *in vacuo* overnight, carboxylic acid **182** (94 mg, 0.27 mmol, >99%) was obtained as a colorless foam.

The analytical data are in accordance with those reported in the literature.^[205]^

**^1^H-NMR (500 MHz, DMSO-*d*_6_):** δ = 12.74 (bs, 1 H, CO_2_H), 10.66 (s, 1 H, CONH), 8.09 ‑ 8.06 (m, 2 H, H‑3, H‑3’), 7.96 ‑ 7.93 (m, 2 H, H‑8, H‑8’), 7.91 ‑ 7.88 (m, 2 H, H‑7, H‑7’), 7.47 ‑ 7.46 (m, 2 H, H‑2, H‑2’) ppm.

**^13^C-NMR (125 MHz,** **DMSO-*d*_6_):** δ = 166.9 (q, CO_2_H), 164.9 (CONH), 143.0 (q, C‑6), 136.2 (q, C‑4), 130.8 (q, C‑1), 130.3 (t, C‑8, C‑8’), 128.7 (t, C‑3, C‑3‘), 126.6 (t, quartet, *J* = 1.2 Hz, C‑2, C‑2‘), 125.8 (q, C‑9), 121.8 (q, quartet, *J* = 274.9 Hz, CF_3_), 119.6 (t, C‑7, C‑7‘), 28.1 (q, quartet, *J* = 40.2 Hz, C‑5) ppm.

**^19^F-NMR (376 MHz,** **DMSO-*d*_6_):** δ = −64.67 ppm.

**HRMS (ESI):** m/z calculated for C_16_H_9_N_3_O_3_F_3_ [M−H]^−^: 348.0596; found: 348.0586.

**R*_ƒ_*** (PE/EtOAc =5/1): 0.04.

***tert*-Butyl (*S*)-4-(2-(allyloxy)-4-(4-(2-((*tert*-butoxycarbonyl)amino)pent-4‑ynamido)benzamido)-3-isopropoxybenzamido)benzoate (231)**

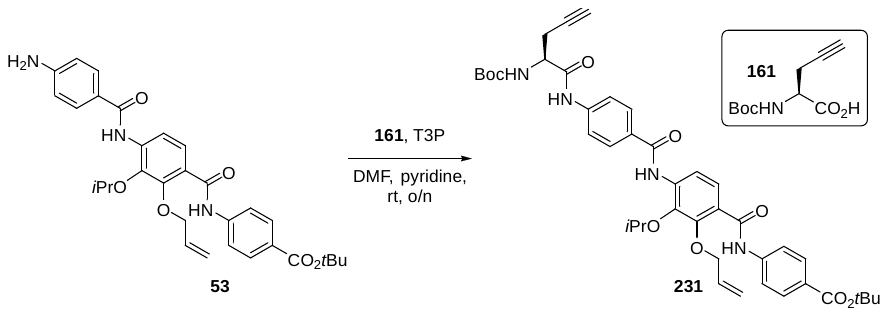

**Supplementary Figure 13.** Synthesis of *tert*-Butyl (*S*)-4-(2-(allyloxy)-4-(4-(2-((*tert*-butoxycarbonyl)amino)pent-4 ynamido)benzamido)-3-isopropoxybenzamido)benzoate (**231**).

Pyridine (1.19 mL) was added to a stirred suspension of amine **53** (2.47 g, 4.48 mmol, 1.00 eq.) and carboxylic acid **161** (1.05 g, 4.92 mmol, 1.10 eq.) in DMF (2.4 mL) at 0 °C. Then, T3P (5.7 mL, 50% in DMF, 8.96 mmol, 2.00 eq.) was added dropwise and the reaction mixture was stirred overnight while slowly warming up to room temperature. After addition of aq. 0.5 M HCl (100 mL) at 0 °C, the suspension was stirred at room temperature for 2 h. The resulting precipitate was filtered, washed with an excess of water (3x), transferred to a clean flask, and stirring was continued for 1 h in water (100 mL). Once again, the precipitate was filtered and washed with an excess of water (3x) to afford crude amide **231** (3.32 g, 4.48 mmol, >99%) as an orange foam which was used in the next step without further purification.

**^1^H-NMR (400 MHz, DMSO-*d*_6_):** δ = 10.53 (s, 1 H, CONH), 10.43 (s, 1 H, CONH), 9.52 (s, 1 H, CONH), 7.99 ‑ 7.95 (m, 3 H, Ar‑*H*), 7.90 ‑ 7.88 (m, 2 H, Ar‑*H*), 7.84 ‑ 7.81 (m, 2 H, Ar‑*H*), 7.79 ‑ 7.77 (m, 2 H, Ar‑*H*), 7.41 (d, *J* = 8.4 Hz, 1 H, Ar‑*H*), 7.22 (d, *J* = 7.9 Hz, 1 H, N*H*Boc), 6.02 (ddt, *J* = 17.2, 10.6, 5.4 Hz, 1 H, -CH=), 5.37 (pseudo dq, *J* = 17.2, 1.6 Hz, 1 H, =CH_2_ *trans*), 5.20 (pseudo dq, *J* = 10.4, 1.4 Hz, 1 H, =CH_2_ *cis*), 4.61 (d, *J* = 5.5 Hz, 2 H, CH_2_ *allylic*), 4.50 (septet, *J* = 6.2 Hz, 1 H, C*H*Me_2_), 4.30 (q, *J* = 7.3 Hz, 1 H, *H*CNHBoc), 2.91 (t, *J* = 2.4 Hz, 1 H, -C≡CH), 2.61 ‑ 2.52 (m, 2 H, CH_2_ *propargylic*), 1.55 (s, 9 H, CO_2_CMe_3_), 1.40 (s, 9 H, HNCO_2_C*Me_3_*), 1.26 (d, *J* = 6.1 Hz, 6 H, CH*Me_2_*) ppm.

**^13^C-NMR (100 MHz,** **DMSO-*d*_6_):** δ = 169.9 (q, C=O), 164.6 (q, C=O), 164.5 (q, C=O), 164.3 (q, C=O), 155.2 (HN*C*O_2_CMe_3_), 149.5 (q, Ar‑C), 143.0 (q, Ar‑C), 142.5 (q, Ar‑C), 142.1 (q, Ar‑C), 138.5 (t, Ar‑C), 135.6 (q, Ar‑C), 133.6 (t, -CH=), 130.1 (t, Ar‑C), 128.5 (q, Ar‑C), 128.5 (t, Ar‑C), 127.1 (q, Ar‑C), 126.0 (q, Ar‑C), 123.6 (t, Ar‑C), 118.9 (t, Ar‑C), 118.8 (t, Ar‑C), 117.8 (s, =CH_2_), 80.5 (q, ‑*C*≡CH), 80.3 (q, CO_2_*C*Me_3_), 78.4 (q, HNCO_2_*C*Me_3_), 76.2 (t, *C*HMe_2_), 74.3 (s, CH_2_ *allylic*), 73.2 (t, ‑C≡*C*H), 54.1 (t, H*C*NHBoc), 28.2 (p, HNCO_2_C*Me_3_*), 27.8 (p, CO_2_C*Me_3_*), 22.3 (p, CH*Me_2_*), 21.7 (s, CH_2_ *propargylic*) ppm.

**HRMS (ESI):** m/z calculated for C_41_H_48_N_4_O_9_Na [M+Na]^+^: 763.3319; found: 763.3317.

$\boldsymbol{[\alpha]}_{\mathbf{D}}^{\boldsymbol{24.0}}$ = −11.4° (*c* 0.8, MeCN).

**R*_ƒ_*** (PE/EtOAc = 1/1): 0.48.

***tert*-Butyl (*S*)-4-(2-(allyloxy)-4-(4-(2-aminopent-4-ynamido)benzamido)-3‑isopropoxybenzamido)benzoate (162)**

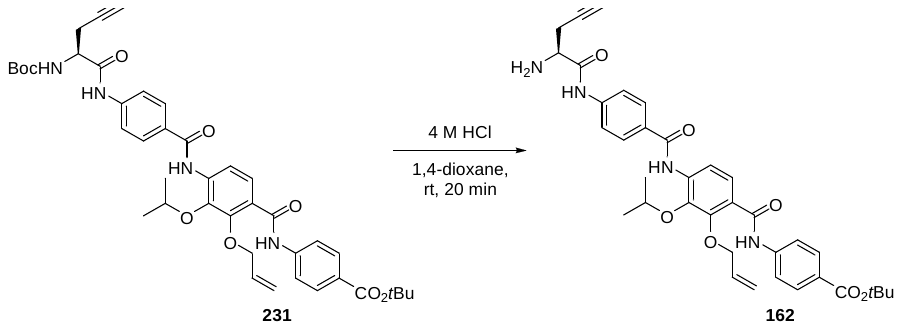

**Supplementary Figure 14.** Synthesis of *tert*-Butyl (*S*)-4-(2-(allyloxy)-4-(4-(2-aminopent-4-ynamido)benzamido)-3 isopropoxybenzamido)benzoate (**162**).

HCl (2.8 mL, 4 M in 1,4-dioxane, 10.8 mmol, 100 eq.) was added to amide **231** (80 mg, 108 µmol, 1.00 eq.) at 0 °C. The solution was warmed up to room temperature (over a period of 5 min) and stirred for further 15 min. Then, the reaction mixture was poured into a vigorously stirred mixture of EtOAc (50 mL) and sat. aq. NaHCO_3_ (50 mL). The phases were separated, the aqueous phase was extracted with EtOAc (3x 50 mL), the combined organic phases were washed with brine, dried over MgSO_4_, filtered, and concentrated *in vacuo*. Flash column chromatography (washing with PE/EtOAc = 1/5, then eluting with 100% EtOAc) afforded amine **162** (65 mg, 0.10 mmol, 93%) as an orange foam.

The analytical data are in accordance with those reported in the literature.^5^

**^1^H-NMR (400 MHz, DMSO-*d*_6_):** δ = 10.53 (s, 1 H, CONH), 9.52 (s, 1 H, CONH), 7.99 ‑ 7.97 (m, 2 H, Ar‑*H*), 7.90 ‑ 7.88 (m, 2 H, Ar‑*H*), 7.84 ‑ 7.79 (m, 5 H, Ar‑*H*), 7.41 (d, *J* = 8.5 Hz, 1 H, Ar‑*H*), 6.02 (ddt, *J* = 17.2, 10.8, 5.4 Hz, 1 H, -CH=), 5.37 (pseudo dq, *J* = 17.2, 1.6 Hz, 1 H, =CH_2_ *trans*), 5.20 (pseudo dq, *J* = 10.5, 1.3 Hz, 1 H, =CH_2_ *cis*), 4.64 ‑ 4.58 (m, 4 H, CH_2_ *allylic*, NH_2_), 4.50 (septet, *J* = 6.1 Hz, 1 H, C*H*Me_2_), 3.55 (t, *J* = 6.3 Hz, 1 H, *H*CNH_2_), 2.87 (t, *J* = 2.6 Hz, 1 H, -C≡CH), 2.59 ‑ 2.53 (m, 1 H, CH_2_ *propargylic*), 2.52 ‑ 2.46 (m, 1 H, CH_2_ *propargylic*), 1.55 (s, 9 H, CO_2_CMe_3_), 1.26 (d, *J* = 6.2 Hz, 6 H, CH*Me_2_*) ppm.

**^13^C-NMR (100 MHz,** **DMSO-*d*_6_):** δ = 172.8 (q, C=O), 164.6 (q, C=O), 164.5 (q, C=O), 164.3 (q, C=O), 149.5 (q, Ar‑C), 143.0 (q, Ar‑C), 142.6 (q, Ar‑C), 142.1 (q, Ar‑C), 135.6 (q, Ar‑C), 133.7 (t, -CH=), 130.1 (t, Ar‑C), 128.4 (t, Ar‑C), 128.4 (q, Ar‑C), 127.1 (q, Ar‑C), 126.0 (q, Ar‑C), 123.6 (t, Ar‑C), 119.0 (t, Ar‑C), 118.8 (t, Ar‑C), 118.7 (t, Ar‑C), 117.8 (s, =CH_2_), 81.2 (q, -*C*≡CH), 80.3 (q, CO_2_*C*Me_3_), 76.2 (t, *C*HMe_2_), 74.3 (s, CH_2_ *allylic*), 73.1 (t, -C≡*C*H), 54.6 (t, HCNH_2_), 27.9 (p, CO_2_C*Me_3_*), 24.8 (s, CH_2_ *propargylic*), 22.3 (p, CH*Me_2_*) ppm.

**HRMS (ESI):** m/z calculated for C_36_H_41_N_4_O_7_ [M+H]^+^: 641.2975; found: 641.2955.

${\boldsymbol{[}\boldsymbol{\alpha}\boldsymbol{]}}_{\mathbf{D}}^{\boldsymbol{23}\boldsymbol{.}\boldsymbol{6}}$ = −13.0° (*c*, 0.3, DMSO-*d*_6_).

**R*_ƒ_*** (PE/EtOAc =1/5): 0.15.

***tert*-Butyl (*S*)-4-(2-(allyloxy)-3-isopropoxy-4-(4-(2-(4-(4-(3-(trifluoromethyl)-3*H*-diazirin-3‑yl)benzamido)benzamido)pent-4-ynamido)benzamido)benzamido)benzoate (183)**

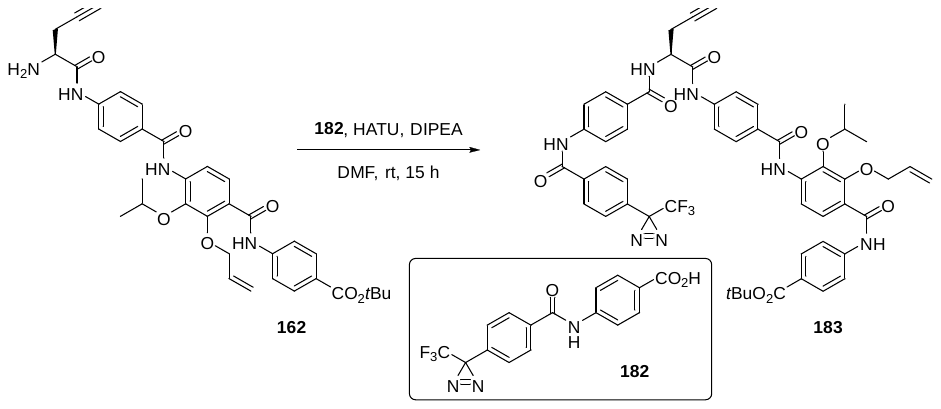

**Supplementary Figure 15.** Synthesis of *tert*-Butyl (*S*)-4-(2-(allyloxy)-3-isopropoxy-4-(4-(2-(4-(4-(3-(trifluoromethyl)-3*H*-diazirin-3 yl)benzamido)benzamido)pent-4-ynamido)benzamido)benzamido)benzoate (**183**).

DIPEA (0.21 mL, 1.22 mmol, 13.0 eq.) was added dropwise to a solution of carboxylic acid **182** (82 mg, 234 µmol, 2.50 eq.) and HATU (89 mg, 234 µmol, 2.50 eq.) in DMF (2.2 mL). After 5 min, the resulting mixture was slowly transferred to a solution of amine **162** (60 mg, 94 µmol, 1.00 eq.) in DMF (1.3 mL) at 0 °C. Then, the reaction mixture was stirred for 15 h while slowly warming up to room temperature. Next, the mixture was diluted with EtOAc (10 mL), washed with aq. HCl (7 mL, 0.1 M), aq. NaCl (5x 20 mL, 5% sat.), dried over MgSO_4_, filtered, and concentrated under reduced pressure in the presence of silica gel. Flash column chromatography (PE/EtOAc = 5/1, then 1/1) afforded amide **183** (91 mg, 94 µmol, >99%) as a yellow foam which was of enough purity to be used in the next step.

**^1^H-NMR (500 MHz, DMSO-*d*_6_):** δ = 10.67 (s, 1 H, CONH), 10.64 (s, 1 H, CONH), 10.53 (s, 1 H, CONH), 9.53 (s, 1 H, CONH), 8.77 (d, *J* = 7.5 Hz, 1 H, CONH), 8.10 ‑ 8.06 (m, 4 H, Ar‑*H*), 7.95 ‑ 7.93 (m, 4 H, Ar‑*H*), 7.90 ‑ 7.88 (m, 5 H, Ar‑*H*), 7.84 ‑ 7.80 (m, 2 H, Ar‑*H*), 7.47 ‑ 7.45 (m, 3 H, Ar‑*H*), 6.02 (ddt, *J* = 17.2, 10.8, 5.4 Hz, 1 H, -CH=), 5.37 (pseudo dq, *J* = 17.3, 1.6 Hz, 1 H, =CH_2_ *trans*), 5.20 (pseudo dq, *J* = 10.6, 1.5 Hz, 1 H, =CH_2_ *cis*), 4.80 (q, *J* = 7.4 Hz, 1 H, C*H*NH), 4.61 (d, *J* = 5.5 Hz, 2 H, CH_2_ *allylic*), 4.49 (septet, *J* = 6.1 Hz, 1 H, C*H*Me_2_), 2.93 (t, *J* = 2.6 Hz, 1 H, -C≡CH), 2.84 ‑ 2.73 (m, 2 H, CH_2_ *propargylic*), 1.55 (s, 9 H, CO_2_CMe_3_), 1.26 (d, *J* = 6.1 Hz, 6 H, CH*Me_2_*) ppm.

**^13^C-NMR (125 MHz,** **DMSO-*d*_6_):** δ = 169.7 (q, C=O), 166.9 (q, C=O), 166.0 (q, C=O), 164.9 (q, C=O), 164.6 (q, C=O), 164.3 (q, C=O), 149.5 (Ar‑C), 143.0 (Ar‑C), 142.6 (Ar‑C), 141.8 (Ar‑C), 136.2 (Ar‑C), 135.6 (Ar‑C), 133.7 (t, -CH=), 130.8 (Ar‑C), 130.2 (Ar‑C), 130.1 (Ar‑C), 128.8 (Ar‑C), 128.7 (Ar‑C), 128.7 (Ar‑C), 128.4 (Ar‑C), 128.4 (Ar‑C), 127.1 (Ar‑C), 126.5 (Ar‑C), 126.0 (Ar‑C), 123.6 (Ar‑C), 121.8 (q, quartet, *J* = 274.6 Hz, CF_3_), 119.6 (Ar‑C), 119.5 (Ar‑C), 118.9 (Ar‑C), 118.8 (Ar‑C), 117.8 (s, =CH_2_), 80.5 (q, -*C*≡CH), 80.3 (q, CO_2_*C*Me_3_), 76.3 (t, *C*HMe_2_), 74.3 (s, CH_2_ *allylic*), 73.2 (t, -C≡*C*H), 53.5 (t, CHNH), 28.1 (q, quartet, *J* = 40.1 Hz, *C*-CF_3_), 27.9 (p, CO_2_C*Me_3_*), 24.1 (s, CH_2_ *propargylic*), 22.3 (p, CH*Me_2_*) ppm.

**^19^F-NMR (376 MHz,** **DMSO-*d*_6_):** δ = −64.67 ppm.

**HRMS (ESI):** m/z calculated for C_52_H_49_N_7_O_9_F_3_ [M+H]^+^: 972.3544; found: 972.3550.

${\boldsymbol{[}\boldsymbol{\alpha}\boldsymbol{]}}_{\mathbf{D}}^{\boldsymbol{24}\boldsymbol{.}\boldsymbol{0}}$ = +14.5° (*c* 0.2, DMSO-*d*_6_).

**R*_ƒ_*** (PE/EtOAc =1/5): 0.64.

***tert*-Butyl (*S*)-4-(2-hydroxy-3-isopropoxy-4-(4-(2-(4-(4-(3-(trifluoromethyl)-3*H*-diazirin-3‑yl)benzamido)benzamido)pent-4-ynamido)benzamido)benzamido)benzoate (232)**

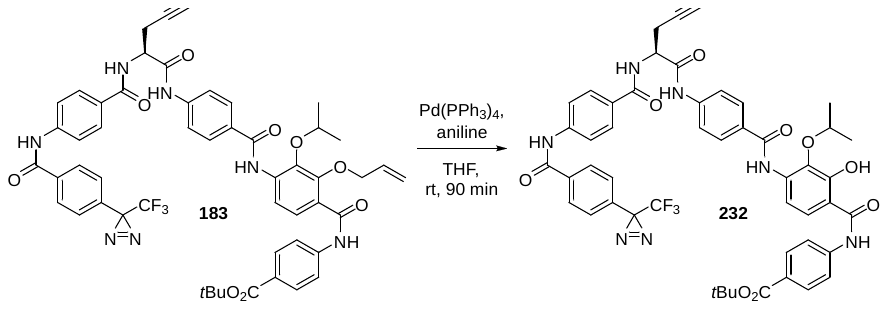

**Supplementary Figure 16.** Synthesis of *tert*-Butyl (*S*)-4-(2-hydroxy-3-isopropoxy-4-(4-(2-(4-(4-(3-(trifluoromethyl)-3*H*-diazirin-3 yl)benzamido)benzamido)pent-4-ynamido)benzamido)benzamido)benzoate (**232**).

Tetrakis(triphenylphosphine)palladium(0) (12 mg, 10 µmol, 0.10 eq.) was added in one portion to a stirred solution of ether **183** (100 mg, 103 µmol, 1.00 eq.) and aniline (30 µL, 340 µmol, 3.30 eq.) in THF (4.5 mL). After 90 min, the mixture was concentrated in the presence of silica gel *in vacuo*. Flash column chromatography (washing with PE/EtOAc = 5/1, then eluting with PE/EtOAc = 1/1) afforded phenol **232** (26 mg, 28 µmol, 28%) as a yellow foam.

**^1^H-NMR (500 MHz, DMSO-*d*_6_):** δ = 12.28 (bs, 1 H, OH), 10.61 (s, 1 H, CONH), 10.58 (s, 1 H, CONH), 10.16 (s, 1 H, CONH), 9.39 (bs, 1 H, CONH), 8.76 (d, *J* = 7.5 Hz, 1 H, CONH), 8.10 ‑ 8.07 (m, 2 H, Ar‑*H*), 8.00 ‑ 7.98 (m, 1 H, Ar‑*H*), 7.97 ‑ 7.93 (m, 4 H, Ar‑*H*), 7.91 ‑ 7.90 (m, 3 H, Ar‑*H*), 7.86 ‑ 7.74 (m, 5 H, Ar‑*H*), 7.48 ‑ 7.46 (m, 2 H, Ar‑*H*), 7.37 ‑ 7.33 (m, 1 H, Ar‑*H*), 4.80 (q, *J* = 7.5 Hz, 1 H, C*H*NH), 4.55 (bs, 1 H, C*H*Me_2_), 2.94 (t, *J* = 2.6 Hz, 1 H, -C≡CH), 2.84 ‑ 2.72 (m, 2 H, CH_2_ *propargylic*), 1.55 (s, 9 H, CO_2_CMe_3_), 1.26 (d, *J* = 6.2 Hz, 6 H, CH*Me_2_*) ppm.

**^13^C-NMR (125 MHz,** **DMSO-*d*_6_):** δ = 169.7 (q, C=O), 166.0 (q, C=O), 164,8 (q, C=O), 164.8 (q, C=O), 164.8 (q, C=O), 164.6 (q, C=O), 154.1 (q, COH; not visible in ^13^C but in HMBC), 142.2 (Ar‑C), 141.9 (Ar‑C), 141.8 (Ar‑C), 139.2 (Ar‑C),136.2 (Ar‑C),130.8 (Ar‑C), 129.9 (Ar‑C), 128.7 (Ar‑C), 128.7 (Ar‑C), 128.6 (Ar‑C), 128.5 (Ar‑C), 128.4 (Ar‑C), 128.3 (Ar‑C), 126.6 (Ar‑C), 126.5 (Ar‑C), 123.5 (Ar‑C), 121.8 (q, quartet, *J* = 274.8 Hz, CF_3_), 120.3 (Ar‑C), 119.5 (Ar‑C), 119.4 (Ar‑C), 119.4 (Ar‑C), 119.0 (Ar‑C), 80.6 (q, CO_2_*C*Me_3_), 80.5 (q, -*C*≡CH), 80.3 (t, *C*HMe_2_), 73.2 (t, -C≡*C*H), 53.5 (t, CHNH), 28.1 (q, quartet, *J* = 40.1 Hz, 1 H, *C*-CF_3_), 27.8 (p, CO_2_C*Me_3_*), 24.1 (CH_2_ *propargylic*), 22.3 (p, CH*Me_2_*) ppm.

**^19^F-NMR (376 MHz,** **DMSO-*d*_6_):** δ = −64.66 ppm.

**HRMS (ESI):** m/z calculated for C_49_H_43_N_7_O_9_F_3_ [M−H]^−^: 930.3074; found: 930.3071.

${\boldsymbol{[}\boldsymbol{\alpha}\boldsymbol{]}}_{\mathbf{D}}^{\boldsymbol{24}\boldsymbol{.}\boldsymbol{3}}$ = +25.0° (*c*, 0.1, DMSO-*d*_6_).

**R*_ƒ_*** (PE/EtOAc =1/5): 0.44.

**(*S*)-4-(2-Hydroxy-3-isopropoxy-4-(4-(2-(4-(4-(3-(trifluoromethyl)-3*H*-diazirin-3‑yl)benzamido)benzamido)pent-4-ynamido)benzamido)benzamido)benzoic acid (184)**

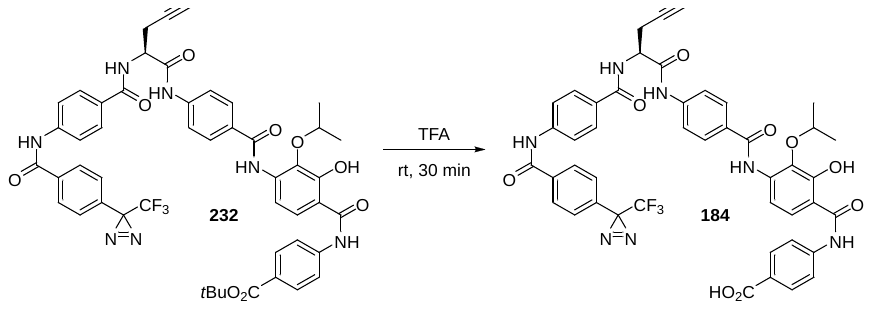

**Supplementary Figure 17.** Synthesis of (*S*)-4-(2-Hydroxy-3-isopropoxy-4-(4-(2-(4-(4-(3-(trifluoromethyl)-3*H*-diazirin-3 yl)benzamido)benzamido)pent-4-ynamido)benzamido)benzamido)benzoic acid (**184**) or Cysto-354.

Ester **232** (23 mg, 25 µmol, 1.00 eq.) was dissolved in trifluoroacetic acid (1.6 mL) at 0 °C. After stirring at room temperature for 30 min, the mixture was cooled to 0 °C. Then, Et_2_O (20 mL) was poured into the reaction mixture. The resulting precipitate was filtered and washed with an excess of Et_2_O. Purification by semi-preparative reverse-phase HPLC (operating time 120 min, H_2_O/CH_3_CN = 90/10 → 50/50 in 60 min, then 100% CH_3_CN, t_r_ = 100 min) furnished carboxylic acid **184** (Cysto-354) (9 mg, 11 µmol, 43%) as a grey foam.

**^1^H-NMR (600 MHz, DMSO-*d*_6_):** δ = 12.78 (bs, 1 H, CO_2_H), 12.30 (bs, 1 H, OH), 10.62 (s, 2 H, CONH), 10.58 (s, 1 H, CONH), 9.40 (bs, 1 H, CONH), 8.76 (d, *J* = 7.6 Hz, 1 H, CONH), 8.10 ‑ 8.08 (m, 2 H, Ar‑*H*), 8.01 ‑ 7.94 (m, 8 H, Ar‑*H*), 7.90 ‑ 7.89 (m, 2 H, Ar‑*H*), 7.87 ‑ 7.85 (m, 2 H, Ar‑*H*), 7.83 ‑ 7.81 (m, 2 H, Ar‑*H*), 7.48 ‑ 7.47 (m, 2 H, Ar‑*H*), 4.81 (q, *J* = 7.4 Hz, 1 H, C*H*NH), 4.55 (bs, 1 H, C*H*Me_2_), 2.94 (t, *J* = 2.6 Hz, 1 H, -C≡CH), 2.83 ‑ 2.74 (m, 2 H, CH_2_ *propargylic*), 1.27 (d, *J* = 6.1 Hz, 6 H, CH*Me_2_*) ppm.

**^13^C-NMR (150 MHz,** **DMSO-*d*_6_):** δ = 169.7 (q, C=O), 168.4 (q, C=O), 166.9 (q, C=O), 166.0 (q, C=O), 164.8 (q, C=O), 164.2 (q, C=O), 154.1 (q, COH; not visible in 13C but in HMBC), 142.2 (Ar‑C), 142.0 (Ar‑C), 141.8 (Ar‑C), 136.3 (Ar‑C), 130.8 (Ar‑C), 130.2 (Ar‑C), 128.8 (Ar‑C), 128.8 (Ar‑C), 128.7 (Ar‑C), 128.6 (Ar‑C), 128.4 (Ar‑C), 128.3 (Ar‑C), 127.6 (Ar‑C), 126.6 (Ar‑C), 122.9 (Ar‑C), 121.8 (q, quartet, *J* = 274.8 Hz, CF_3_), 120.7 (Ar‑C), 120.3 (Ar‑C), 119.4 (Ar‑C), 119.4 (Ar‑C), 119.0 (Ar‑C), 112.9 (Ar‑C), 80.6 (q, -*C*≡CH), 73.2 (t, *C*HMe_2_), 69.8 (t, -C≡*C*H), 53.5 (t, CHNH), 28.1 (q, quartet, *J* = 39.7 Hz, *C*‑CF_3_), 22.3 (p, CH*Me_2_*), 21.4 (s, CH_2_ *propargylic*) ppm.

**^19^F-NMR (376 MHz,** **DMSO-*d*_6_):** δ = −64.66 ppm.

**HRMS (ESI):** m/z calculated for C_45_H_35_N_7_O_9_F_3_ [M−H]^−^: 874.2448; found: 874.2458.

${\boldsymbol{[}\boldsymbol{\alpha}\boldsymbol{]}}_{\mathbf{D}}^{\boldsymbol{24}\boldsymbol{.}\boldsymbol{4}}$ = +17.0° (*c*, 0.1, DMSO-*d*_6_).

**R*_ƒ_*** (PE/EtOAc =1/5): 0.13.

**NMR spectra**

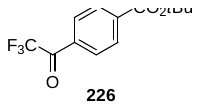

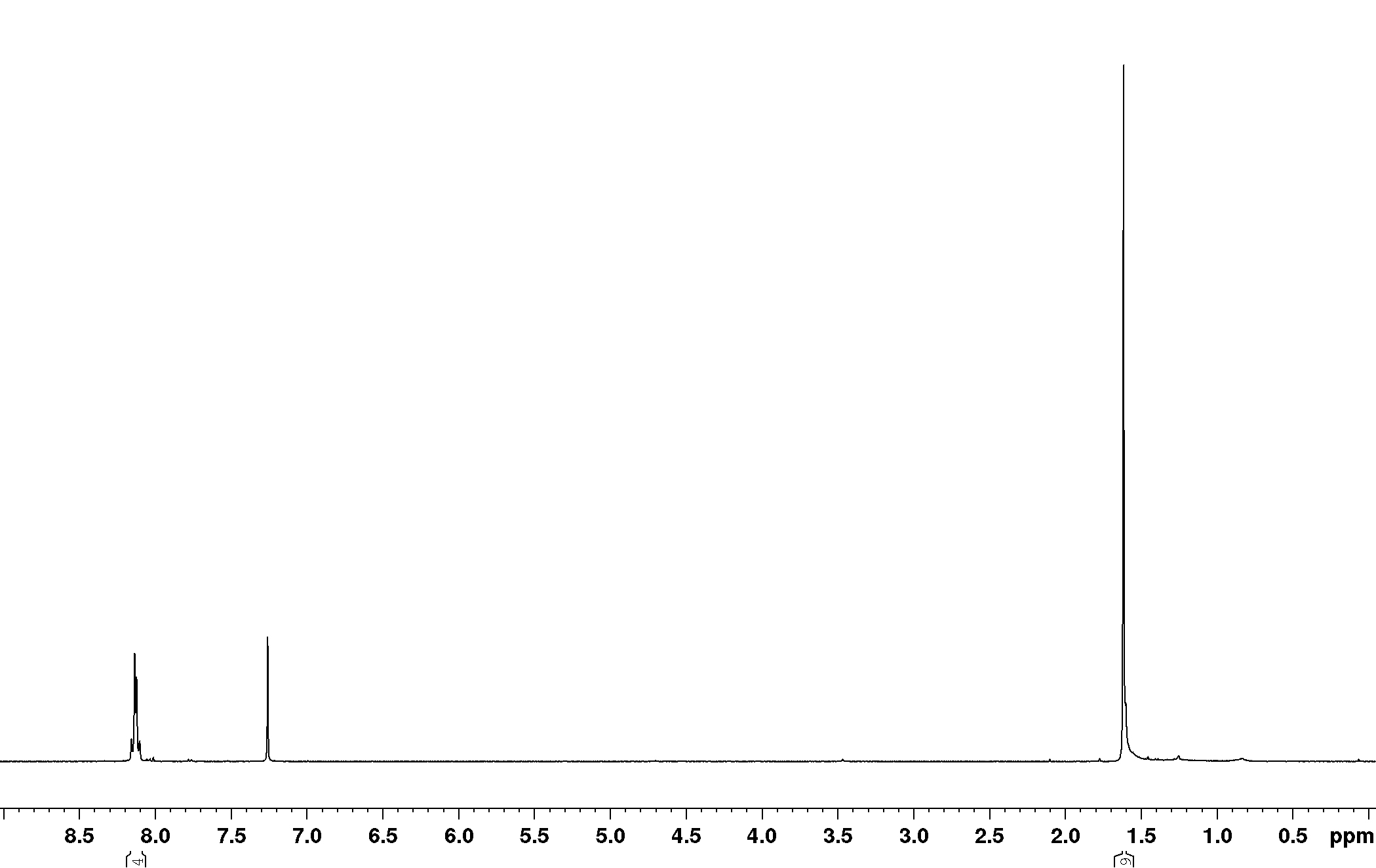

**Supplementary Figure 18.** ^1^H-NMR spectrum of compound **226**.

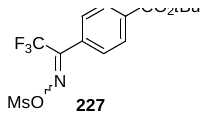

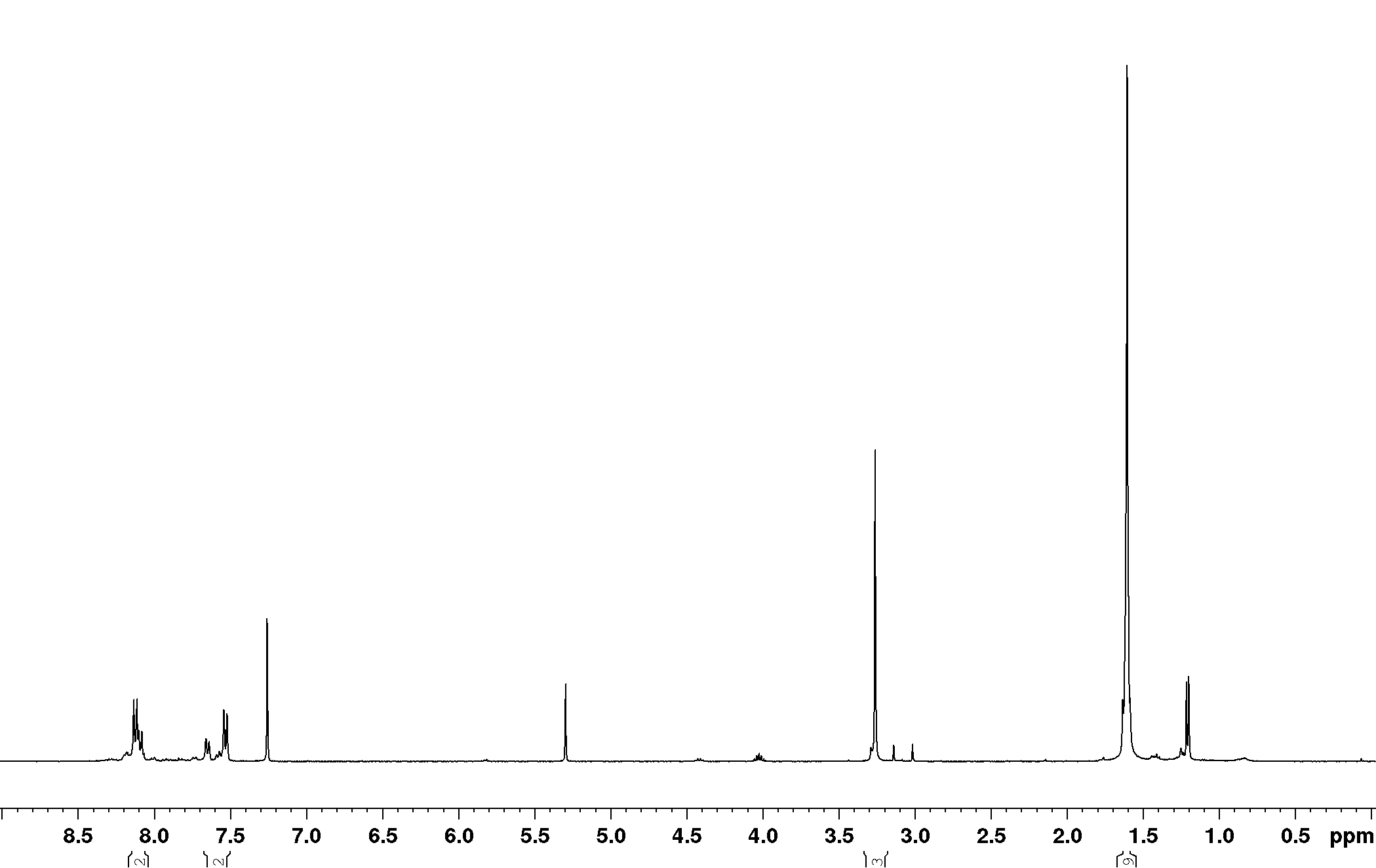

**Supplementary Figure 19.** ^1^H-NMR spectrum of compound **227**.

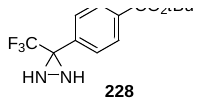

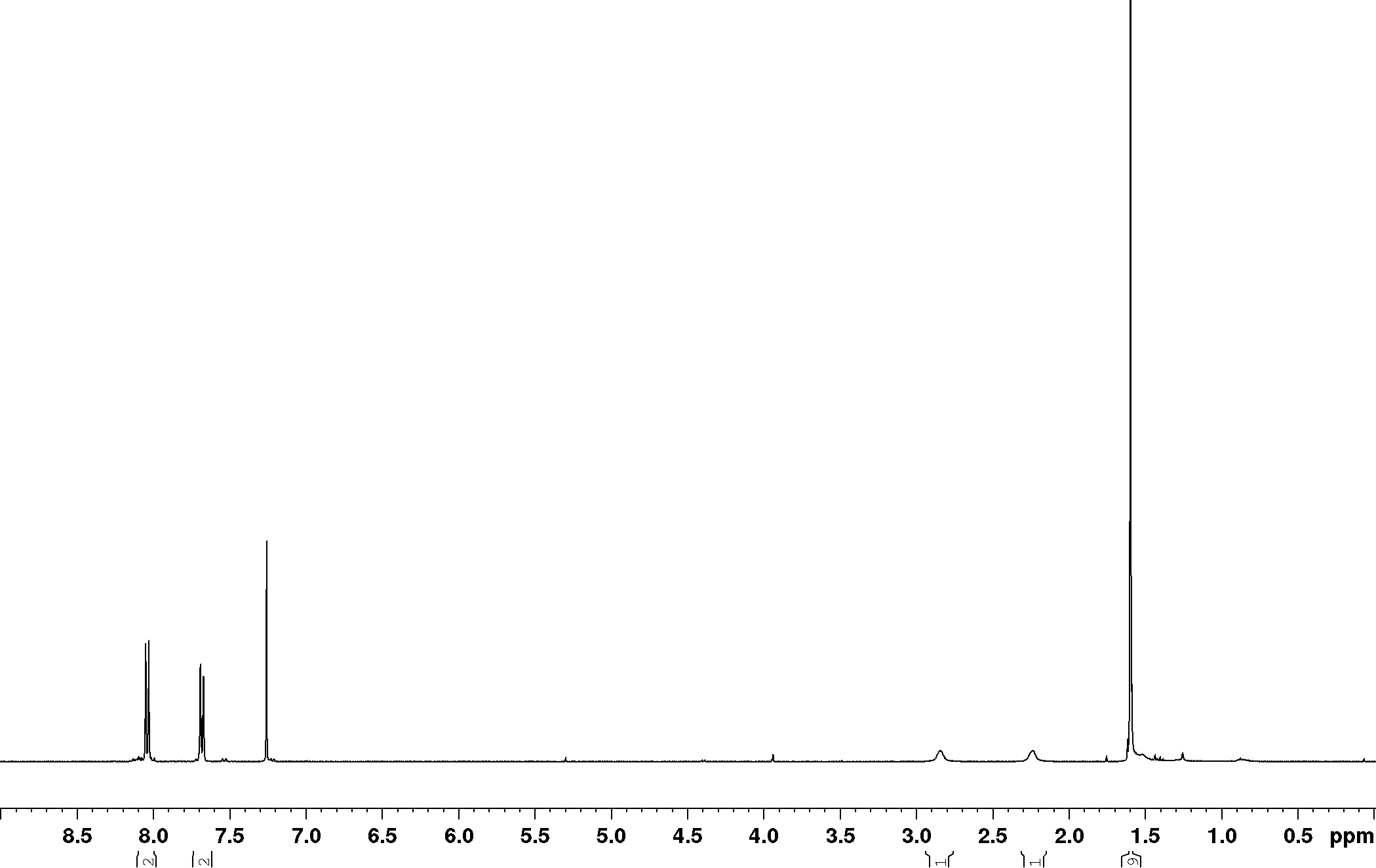

**Supplementary Figure 20.** ^1^H-NMR spectrum of compound **228**.

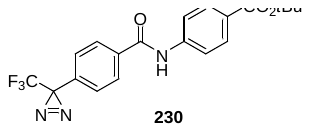

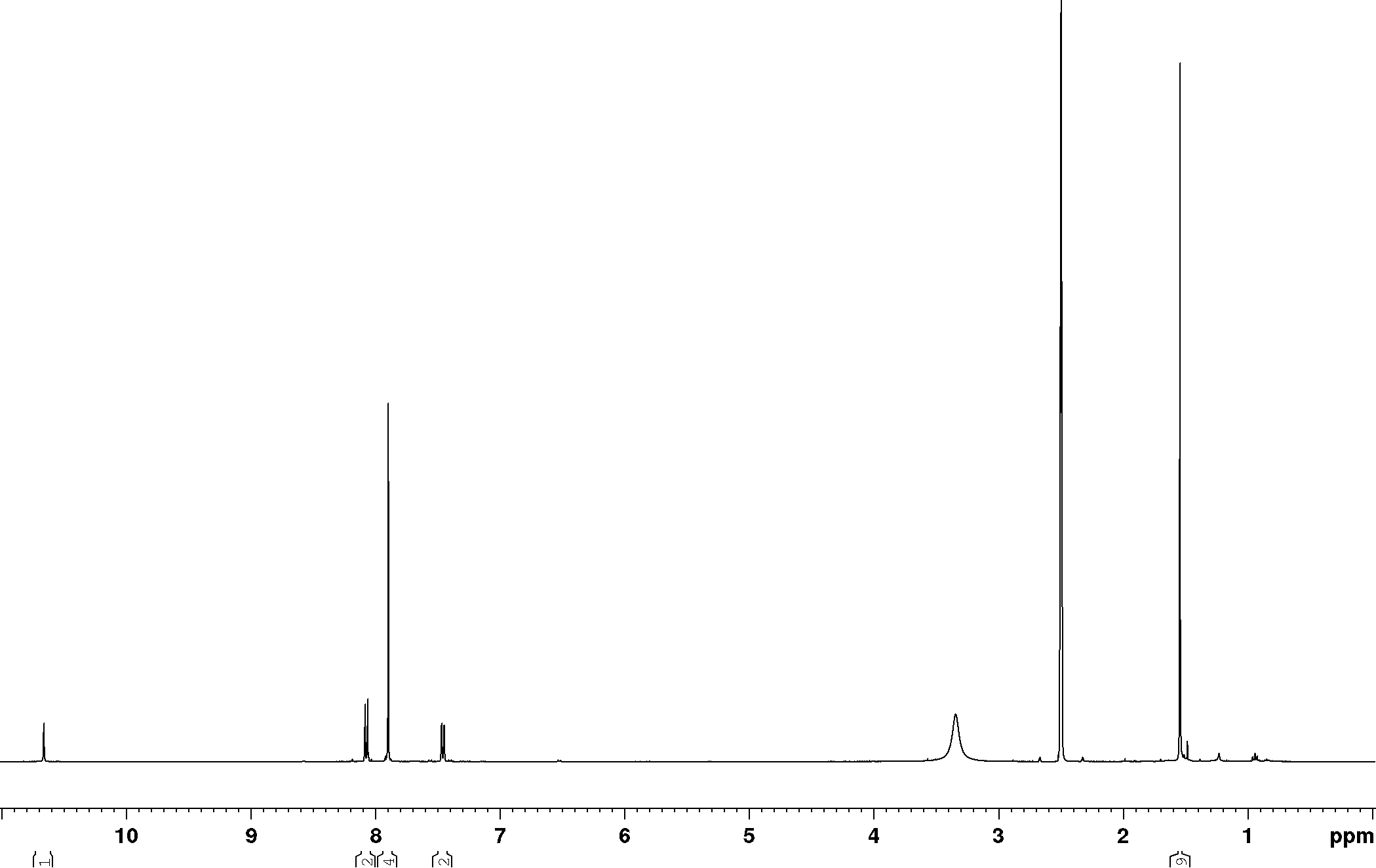

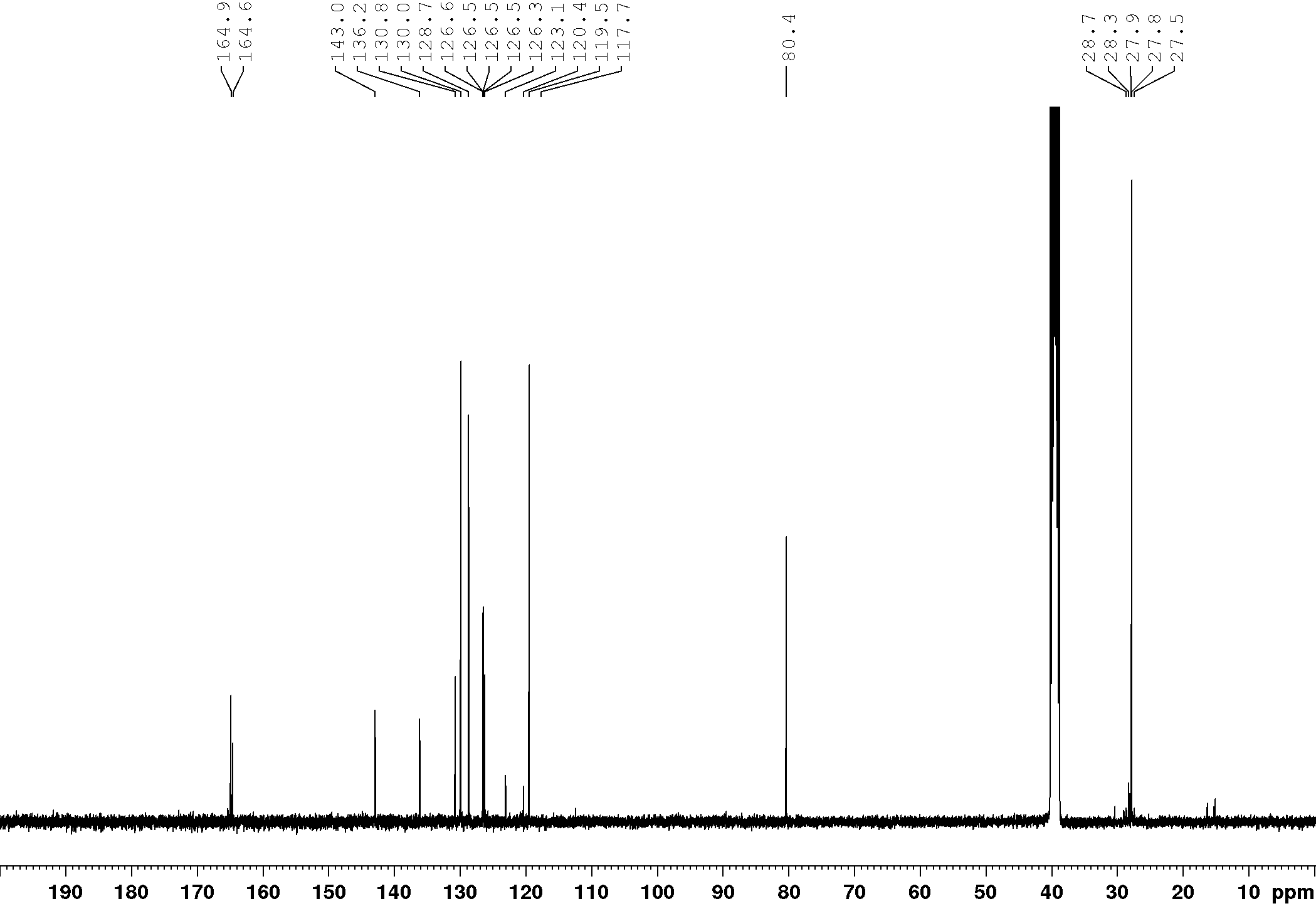

**Supplementary Figure 21.** ^1^H- and ^13^C-NMR spectra of compound **230**.

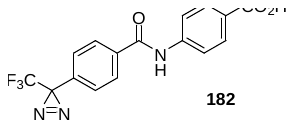

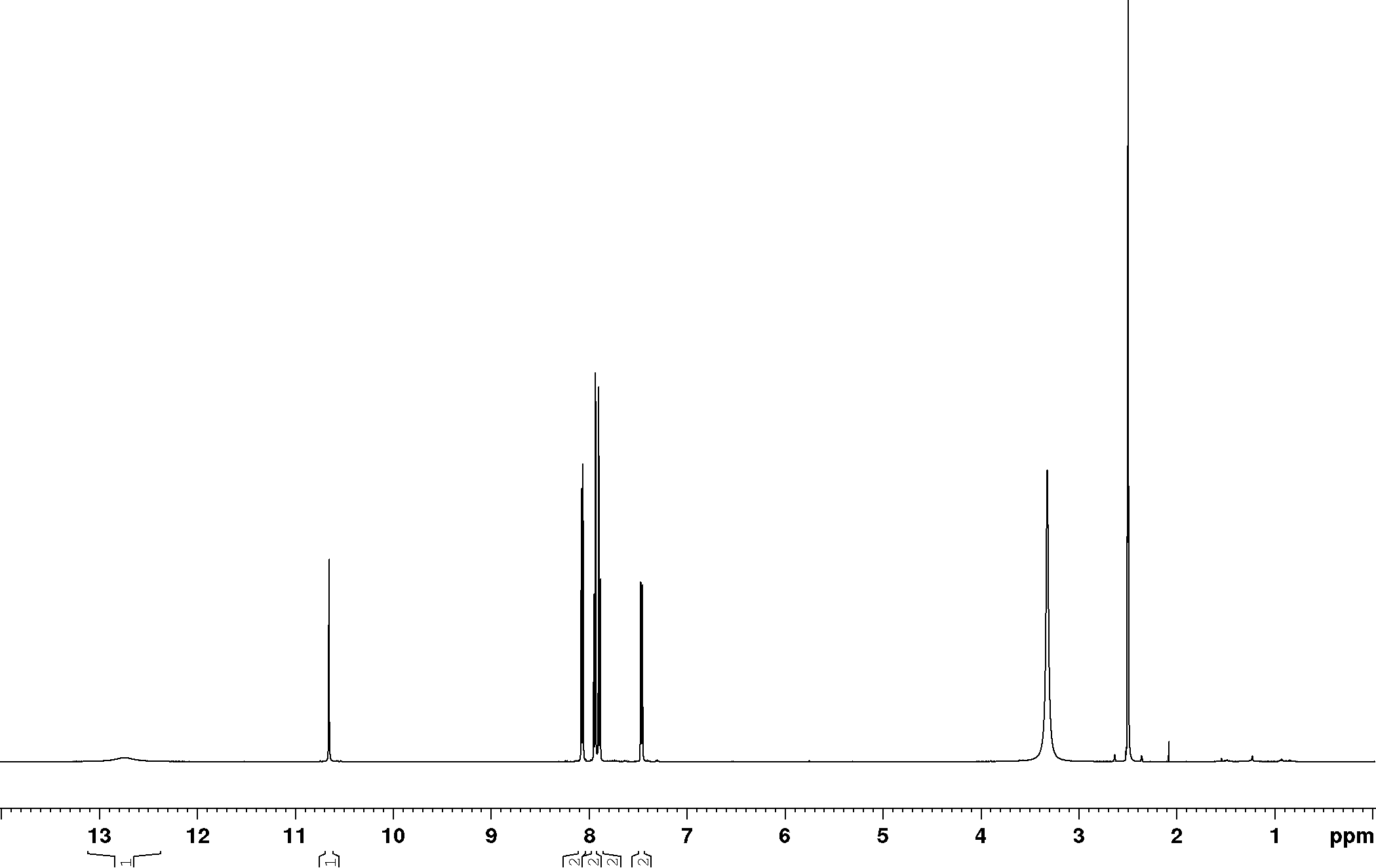

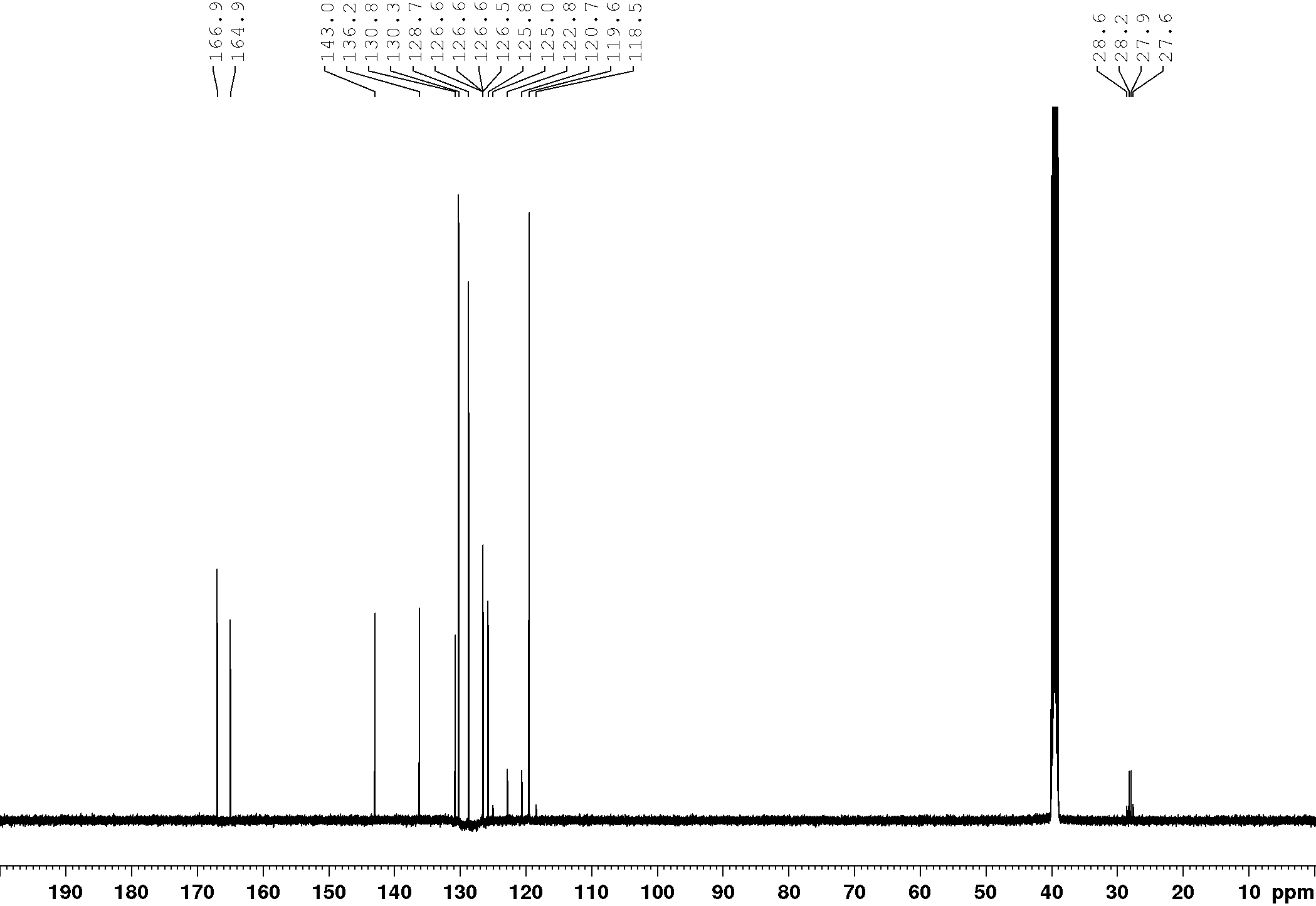

**Supplementary Figure 22.** ^1^H- and ^13^C-NMR spectra of compound **182**.

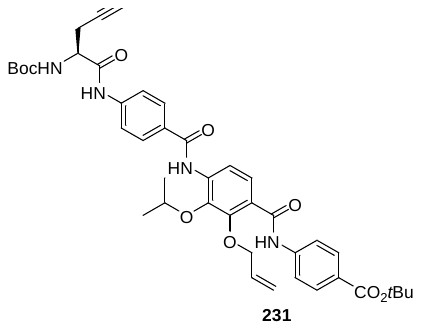

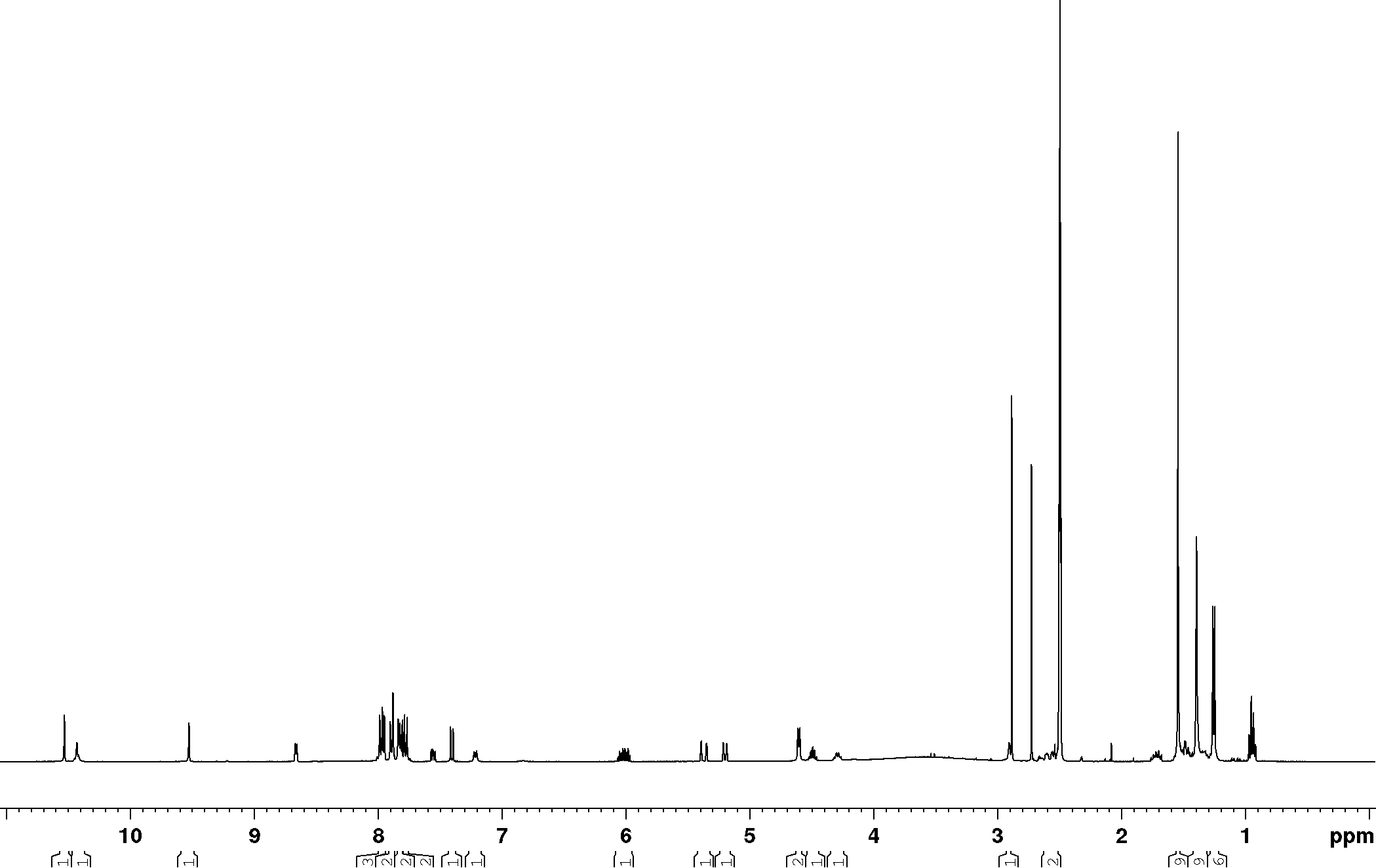

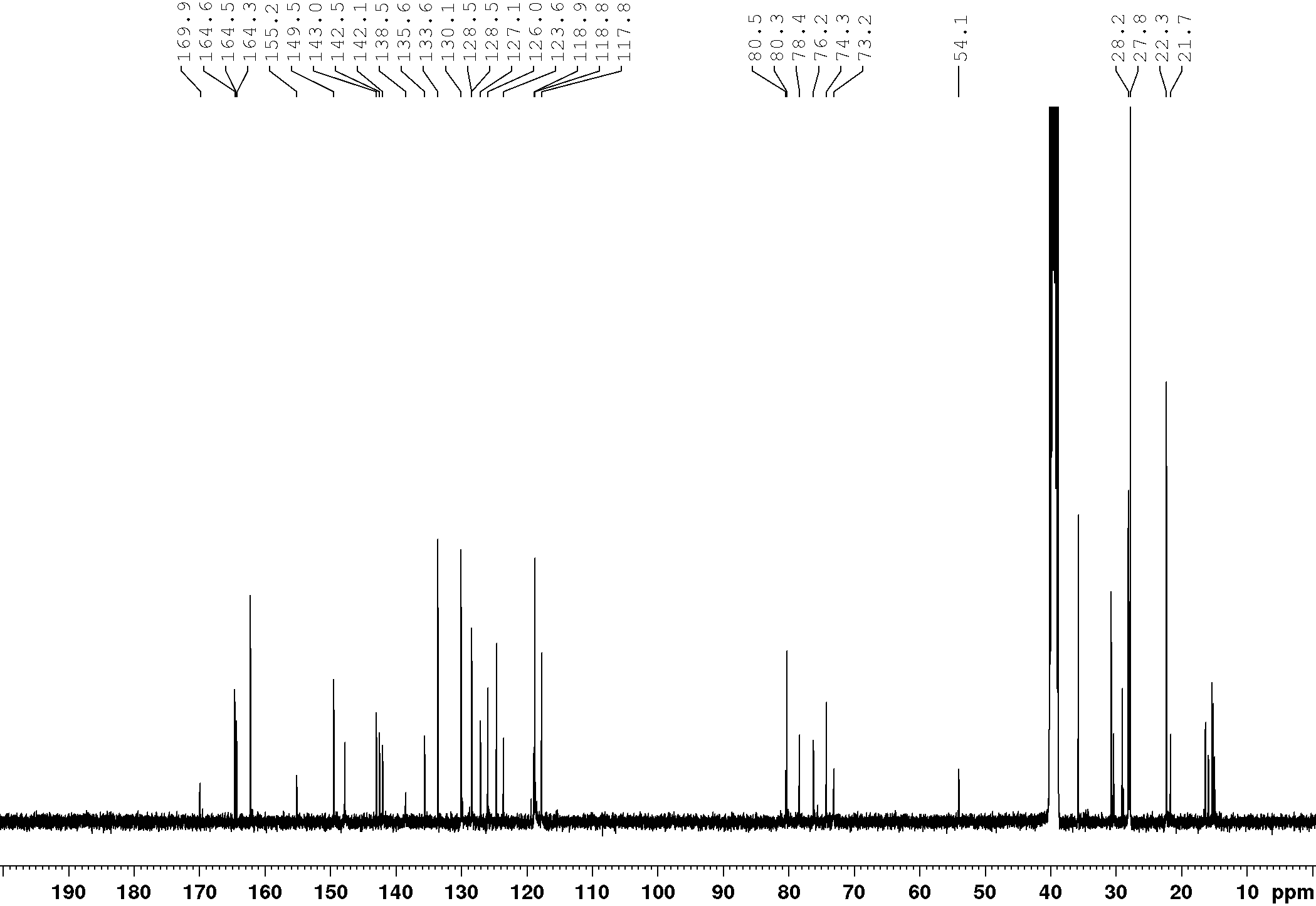

**Supplementary Figure 23.** ^1^H- and ^13^C-NMR spectra of compound **231**.

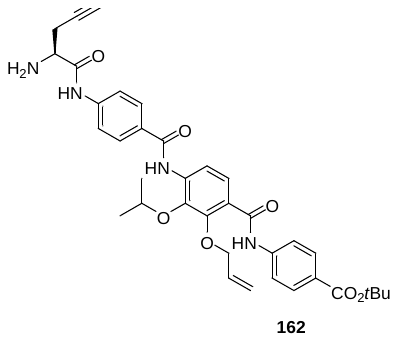

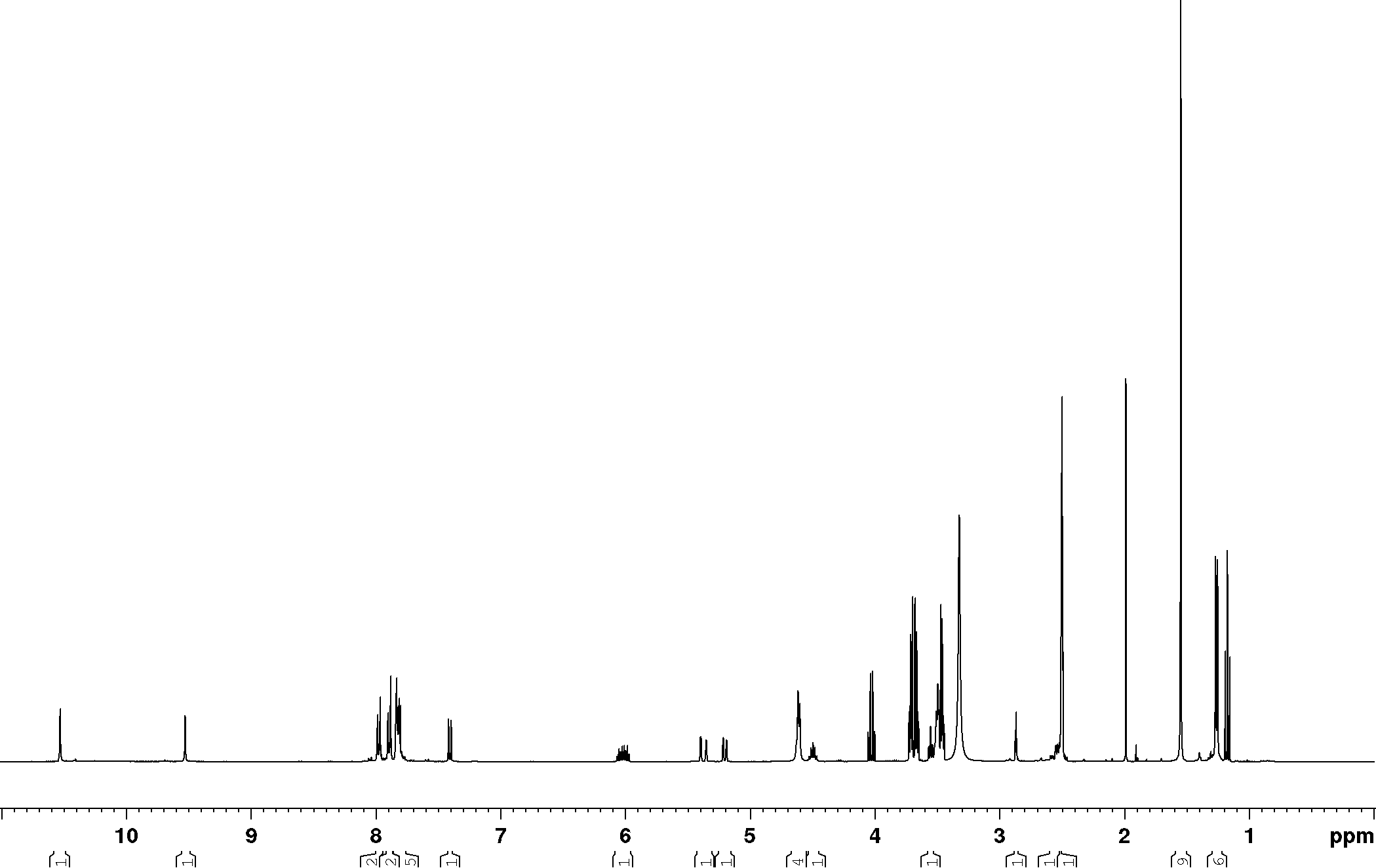

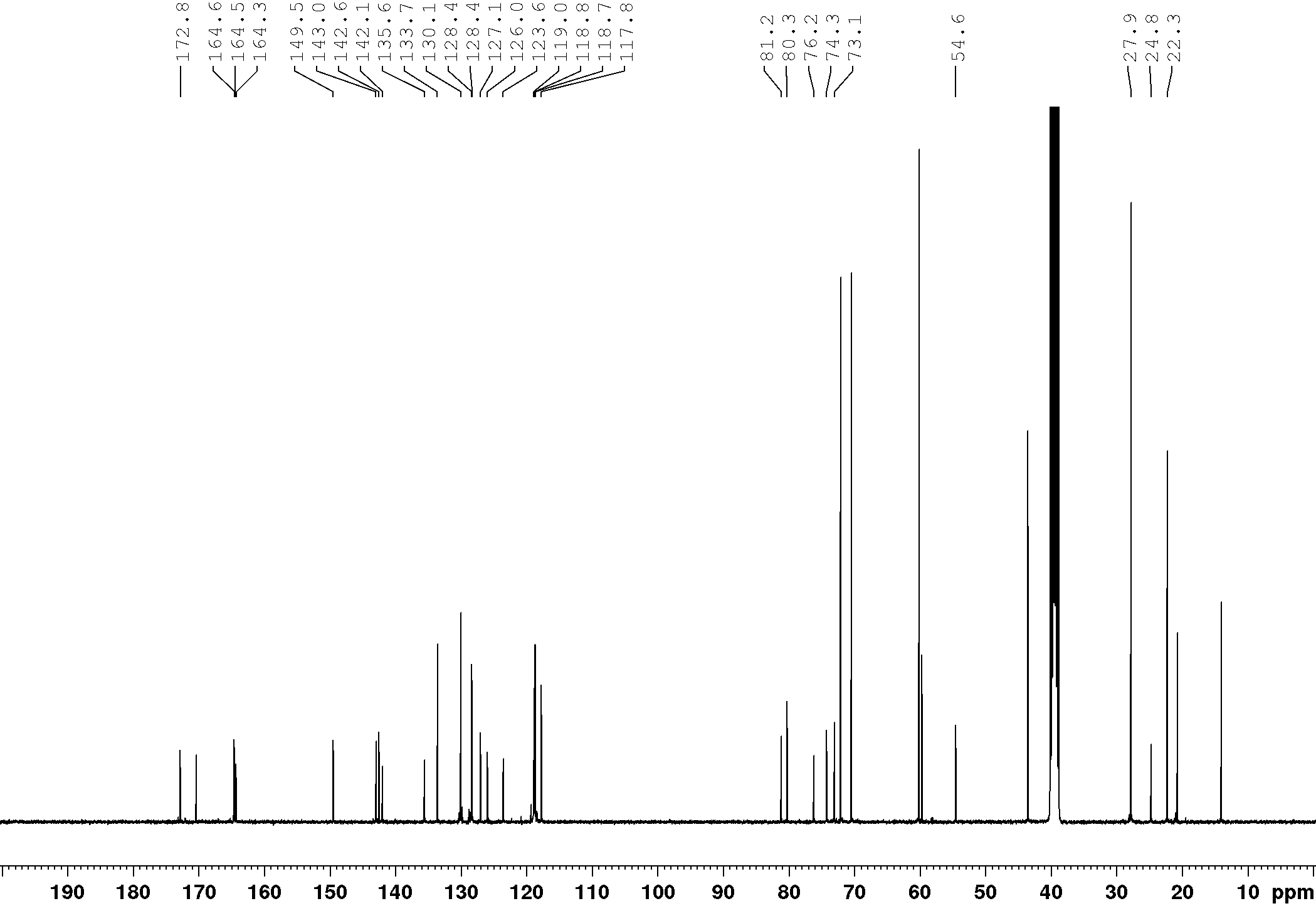

**Supplementary Figure 24.** ^1^H- and ^13^C-NMR spectra of compound **162**.

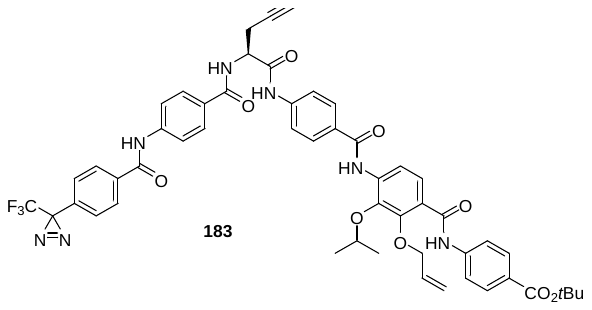

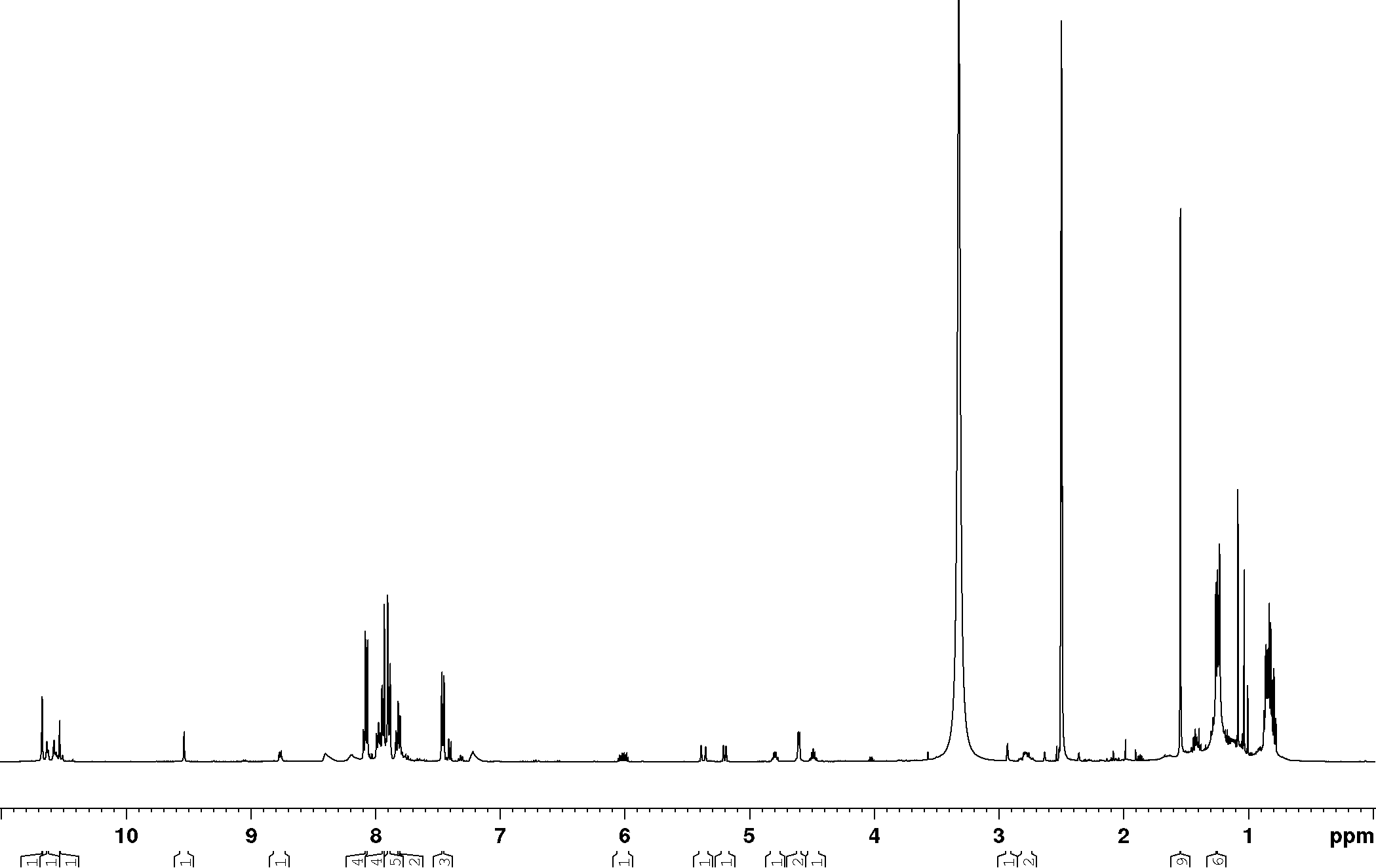

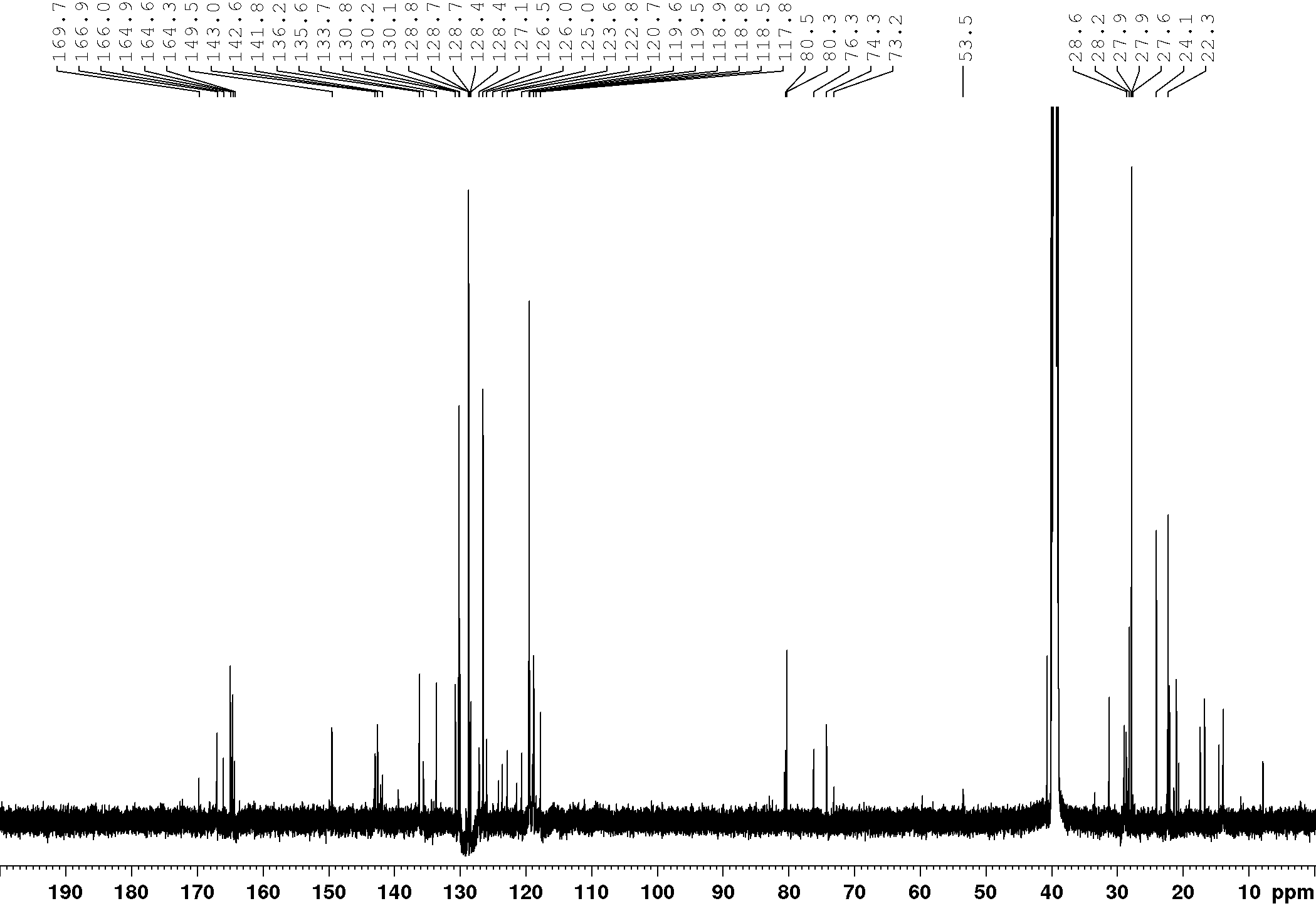

**Supplementary Figure 25.** ^1^H- and ^13^C-NMR spectra of compound **183**.

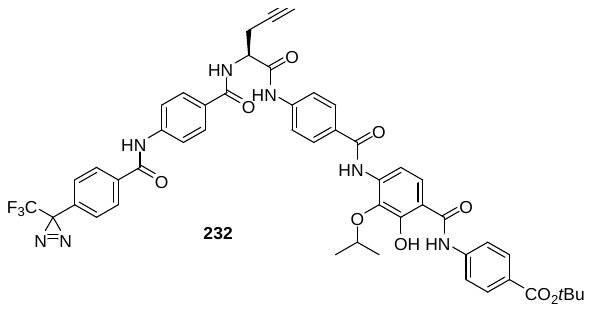

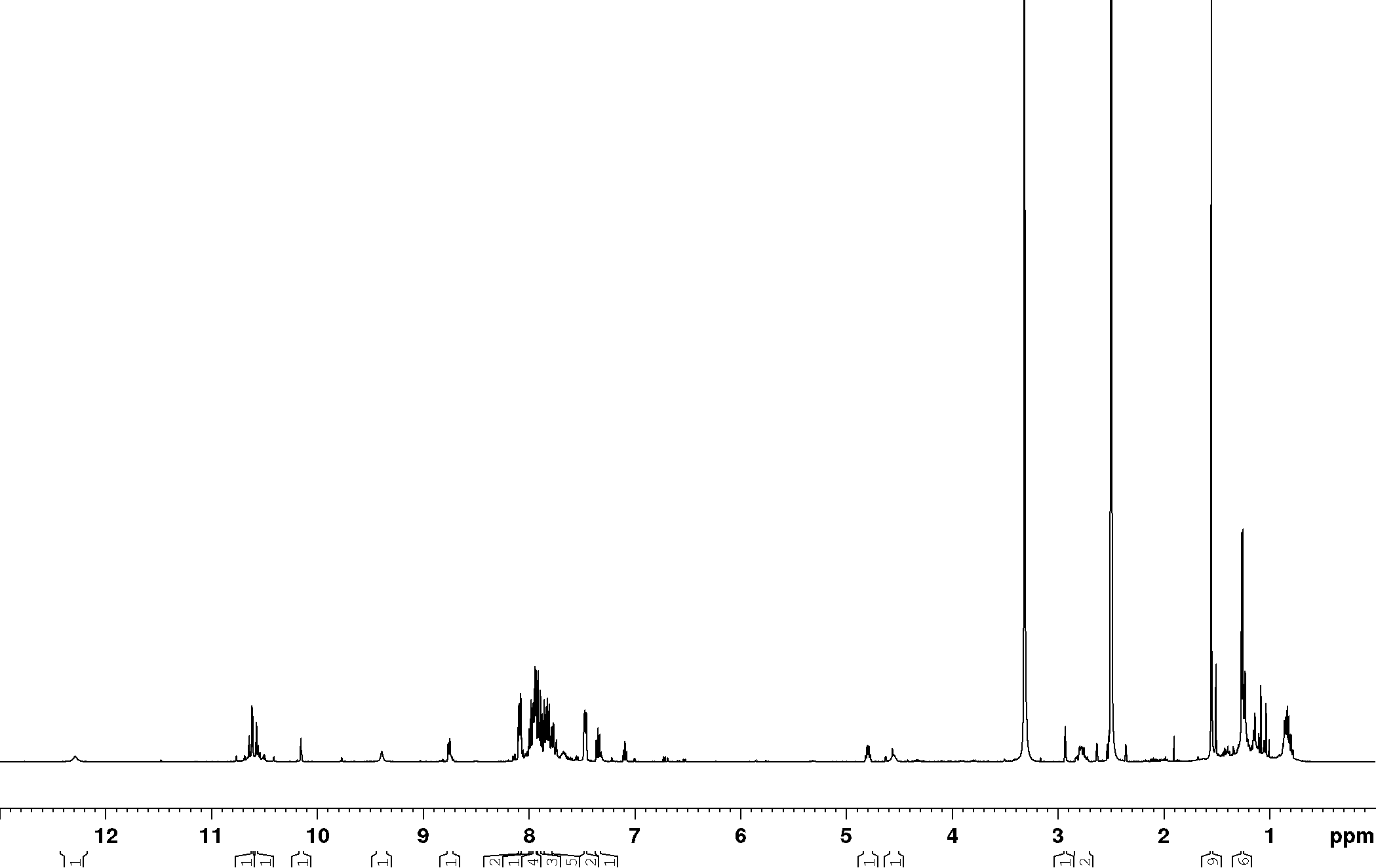

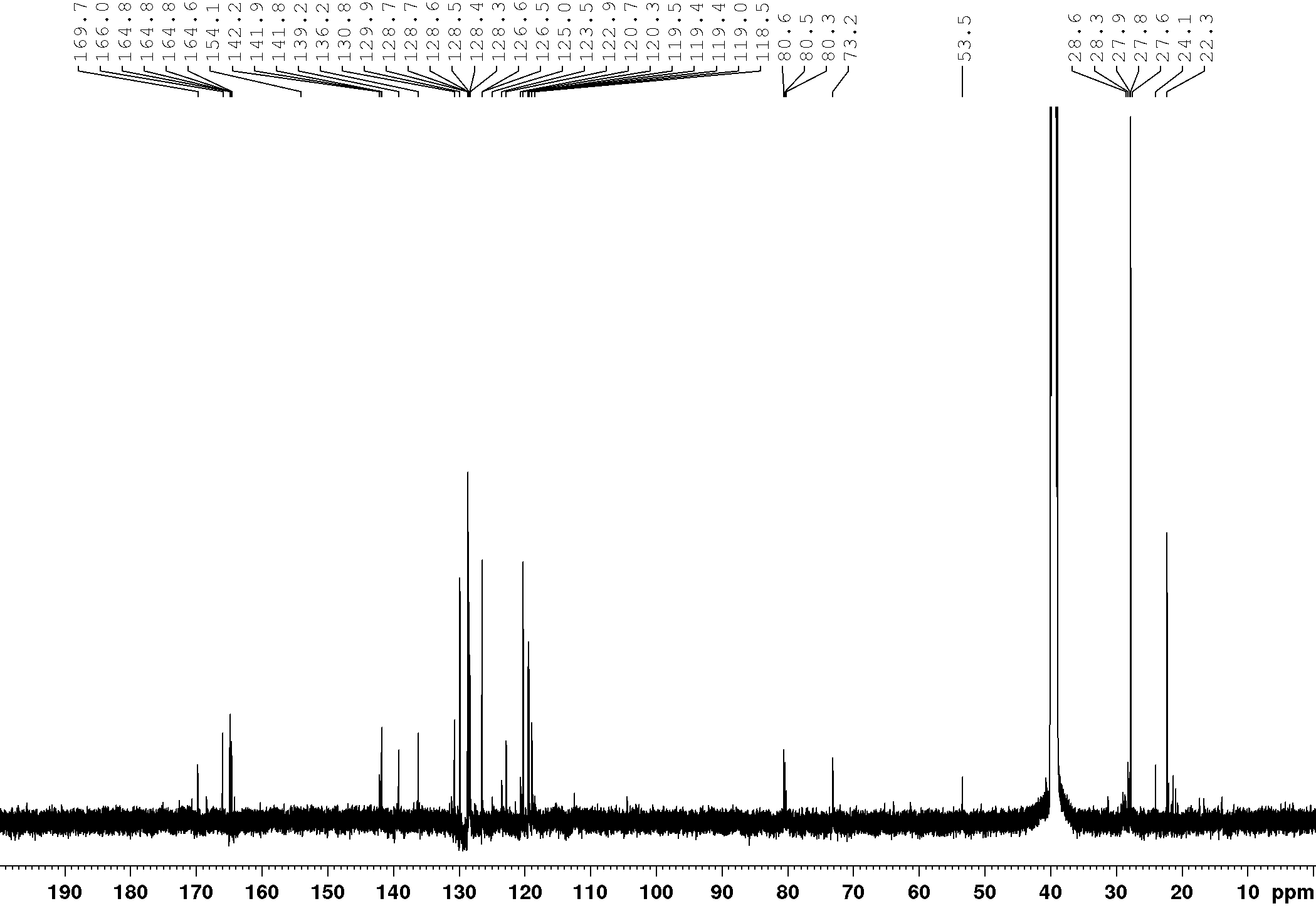

**Supplementary Figure 26.** ^1^H- and ^13^C-NMR spectra of compound **232**.

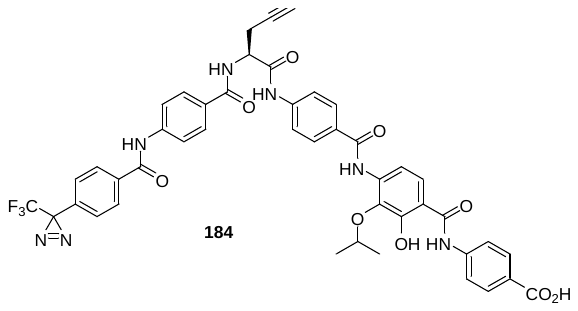

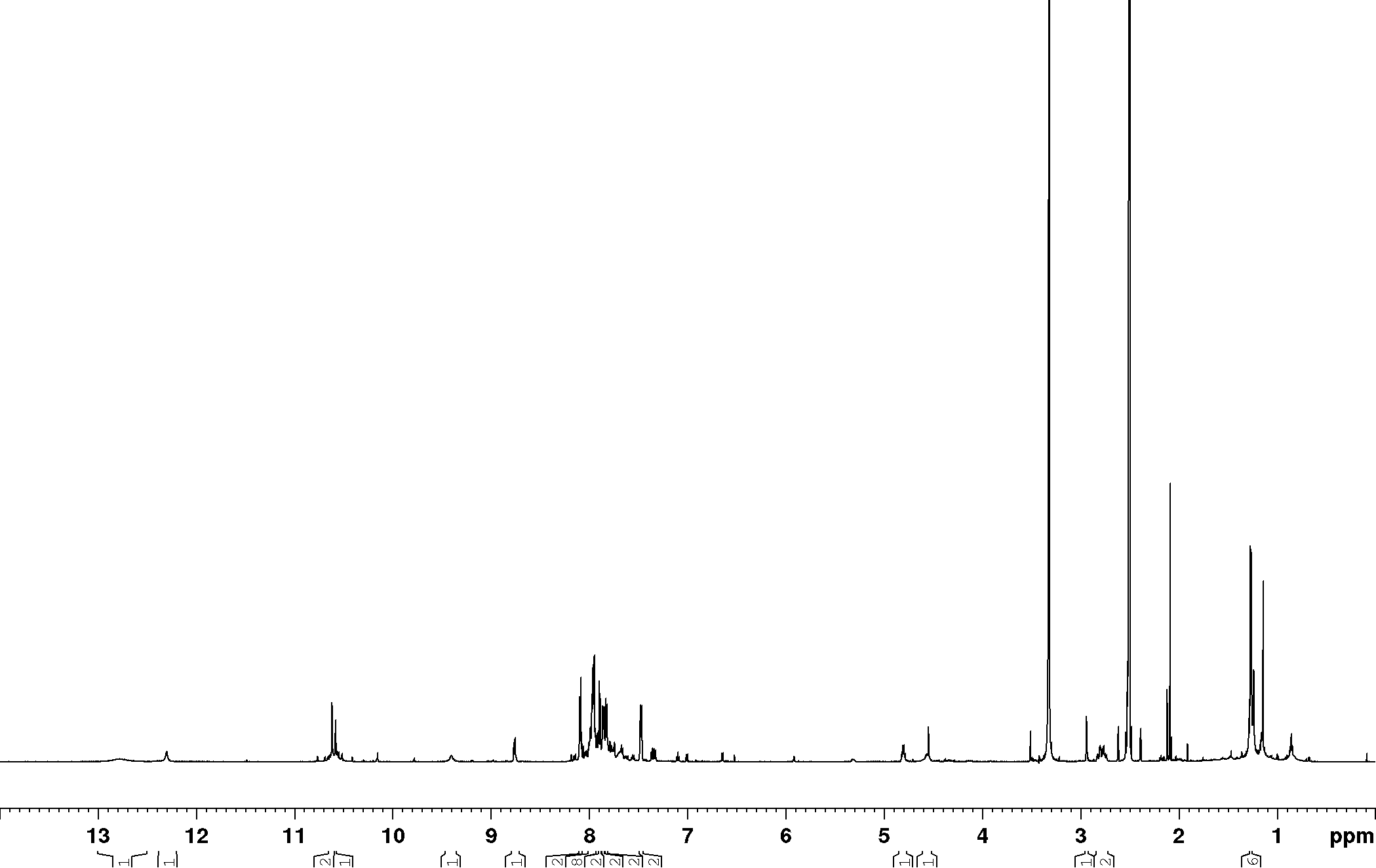

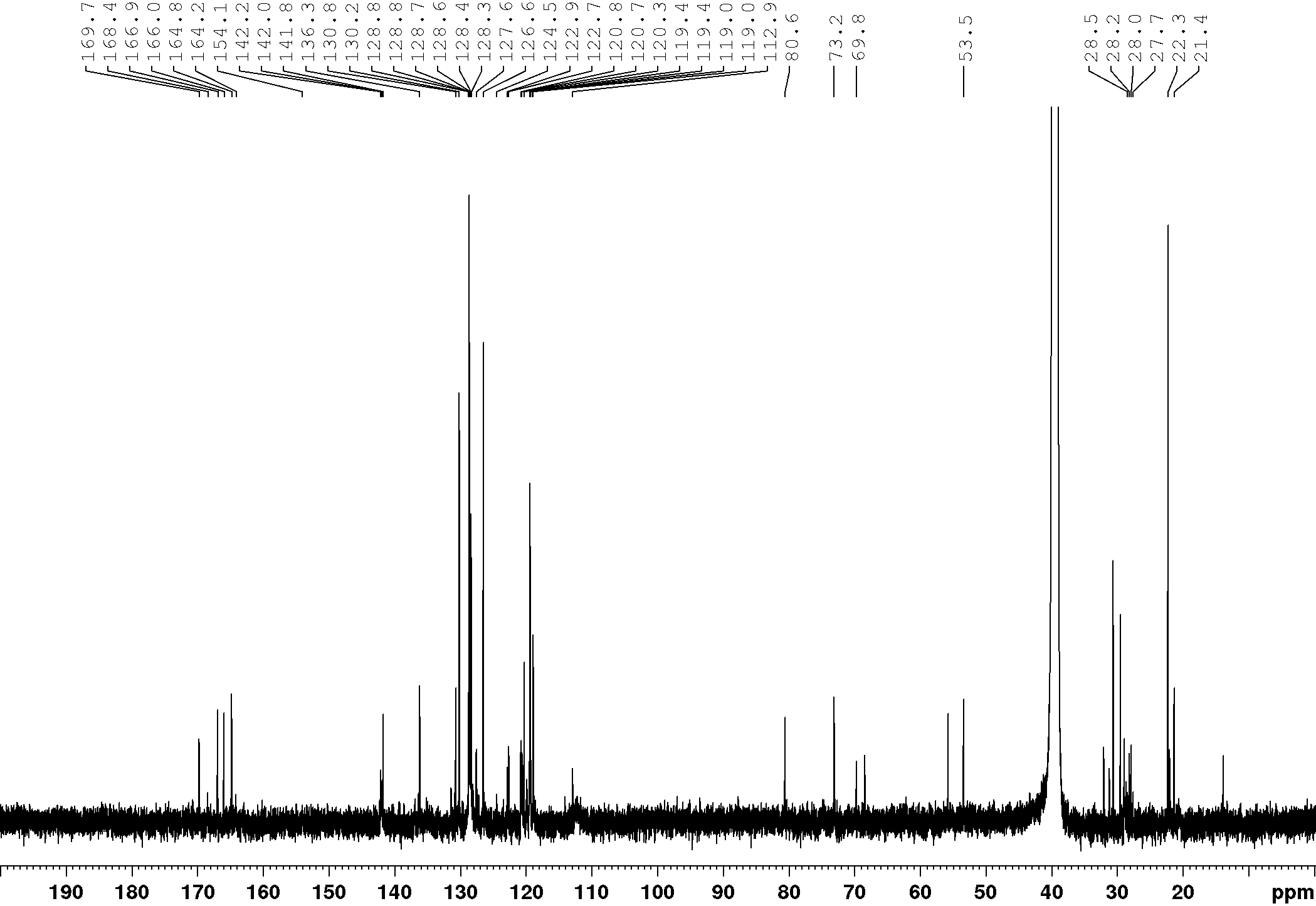

**Supplementary Figure 27.** ^1^H- and ^13^C-NMR spectra of compound **184** or Cysto-354.

Cytotoxic evaluation of Cysto-354

MTT assay was performed as described in the methods section. Cysto-354 showed slightly higher cytotoxic potential on tested cell lines than non-functionalized CYS derivatives (see also FIG 1C).

**Supplementary Figure 28.** Cysto-354 showed reduction of cell viability at the highest tested concentration (37 µM). (*n* = 3)

No enrichment of human topoisomerases could be observed

Topoisomerase enrichment after Cysto-354 treatment was checked, since CYS inhibit bacterial type II DNA topoisomerases and human topoisomerase IIα (TOP2A) *in vitro*. No significant enrichment of TOP1, TOP2A nor TOP2B could be observed, indicating whole cell selectivity of CYS to bacterial topoisomerases.

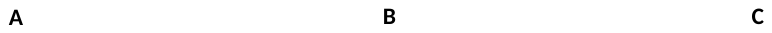

**Supplementary Figure 29 (Related to Figure 5).** Human topoisomerase (TOP1, TOP2A and TOP2B) affinity enrichment. **(A)** AfBPP in HEK293 did not show significant enrichment of human topoisomerases. **(B)** AfBPP in HeLa-CCL-2 did not show significant enrichment of human topoisomerases. **(C)** AfBPP in HepG2 did not show significant enrichment of human topoisomerases. (*n* = 4)

VSVpp entry assay supported specificity of CYS for SCARB1 related HCVpp cell entry inhibition

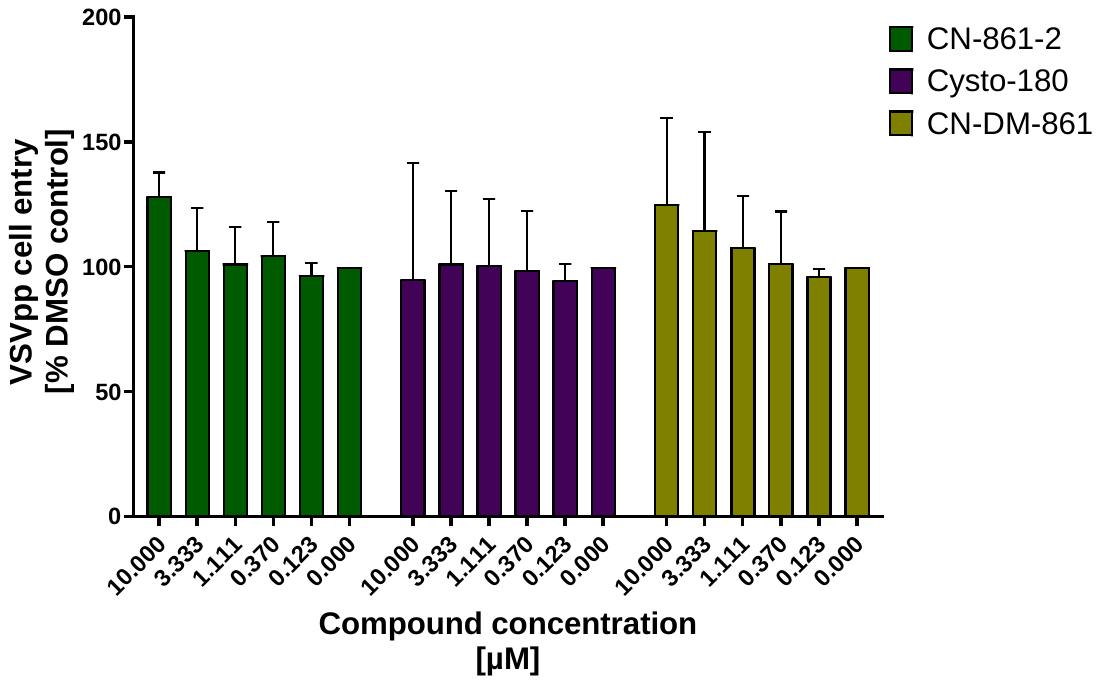

**Supplementary Figure 30 (Related to Figure 6).** Lentivirus-pseudoparticle (pp) cell entry assay

CYS treatment did not demonstrate inhibitory activity on VSVpp cell entry. (*n* = 3)

Molecular docking of CYS to SCARB1

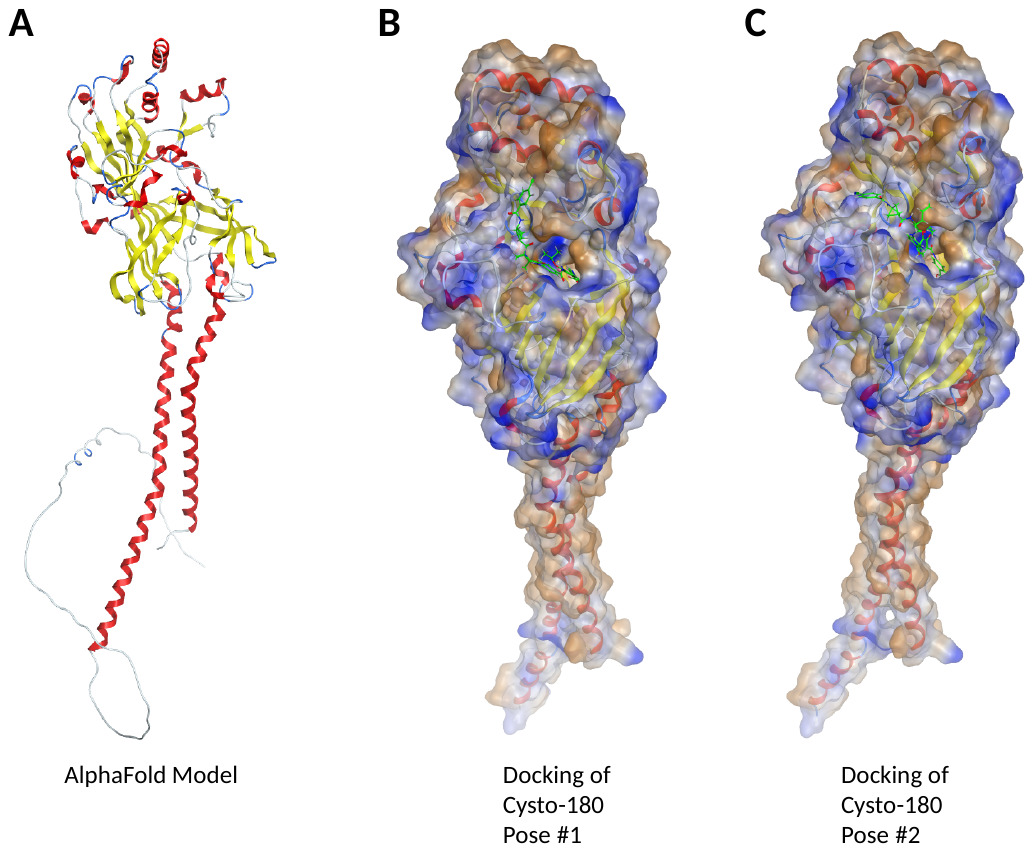

**Supplementary Figure 31.** Molecular docking of CYS to the AlphaFold model of SCARB1. **(A)** AlphaFold model of SCARB1. **(B)** First pose of Cysto-180 molecular docking to SCARB1 model indicates a potential binding pocket in the head group of the protein. **(C)** Second pose of Cysto-180 shows that multiple poses of CYS inside the pocket seem to be possible.

**Supporting References**

1.	Hüttel, S. *et al.* Discovery and Total Synthesis of Natural Cystobactamid Derivatives with Superior Activity against Gram-Negative Pathogens. *Angew. Chemie - Int. Ed.* **56**, 12760–12764 (2017).

2.	Testolin, G. *et al.* Synthetic studies of cystobactamids as antibiotics and bacterial imaging carriers lead to compounds with high: In vivo efficacy. *Chem. Sci.* **11**, 1316–1334 (2020).

3.	Risch, T., Kolling, D., Mostert, D., Herrmann, J. & Müller, R. Point mutations in the ygiV promoter region lead to cystobactamid resistance and reduced virulence in. *ResearchSquare* (2024) doi:https://doi.org/10.21203/rs.3.rs-3751821/v1

4.	Sakurai, K., Yasui, T. & Mizuno, S. Comparative Analysis of the Reactivity of Diazirine-Based Photoaffinity Probes toward a Carbohydrate-Binding Protein. *Asian J. Org. Chem.* **4**, 724–728 (2015).

5.	Kohnhäuser, D. Synthetic optimization of cystobactamid analogs as antibiotics. (2021). doi:https://doi.org/10.15488/11137.
